# Investigating the impact of adverse life events on neuroanatomy across adolescence

**DOI:** 10.64898/2026.05.06.723376

**Authors:** Lisa Wiersch, Katharina Brosch, Erynn Christensen, Elvisha Dhamala

**Affiliations:** Northwell, New Hyde Park, NY, USA; Institute of Behavioral Science, Feinstein Institutes for Medical Research, Manhasset, USA; Department of Psychiatry, Beth Israel Deaconess Medical Center, Harvard Medical School, MA, USA; Department for Psychiatry, Donald and Barbara Zucker School of Medicine at Hofstra/Northwell, Uniondale, USA

**Keywords:** Adverse life events, adolescence, neuroanatomy, machine-learning, brain-based predictive modeling

## Abstract

**Background:** Adversity during adolescence can increase vulnerability to neuropsychiatric disorders. However, how various factors of adversity and stress can shape these effects, and whether these effects vary by sex, developmental stage, or reporting source have not been established.

**Methods:** Here, we used multivariate brain-based predictive models to examine how the number, appraisal (positive or negative), and impact of adverse life events during adolescence (reported either by the youth or their caregivers) are reflected in neuroanatomy (cortical thickness, surface area, cortical and subcortical grey matter volume). We used data from the Adolescent Brain Cognitive Development (ABCD) study at 2-year (*N=*6,366, age 11-12), 4-year (*N*=5,035, age 13-14), and 6-year (*N*=5,953, age 15-16) follow-ups to examine neural correlates of adversity.

**Results:** Descriptively, youth, compared to their caregivers, reported more adverse life events, higher impact, and were more likely to appraise events negatively. Multivariate associations between adverse life events and neuroanatomy were sparse at the 2-year and 4-year follow-ups but emerged at the age of 15-16 (6-year follow-up). Among the significant associations, positively appraised events were predominantly associated with higher measures of cortical thickness, surface area, and volumes, whereas negatively appraised events mapped predominantly negatively on neuroanatomy.

**Conclusions:** Our analyses suggest that associations between childhood adversity and the brain vary across adolescent development. Furthermore, our results illustrate important differences between youth and caregiver reports, highlighting the importance of differentiating between reporting sources when interpreting the impact of adverse life events on neurodevelopment.

---

Adolescence is characterized by high neural plasticity, reflecting a critical window of vulnerability (Bethlehem et al., 2022; Gabard-Durnam & McLaughlin, 2020; Lockhart et al., 2018). Adversity during adolescence can increase the risk of developing a neuropsychiatric disorder (Dunn, 2018; Hardi, Beltz, et al., 2024; Hardi, Peckins, et al., 2024; Lockhart et al., 2018; Sisk & Gee, 2022). Moreover, exposure to adverse or stressful life events during youth can shape structural and functional brain organization (Gee, 2021, 2022; Teicher et al., 2003, 2016), involving regions such as the medial prefrontal cortex, amygdala, and hippocampus, among others (Gee, 2022; Mclaughlin et al., 2019). Alterations in brain organization can also extend to large-scale brain networks that mediate emotion regulation, reward processing, social information processing, and associative learning (Gee, 2021).

There is a growing body of research studying the associations between various forms of adversity, brain organization, and psychopathology, leveraging multivariate approaches and publicly available datasets with large sample sizes (e.g., the Adolescent Brain Cognitive Development (ABCD) Study). Alterations in cortical thickness and volume in key regions are associated with attention, executive functioning, and emotion regulation following trauma exposure (Jeong et al., 2021). Vulnerability in response to adversity is evident in subcortical structures, with the putamen connecting environmental stress to psychotic symptoms (Ma et al., 2025). Adversity is also related to dynamic alterations in functional connectivity (Elton et al., 2025; Vedechkina et al., 2024). Importantly, these effects are not necessarily uniformly negative: alterations in cortico-limbic resting-state functional connectivity in response to negative life events may represent accelerated neurobiological maturation, serving as a potentially short-term protective mechanism (Brieant et al., 2021; Petrican et al., 2021; Teicher et al., 2016). Further, the impact on brain maturation may depend on the specific dimensions of early life adversity.

Caregiver lack of support and supervision has been associated with younger-looking brains, while caregiver psychopathology, socioeconomic disadvantage, or trauma exposure are linked to older-looking brains, as revealed by brain age gap analyses (Beck et al., 2025). It is therefore important to characterize how adversity-brain relationships may vary across adversity dimensions, neural features, and developmental stages.

Moreover, understanding whole-brain neural signatures also requires an examination of subjective appraisal (i.e., whether an event is appraised as positive or negative) and its impact. The same life event (e.g., relocation or parental divorce) may be appraised differently among youth, and this appraisal may be uniquely linked to neurobiology. The total number of adverse life events, along with their cumulative impact, i.e., how much it affected a person’s life, regardless of valence, is significantly associated with left medial orbitofrontal cortex volumes in adults, driven by both positive and negative event appraisals (Ringwald et al., 2021). Additionally, previous research has shown inconsistent findings regarding the discrepancy and agreement between children and their caregivers in reporting life events (Allen et al., 2012; Johnston et al., 2003; Tingskull et al., 2015). Therefore, the assessment of adverse life events, their appraisal, and impact, especially in developmental contexts, may differ depending on whether they are assessed by youth or their caregivers.

A critical gap in research is how the neural signatures of stress and adversity vary across neuroanatomical features. Studies have predominantly focused on a single feature (e.g., thickness), but the effects are not necessarily uniform. Cortical thickness is the distance between the grey-white matter junction and the pial surface (Fehlbaum et al., 2022). It rapidly increases during prenatal and neonatal development, peaks in early childhood, and subsequently declines throughout adolescence and adulthood (Bethlehem et al., 2022). Surface area is the area of the cortical outer layer (Fehlbaum et al., 2022). It continuously increases throughout childhood, reaching its peak in adolescence, and then gradually declines (Bethlehem et al., 2022). Grey matter volume is the total amount of neuronal tissue. Cortical grey matter volumes are estimated as the product of cortical thickness and surface area (Fehlbaum et al., 2022; Vijayakumar et al., 2016), whereas subcortical grey matter volumes refer to grey matter nuclei in the subcortex. Cortical volume increases throughout early childhood and peaks between early and late childhood, followed by a slow and steady decrease until late adulthood. In contrast, subcortical volumes show a slower, more prolonged increase, a later peak in adolescence, and a slower decline (Bethlehem et al., 2022). Considering the inherent differences and distinct developmental trajectories (Bethlehem et al., 2022) of the individual measures, it is crucial to analyze the neural signatures of stress across multiple neuroanatomical features throughout development.

Timing and sex are two particularly important determinants that may shape the effects of stress or adversity (Bath, 2020; Ho et al., 2024; Michael et al., 2025). These effects may be heightened if adverse life events occur during sensitive developmental periods (Eiland & Romeo, 2013; Gee & Casey, 2015). Sex can also influence the processing of stress induced by adverse life events, the trajectories of neurodevelopment, and neuroendocrine responses (Bangasser & Valentino, 2014; Bangasser & Wicks, 2017; Hodes et al., 2023). Regarding the effects of developmental timing, female adolescents reach peak grey matter volumes one to two years earlier than male adolescents and have larger subcortical volumes after puberty; in contrast, male adolescents show larger cortical surface area until the age of 15 (Gur & Gur, 2016; Koolschijn & Crone, 2013; Lenroot et al., 2007). Regarding the effects of timing of the adverse life event, some events may be more likely to occur or to be recognized among older than younger adolescents (e.g., changes in the parental financial situation). Thus, it is important to consider both timing and sex when characterizing the neural signatures of stress.

Here, we investigate the associations between adverse life events and brain structure across three time points during adolescence using brain-based predictive models leveraging data from the Adolescent Brain Cognitive Development (ABCD) dataset. We use machine learning analyses, which provide a data-driven framework to identify neural signatures distributed across multiple brain regions. Specifically, we examine how the number, appraisal, and impact of adverse life events relate to different neuroanatomical measures (cortical thickness, surface area, cortical and subcortical grey matter volumes) at the 2-year (ages 11-12), 4-year (ages 13-14), and 6-year (ages 15-16) follow-ups. To capture potential sex-specific effects, we also examine these relationships separately for female and male adolescents. Furthermore, we compute separate models for youth- and caregiver-reported adverse life events to account for potential discrepancies between self-report and external observations. Following an exploratory approach, we examine multivariate associations between adverse life events and neuroanatomy, including their spatial and temporal patterns.

## Methods

### Dataset

The ABCD study is a prospective longitudinal study assessing children starting at ages 9-10 and following participants for 10 years (Casey et al., 2018). The study recruited participants from 21 research sites across the United States, comprising a cohort of 11,860 participants at baseline. Participants complete a comprehensive battery of neuroimaging, behavioral, developmental, and psychiatric assessments. Here, we used neuroimaging data and the Adverse Life Events Scale of the PhenX Toolkit from the 2-year (ages 11-12), 4-year (ages 13-14), and 6-year (ages 15-16) follow-up time points from the ABCD 7.0 release. We included subjects with complete and quality-controlled neuroimaging data, as well as complete demographic and life events data relevant for our analyses (see Figure S1 A-C for details of the inclusion pipelines). We randomly excluded members of the same family to ensure no siblings were included in the sample. Overall, we included 6,366 subjects at 2-year follow-up (2,924 female adolescents), 5,035 subjects at 4-year follow-up (2,334 female adolescents), and 5,953 subjects at 6-year follow-up (2,834 female adolescents). Demographic information for each timepoint is reported in Table S1 A-C.

### Neuroimaging data

Structural T1-weighted neuroimaging data were collected using 3T MRI scanners. The neuroimaging protocol and parameters for the acquisition of T1-weighted images have been detailed in prior work (Casey et al., 2018; Hagler et al., 2019). We only included subjects whose structural neuroimaging data were recommended for inclusion (’mr_y_qc incl smri t1_indicator’=1) and had no clinically significant findings (’mr_y_qc clfind_score’=1 or 2). We used measures of total surface area, total grey matter volume, and mean cortical thickness based on 34 bilateral cortical regions (68 in total) defined by the Desikan-Killiany atlas (Desikan et al., 2006). We also included subcortical grey matter volumes covering 19 regions (bilaterally: accumbens, amygdala, cerebellar cortex, caudate, hippocampus, pallidum, putamen, thalamus, ventral diencephalon; unilaterally: brainstem). Area and volume measures, but not thickness, were proportionally corrected for individual differences in total intracranial volume. To ensure generalizability, we repeated our cortical analyses using the Destrieux atlas (Destrieux et al., 2010), which includes 74 bilateral cortical regions (148 total). All neuroanatomical measures were provided by the ABCD study’s standard FreeSurfer preprocessing pipeline.

### Adverse life events questionnaire

The ABCD study uses the Adverse Life Events Scale of the PhenX Toolkit to measure exposure and appraisal of adverse life events (PhenX Toolkit, *Life Events - Child*. https://www.phenxtoolkit.org/protocols/view/211501; Tiet et al. (1998). The scale consists of 25 items administered directly to the child/adolescent (youth-report) and the parent or primary caregiver (caregiver-report), individually, covering various events (e.g., injuries, death, crime, illness, relocations, new siblings) experienced during the previous year. If an event occurred, follow-up questions are asked about appraisal and impact (“Was this a good or a bad experience?”/”Was this a good or a bad experience for your child?”; “How much did the event affect you?”/”How much did the event affect your child?”). After the initial administration based on the 25 items at the 2-year follow-up, some items were added at the 4-year and 6-year follow-ups and are thus unavailable at the 2-year follow-up (e.g., school shootings, homelessness, deportation, parental hospitalization, foster care, and witnessing violence; detailed description of items added at later time points: https://docs.abcdstudy.org/latest/documentation/non_imaging/mh.html#mh_y_ple). Given that the adverse life events questionnaire assesses lifetime occurrence of events, later time points likely reflect events that were already reported at earlier assessments, resulting in a cumulative signal over time.

We focused on six summary variables: (1) the **number of total events** (count of endorsed events), (2) the **impact of total events** (sum of impact ratings for all endorsed events on a scale of 0 (“Not at all”) to 3 (“A lot”)), (3) the **number of positively appraised events** (count of endorsed events rated as “good”), (4) the **impact of positively appraised events** (sum of impact rating for endorsed events rated as “good”), (5) the **number of negatively appraised events** (count of endorsed events rated as “bad”), and (6) the **impact of negatively appraised events** (sum of impact rating for endorsed events rated as “bad”). Summary variables of impact reflect the count of events weighted by how much each event affected the individual. Data for these six summary variables were collected from either the youth or the caregiver. We only included participants for whom data from both sources were available, resulting in each of the six summary variables being represented twice (once from the youth report and once from the caregiver report) for a total of 12 variables. Although we used a cross-sectional design in the current study, the adverse life events are assessed cumulatively, so earlier occurrences should be reflected in all subsequent assessments.

### Descriptive analyses

We assessed sex differences, reporter differences, and appraisal differences using t-tests comparing across sexes, across youth- and caregiver-reported measures, and across positively and negatively appraised events, respectively.

### Brain-Based Predictive Modeling

We used a cross-validated brain-based predictive modeling framework to establish associations between neuroanatomical measures and adverse life events. This framework minimizes the risk of overfitting and prevents data leakage, while capturing robust and interpretable associations between neuroimaging-derived measures and phenotypic data (Dhamala et al., 2023). Here, we used linear ridge regression to predict the summary variables based on regional neuroanatomical measures. We trained separate models for each combination of neuroanatomical measure and adverse life event variable at each time point. Additionally, we developed sex-independent and sex-specific models to explore potential sex-specific associations between adverse life events and neuroanatomy. Overall, the combination of four neuroanatomical measures, 12 target variables (6 youth-report and 6 caregiver-report summary variables) and three groups (sex-independent group (the entire sample), and sex-specific groups (female or male adolescents only)) and three time points led to 432 models in total (Figure S2).

For each model, we performed 100 iterations of a random train-test (80%-20%) split. For each iteration, participants from the same site were not split across train/test sets to account for potential variability introduced by the imaging site. Within each train-test split, we optimized the regularization hyperparameter using a 5-fold cross-validation, ensuring that participants from a given site were not split across folds. After hyperparameter optimization in the training set, the model was fit to the training data using the optimized hyperparameter and evaluated on the corresponding test set. Model performance was assessed using prediction accuracy, defined as the correlation between the predicted values and the observed values, consistent with prior work (Chen et al., 2022, 2023; Dhamala et al., 2023). The 100 iterations of the train-test split resulted in a distribution of prediction accuracy values for each set of models.

To evaluate the significance of each set of models, we compared the distribution of prediction accuracy values to corresponding distributions of null models. Null models were generated by randomly permuting the target variable (the respective adverse life events summary variable) within each site and using the permuted data to train and test models using the optimized hyperparameters. This process was repeated for 1,000 train-test splits, generating a null distribution of accuracy values. We calculated p-values for model performance as the proportion of null models with prediction accuracies greater than or equal to the mean of the original model’s accuracy.

We corrected p-values for multiple comparisons across the 12 target variables for the respective groups (whole group, male adolescents only, female adolescents only) individually using the Benjamini-Hochberg False Discovery Rate procedure (Benjamini & Hochberg, 1995). Results were considered statistically significant if *p_FDR_* ≤ 0.05.

### Feature weights

To increase interpretability and reliability, we transformed the feature weights obtained from the models using the Haufe transformation (Haufe et al., 2014). We then calculated mean feature importance for each set of models. We assessed similarity between feature weights of the models using Pearson correlations.

### Sensitivity Analysis

Zero inflation in the data can influence brain-based predictions. We examined the distribution of all target variables to identify potential zero-inflation and conducted sensitivity analyses excluding zeros from each target variable individually to account for a potential zero-inflation (details provided in Supplementary Materials).

### Covariate analysis

Exposure to adverse life events is also related to socio-economic status (SES; Hatch & Dohrenwend, 2007; Lantz et al., 2005; Reiss et al., 2019). To quantify the influence of SES, we correlated SES with the target variables across all time points. To disentangle the effects of SES from the relationship between adverse life events and neuroanatomy, we repeated the brain-based predictive modeling analysis described above, incorporating SES, operationalized as income-to-needs ratio, as a covariate (see Supplementary Materials for details).

All analyses were conducted in Python (3.12.4) using Jupyter Notebook. Visualizations were generated using the seaborn package and the ENIGMA toolbox (Larivière et al., 2021).

## Results

### Descriptive analyses

Across all time points, youth generally reported greater numbers and impact of total and negatively appraised events compared to their caregivers, as well as greater numbers of negatively appraised events compared to positively appraised events (Figure S3, Table S2). At the same time, caregivers generally reported greater numbers of positively appraised events but a greater impact of negatively appraised events (Table S3). Further, caregivers were more likely to perceive certain events (e.g., “Someone in family died”; “Family member was seriously injured”; “Negative change in the parent’s financial situation”) as positive compared to youth (Table S4). Significant sex differences emerged at the 4-year follow-up, where female adolescents self-reported a higher number and impact of events. At the 6-year follow-up, female adolescents continued to exhibit higher numbers and impact of events as reflected in both youth and caregiver reports (Figure S4).

To address a potential zero-inflation in our target variables, we performed a sensitivity analysis by rerunning the analyses after individually removing zero values for each outcome variable. The amount of zeros varied depending on the target variable, with a generally higher proportion of zeros per target variable in the caregiver report compared to the youth report (Figure S5 A-C, Table S5, S6).

### Brain-Based Predictive Modeling

We used brain-based predictive models to capture associations between neuroanatomy and the number, appraisal, and impact of adverse life events at three different time points (2-year, 4-year, and 6-year follow-up).

### Main analyses

At the 2-year follow-up, cortical thickness predicted the impact of positively appraised events in female adolescents based on the caregiver report (*r*=0.114, *p_FDR_*=0.024), all other models were non-significant. At the 4-year follow-up, all models were non-significant. At the 6-year follow-up, cortical thickness, surface area, and volume models yielded accurate predictions across the entire sample (0.065<*r*<0.090, all *p_FDR_*<0.05) for number and impact of total events, and impact of positively and negatively appraised events in the youth report and for number and impact of positively appraised events in the caregiver report (Figure 1.1). Further, in male adolescents, only cortical thickness predicted the number (*r*=0.108, *p_FDR_*=0.042) and impact (*r*=0.108, *p_FDR_*=0.042) of positively appraised events based on the caregiver-report (Figure 1.1, with details in Figure S6).

**Figure 1.**
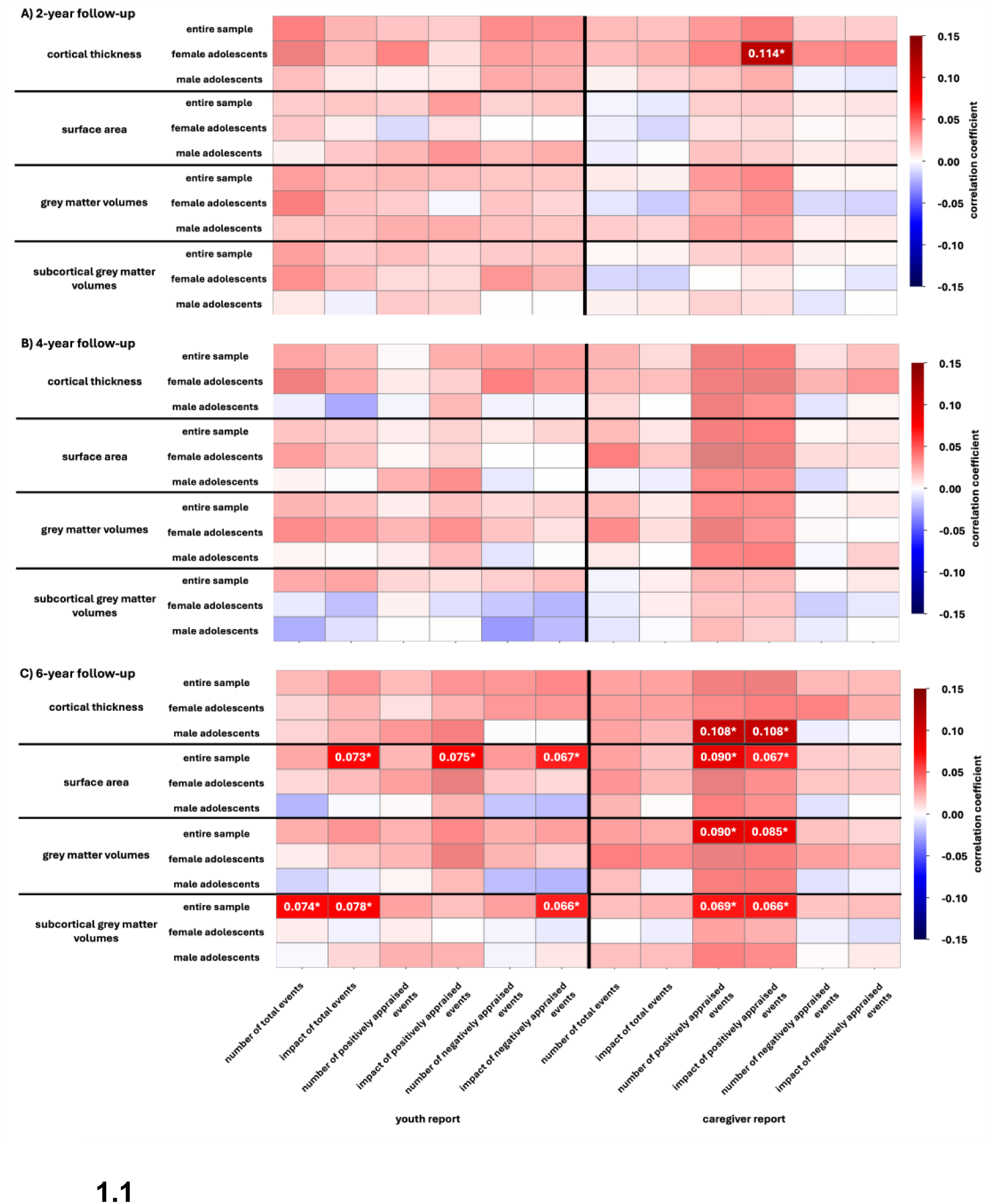
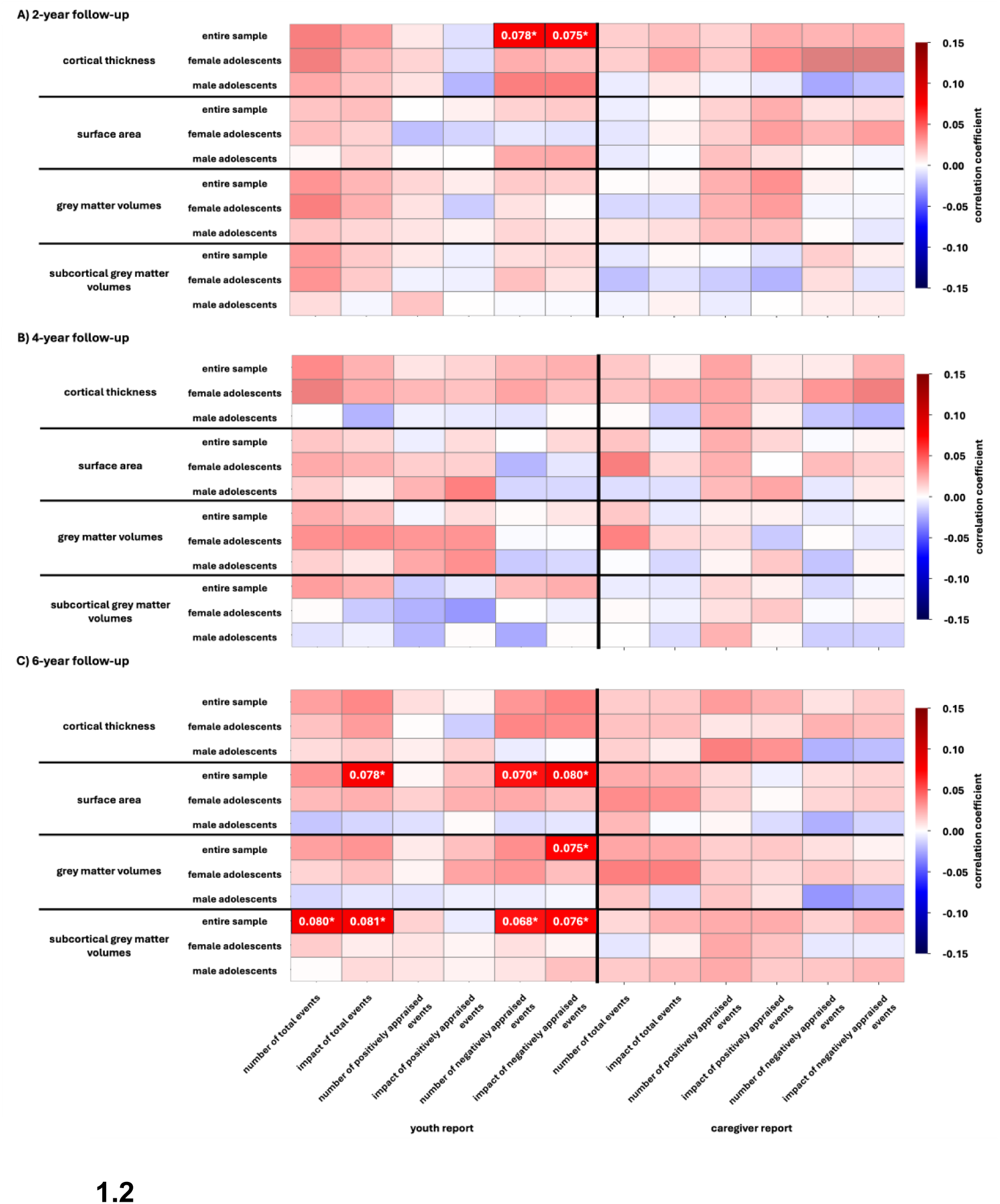
Predictive performance of neuroanatomical measures. These heatmaps illustrate the brain-based predictive modeling results for the (1.1) main analyses and the (1.2) sensitivity analyses. The mean Pearson correlation coefficients capture the association between predicted and actual data for models trained to predict the number, appraisal, or impact of adverse life events based on different neuroanatomical measurements (Cortical thickness, surface area, cortical and subcortical grey matter volumes) at the (A) 2-year follow-up, (B) 4-year follow-up, and (C) 6-year follow-up. The colors of the heatmap indicate the strength of the association depicted by the mean correlation coefficient. Negative associations are depicted in blue, positive associations are depicted in red. Measurements of cortical thickness, surface area, and cortical grey matter volumes are based on the Desikan-Killiany parcellation. Non-significant associations are depicted with reduced opacity. Significant predictions are labeled with their prediction accuracy.

### Sensitivity analyses

At the 2-year follow-up, the sensitivity analysis accounting for zero values yielded significant predictions for the entire sample based on cortical thickness for the number (*r*=0.078, *p_FDR_*=0.042) and impact of negatively appraised events (*r*=0.075, *p_FDR_*=0.042). At 4-year follow-up, all models were non-significant. At the 6-year follow-up, surface area and volume models yielded accurate predictions across the entire sample (0.068<*r*<0.081, all *p_FDR_*<0.05). All significant predictions in the sensitivity analyses were based on the youth report (Figure 1.2, details in Figure S7).

### Feature weights

#### Main analyses

We also examined the features that were important for these predictions. At the 2-year follow-up, the impact of positively appraised events was largely positively linked to cortical thickness in female adolescents (i.e., greater impact = greater thickness), with the strongest associations in the right transverse temporal gyrus, right insula, and bilaterally in the cuneus, pericalcarine, and lingual gyri (Figure 2A). At 6-year follow-up, feature weights for number and impact of adverse life events were predominantly positive and linked to cortical thickness, surface area, and volume, with the strongest associations located in the rostral middle frontal, superior frontal, lateral occipital gyrus, cuneus, and pericalcarine, as well as the thalamus and putamen (Figure 2-5). Distributions of feature weights were predominantly similar within each modality (Figure S8).

**Figure 2.**
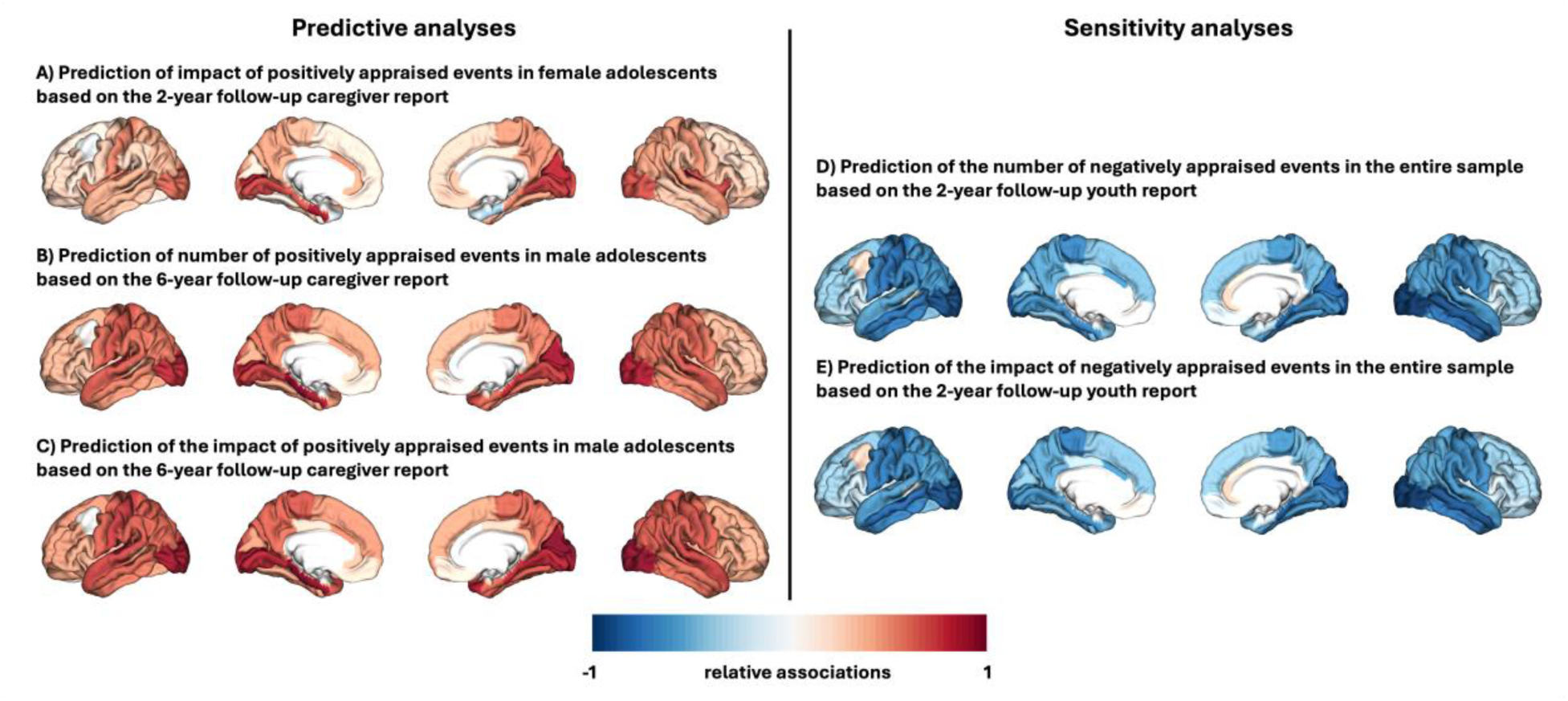
Distributions of feature weights of significant predictions based on cortical thickness. Red colors indicate stronger positive associations, blue colors indicate stronger negative associations.

**Figure 3.**
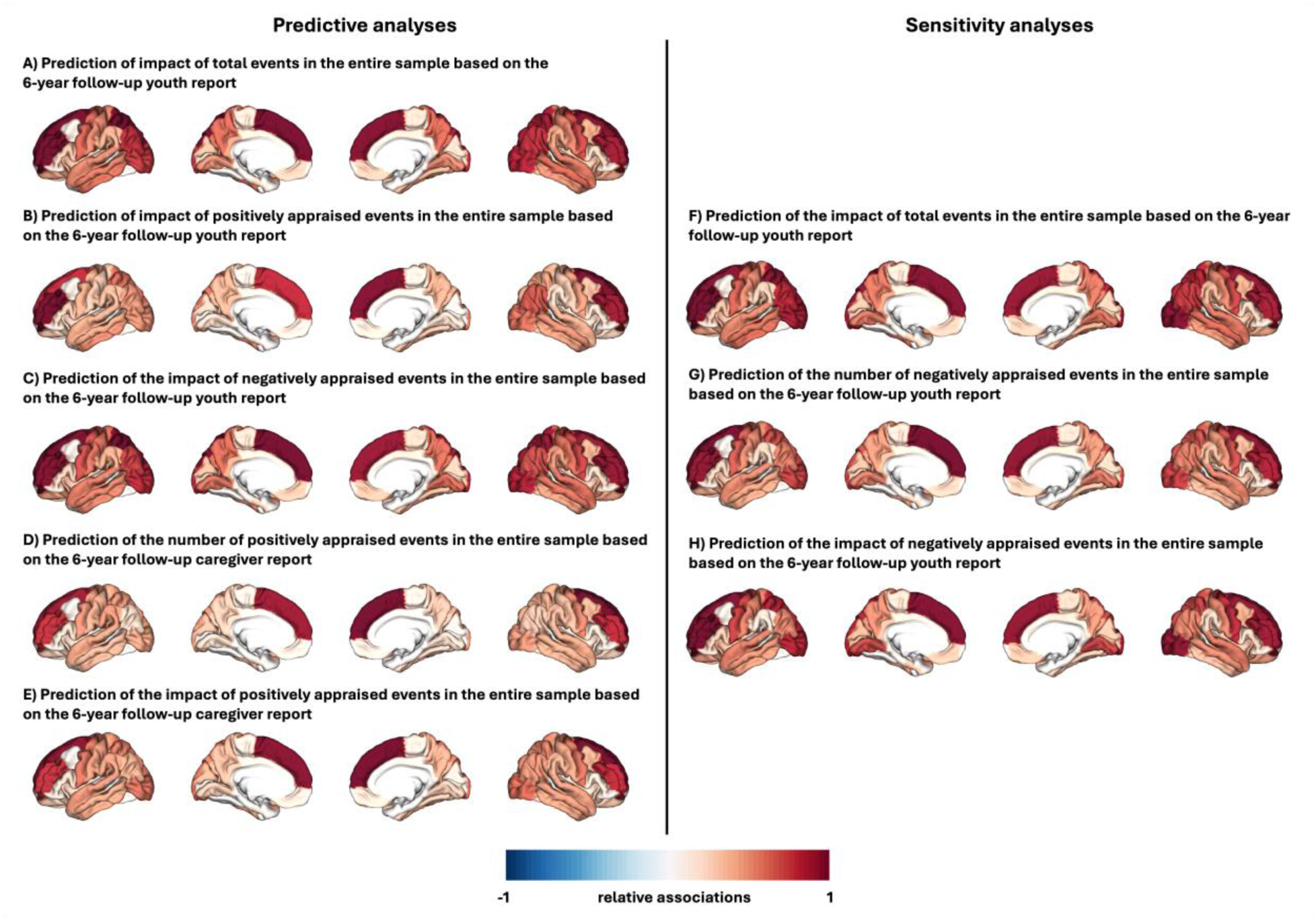
Distributions of feature weights of significant predictions based on surface area. Red colors indicate stronger positive associations, blue colors indicate stronger negative associations.

**Figure 4.**
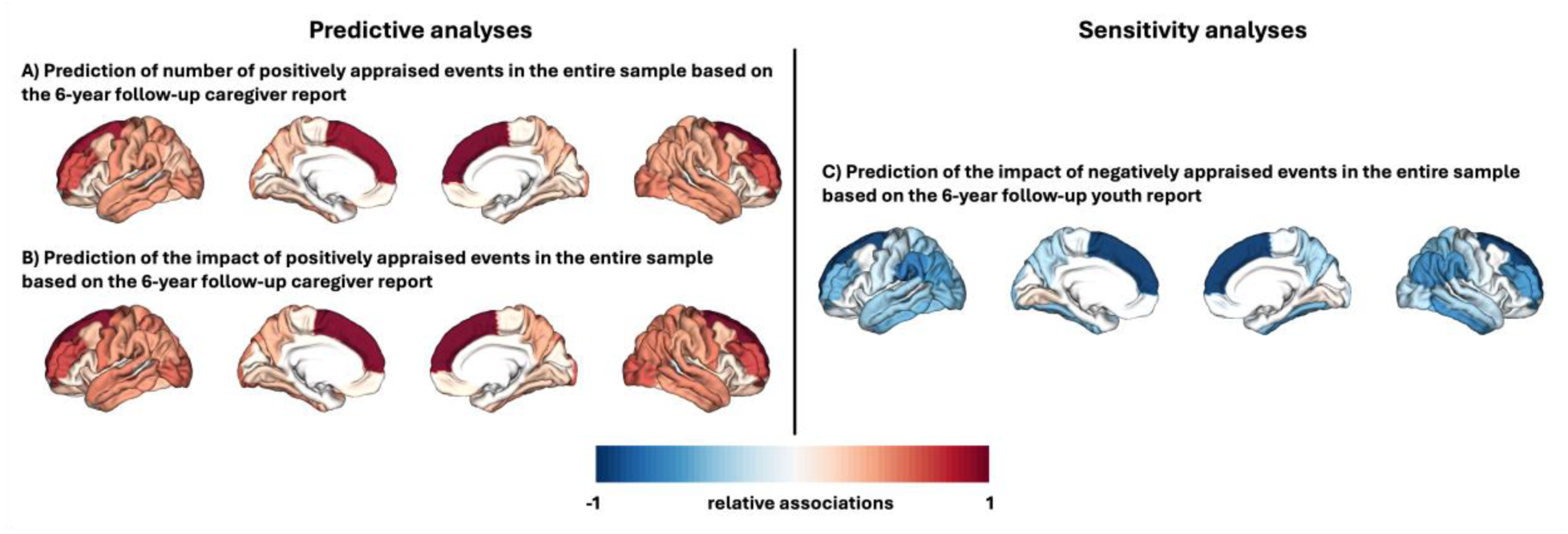
Distributions of feature weights of significant predictions based on grey matter volumes. Red colors indicate stronger positive associations, blue colors indicate stronger negative associations.

**Figure 5.**
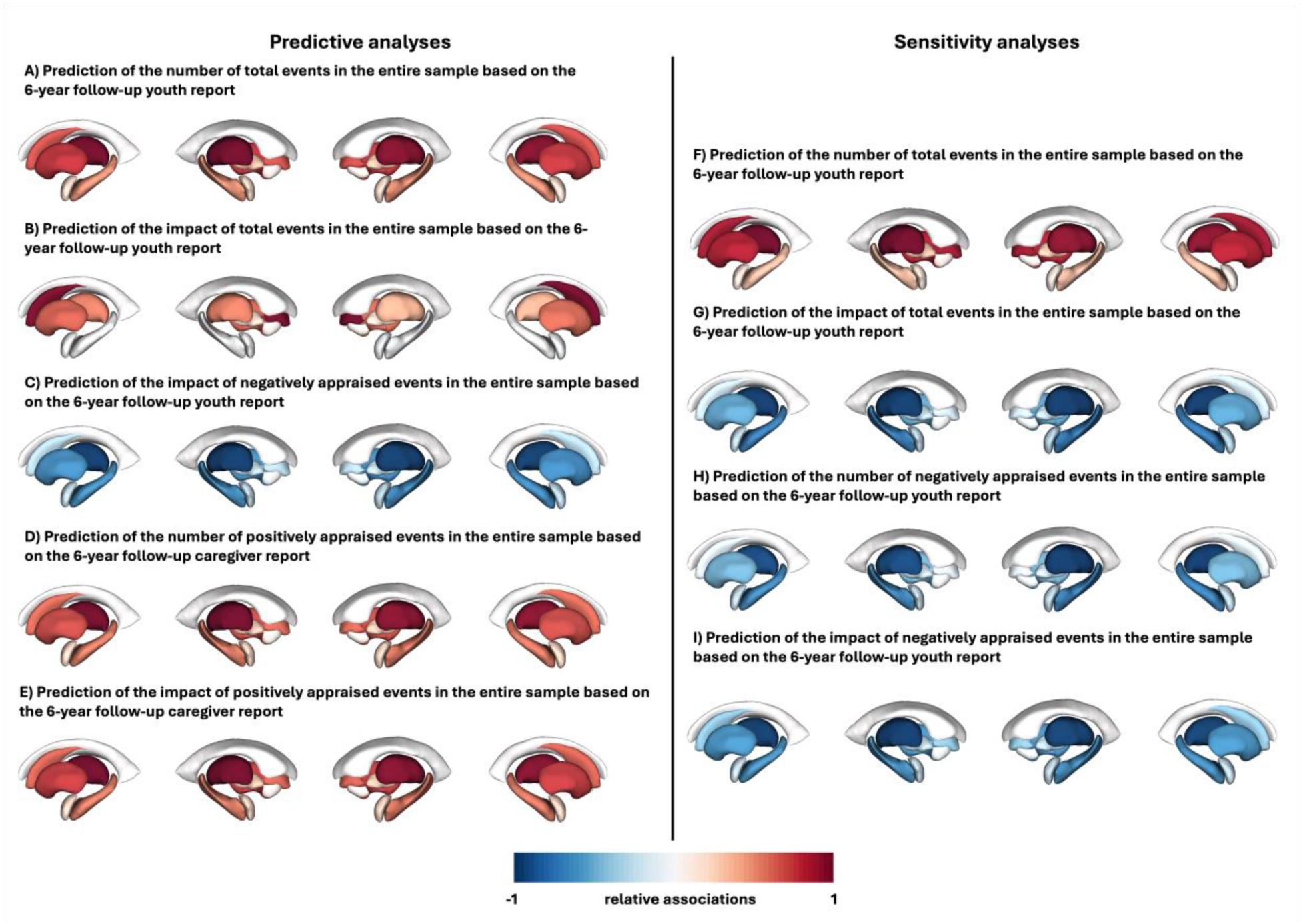
Distributions of feature weights of significant predictions based on subcortical grey matter volumes. Red colors indicate stronger positive associations, blue colors indicate stronger negative associations.

#### Sensitivity analyses

At the 2-year follow-up, the sensitivity analyses accounting for zero values showed that the number and impact of negatively perceived events were largely negatively linked to cortical thickness in the entire sample based on the youth report, with the strongest associations in the rostral and caudal anterior cingulate, isthmus cingulate, posterior cingulate, and medial orbitofrontal cortex (Figure 2D, 2E). At the 6-year follow-up, the number and impact of total and negatively perceived events were linked to surface area and volume based on the youth report. Associations were predominantly positive for surface area, with the strongest associations located bilaterally in the superior frontal, rostral middle frontal, lateral occipital, and superior parietal cortices (Figure 3F-H). In contrast, associations were predominantly negative for volume, with the highest feature weights located in the lingual, pericalcarine, caudal and rostral anterior cingulate, caudal middle frontal cortex, and the left cuneus for cortical volumes (Figure 4C) and the thalamus, ventral diencephalon, and hippocampus for subcortical volumes (Figure 5).

Distributions of feature weights were predominantly similar within each modality for sensitivity analyses. Comparing feature weight distributions between the main analyses and sensitivity analyses revealed similar patterns for surface area, but a converse pattern for thickness and volume (Figure S8).

To assess the robustness of our findings, all analyses were repeated using the Destrieux parcellation. The Destrieux-based models not only replicated the significant predictions obtained with the Desikan parcellation, but also revealed additional significant associations, which may be attributable to its higher spatial resolution (<u>Hastie</u> <u>et al., 2009</u>; Figure S9 and S10).

#### Covariate analyses

To control for potential confounding effects of SES, we analyzed the relationships between SES and the target variables, and repeated the predictive modeling analyses incorporating SES as a covariate. SES consistently showed an inverse relationship with the number and impact of total and negatively appraised events, and a positive relationship with the number and impact of positively appraised events (see Table S7 for full details).

Models controlling for SES yielded fewer significant predictions: at the 6-year follow-up, subcortical volumes predicted the impact of total events based on the youth report (*r*=0.079, *p_FDR_*=0.048) across the entire sample, but all other predictions were non-significant. (Details provided in Figure S11). In contrast, the Destrieux parcellation yielded more significant associations across the entire sample (Figure S12): at the 6-year follow-up, cortical thickness predicted the impact of total events (*r*=0.084, *p_FDR_*=0.02), as well as number (*r*=0.084, *p_FDR_*=0.02) and impact of negatively appraised events (*r*=0.105, *p_FDR_*=0.012) based on the youth report, as well as the impact of total events based on the caregiver report (*r*=0.066, *p_FDR_*=0.045). Surface area predicted the impact of total events (*r*=0.080, *p_FDR_*=0.028) and negatively appraised events (*r*=0.076, *p_FDR_*=0.018) in the youth report and impact of total events based on the caregiver report (*r*=0.078, *p_FDR_*=0.018).

## Discussion

Here, we used brain-based predictive models to investigate sex-independent and sex-specific associations between adverse life events and neuroanatomy across three time points during development covering the ages of 11-12, 13-14, and 15-16. We assessed cortical thickness, surface area, and cortical and subcortical grey matter volume, along with self- and caregiver-reported number, appraisal, and impact of adverse life events. Although significant associations were sparse at the 2-year follow-up and absent at the 4-year follow-up, all neuroanatomical measures were significantly linked to different measures of adverse life events at the 6-year follow-up. Our findings suggest that the associations between adverse life events and neuroanatomy emerge in mid-adolescence (ages 15-16). This might reflect either a developmentally sensitive time period at that age during which these associations are detectable or the progressive accumulation of adverse life events over time reaching a detectable signal at a later time point. Furthermore, our analyses revealed differential results between youth and caregiver reports, demonstrating the need to accurately differentiate between reporting sources.

Cortical thickness exhibited sex-specific associations with adverse life events for the number and impact of positively appraised events. While the association for female adolescents was significant at the 2-year follow-up, significant associations for male adolescents emerged at the 6-year follow-up. However, all predictions relied on overlapping feature weights (Figure 2 and Figure S8A), with predominantly positive associations located in occipital and temporoparietal regions. In previous literature, these regions have been linked to adverse life events but also to resilience. Resilience is positively associated with cortical thickness in the lateral occipital cortex, the fusiform gyrus, the inferior parietal cortex, as well as the middle and inferior temporal cortex (Kahl et al., 2020). Our results indicate that while the same brain regions are implicated across the sexes, the associations emerge earlier for female adolescents.

Surface area was linked to adverse life events across multiple target variables in the entire sample. At the 6-year follow-up, associations were significant for the impact of total, positively, and negatively appraised events in the youth report and for the number and impact of positively appraised events in the caregiver report. These predictions relied on overlapping feature weights with predominantly positive associations in the frontal, parietal, and occipital lobe (Figure 3, Figure S8B). The overlap in feature weights suggests that surface area reflects number and impact of adverse life events rather than differences in appraisal. The positive associations with neuroanatomy in our results contrast with previous literature predominantly reporting decreased surface area linked to adversity exposure such as maltreatment history (Kelly et al., 2013) or trauma exposure and household abuse (Thorsen et al., 2025), which are most likely perceived as negative life events. These discrepancies likely stem from variations in the conceptualization and measurements of adversity across studies. The impact of such nuanced distinctions was shown in a study by (Garrisi et al., 2024); their analyses demonstrated decreased surface area after deprivation exposure in higher-order cognition areas such as the orbitofrontal and superior parietal cortices, whereas after threat exposure, surface area increased in regions involved in socio-emotional tasks such as the inferior temporal and parietal cortices.

Volume and adverse life events exhibited nuanced patterns of associations for cortical and subcortical volumes. For cortical volume, significant predictions were based on the caregiver report only for positively appraised events, with overlapping positive feature weights in the superior frontal, rostral middle frontal, precentral, and lateral occipital gyri (Figure 4, Figure S8C). In contrast, for subcortical volume, significant predictions were based on both youth and caregiver reports for total, negatively, and positively appraised events with dispersed positive and negative feature weights (Figure 5, Figure S8D). Prior studies have reported decreases in grey matter volumes in these cortical and subcortical areas following adverse or stressful experiences (Ansell et al., 2012; Frodl & O’Keane, 2013; Lim et al., 2014; Lim et al., 2018; Tyborowska et al., 2018). The present findings align with these observations considering positively appraised life events were positively associated with neuroanatomical measurements.

The sensitivity analysis, which accounted for zero inflation in the data, yielded significant predictions only for the youth report for number and impact of total and negatively appraised events. The exclusion of zero values has important implications for the interpretation of results, as it reframes the analytic question from examining whether adverse life events are associated with neuroanatomy to examining whether the extent of exposure is associated with neuroanatomy among those who have experienced such events. Notably, zero values may not be uninformative, particularly in the caregiver report, where a value of zero could indicate either true non-exposure or a failure to detect events experienced by the youth. For surface area, the sensitivity analyses demonstrated a similar distribution of feature weights as the main analyses, which aligns with findings of Buimer et al. (2022) that adverse life events are associated with larger cortical surface area in similar frontal areas. In contrast, the pattern of feature weights for cortical thickness was reversed compared to the main analyses: occipital and temporoparietal regions still exhibited the highest feature weights but had a negative instead of positive association. These negative associations align with prior research linking lower cortical thickness in the orbitofrontal cortex with adverse life events (Backhausen et al., 2024; Monninger et al., 2020). Similarly, for predictions based on cortical and subcortical volumes, we observed predominantly negative associations with number and impact of total and negatively appraised events, which align with literature reporting lower grey matter volumes to be associated with adverse life experiences and early-life stress (Ansell et al., 2012; Frodl & O’Keane, 2013; Lim et al., 2014; Lim et al., 2018; Tyborowska et al., 2018). The difference between the modalities might stem from their different developmental trajectories, where surface area continues to expand into adolescence and peaks later than cortical thickness and grey matter volumes (Bethlehem et al., 2022).

Disparities between youth and caregiver reports were evident across our analyses: the main analyses based on the youth report were significant for total, positively and negatively appraised events, while those based on the caregiver report were significant for positively appraised events only. This pattern was partly reflected in the descriptive analyses, where youth reports had higher mean values for all target variables. However, a detailed analysis of the distribution of target variables showed that, despite lower mean values, caregivers report higher maximum values for the number and impact of positively appraised events, indicating a skewed distribution that might lead to higher signal for the main analyses (Table S2). Looking into the differences between youth and caregiver reports on a single-item level (Table S4) reveals patterns that might explain the characteristics of the target variables: (1) youth are more likely to report events as having occurred in the first place; (2) youth and caregiver reports show discrepancies in terms of appraisal, as some items were more negatively appraised by youth and more positively appraised by caregivers; and (3) youth tend to rate the severity of individual events as higher than caregivers (Table S4). These results align with findings of previous studies demonstrating that children report more negative events compared to their parents (Bailey & Garralda, 1990; Johnston et al., 2003; Schreier et al., 2005; Shemesh et al., 2005; Tingskull et al., 2015) and rate the severity of such events differently. Consequently, the findings of our descriptive, main, and sensitivity analyses highlight the disparity between youth and caregiver reports regarding the acknowledgement of adverse life events, their appraisal and their perceived impact.

Furthermore, the sensitivity analyses revealed that the significant findings based on the caregiver report in the main analyses were largely driven by the prevalence of zero values in the data. This difference emphasizes the need to analyze reporting sources individually to capture the distinct perspectives (Rende & Plomin, 1991), especially given that parents’ lack of recognition of a child’s adverse experiences can hinder intervention and thus result in poorer mental health outcomes (Schreier et al., 2005; Tingskull et al., 2015).

Importantly, interpreting the association between neuroanatomy and adverse life events requires considering a broader context and other mediating or moderating variables, such as SES. SES introduces a challenge for isolating the neuroanatomical correlates of stress, given its intricate relationship with adverse life events (e.g., homelessness, eviction) and its impact on neurodevelopment (Rakesh et al., 2023; Tsomokos et al., 2026). To quantify the impact of SES, we reran our analyses including SES as a covariate. This led to a notable reduction in significant associations, and only one association remained significant based on the Desikan parcellation, while a few remained significant based on the Destrieux parcellation. This shows that SES substantially affects the association between adverse life events and the brain, which aligns with the correlations between SES and the target variables (Table S7). Some adverse life events are directly linked to SES, such as “negative changes in the parent’s financial situation” or “parental figure lost job”, whereas others might be more indirectly linked. Beyond exposure to adverse life events, SES may also influence the degree to which these events affect children: families with greater socioeconomic resources might have greater access to resources - such as mental health services and stable living conditions - that can reduce the relative impact on children and their well-being.

Controlling for SES also reduced statistical power to capture the association between adverse life events and neuroanatomy, particularly in lower-dimensional feature sets such as the Desikan parcellation compared to the Destrieux parcellation. Our results confirm the intertwined relationship between SES and adverse life events, aligning with literature demonstrating shared neural correlates of adversity-related variables and SES (Brosch et al., 2025).

It is important that our findings are interpreted in the context of some key limitations. First, our operationalization of stress was limited to specific events captured by the Adverse Life Events scale, which does not fully capture all factors that are related to adversity or other stressful life events. Furthermore, summary scores reflecting numbers only do not account for the differential impact on individuals’ lives. To address this limitation, we included the impact variables that combine event occurrences with youth/caregiver ratings of how much each event affected the children in their personal lives. Second, while our analyses examined multiple neuroanatomical measures and stress summary scores, we exclusively used linear ridge regression and did not test other nonlinear algorithms, which could potentially yield different results. Third, while the ABCD study is a large-scale and heterogeneous dataset collected at multiple sites across the U.S., its exclusive focus on the U.S. may limit the generalizability of our results across international populations (Dhamala et al., 2024; Ricard et al., 2023).

Fourth, we assessed these associations in a cross-sectional design and not a longitudinal design. Future studies could adopt a longitudinal design to examine the developmental trajectories underlying stress-neuroanatomy associations. Additionally, we excluded siblings from the analyses to avoid family relatedness inflating prediction accuracies, resulting in smaller sample sizes. We acknowledge that alternative approaches, such as including a random intercept for family, would be statistically more robust and could be implemented in future work.

In summary, we assessed the neuroanatomical basis of the frequency, appraisal, and impact of adverse life events using a multivariate brain-based predictive modeling approach covering multiple time points during development. Our results demonstrated a predominantly late manifestation of adverse life events in various neuroanatomical measurements. Our analyses also demonstrated differences in reporting sources between youth and caregiver reports, especially in terms of appraisal, highlighting the importance to carefully consider reporting sources as a source of variability when studying adverse life events. Altogether, our findings show the need to integrate reporting sources, appraisal, different neuroanatomical modalities, and multiple time points across adolescence to fully capture the nuanced manifestations of adverse life events in brain structure.

## Implications & Contributions

● The present study leveraged data from multiple time points of a large-scale dataset to study the association between adversity and neuroanatomy, encompassing the number, appraisal, and impact of adverse life events and various neuroanatomical measurements.
● Our findings demonstrate that positively and negatively appraised events uniquely map onto neuroanatomy and that these associations emerge later in adolescence.
● This study also illustrated important differences between youth and caregiver reported data, demonstrating the need to differentiate between reporting sources.

## Author contributions

**LW**: Conceptualization, Writing – original draft, Software, Formal Analysis, Visualization; **KB**: Writing – review & editing; **EC**: Writing – review & editing; **ED**: Methodology, Software, Writing – review & editing, Supervision, Funding acquisition

## Ethical information

The research protocol for the dataset was reviewed and approved by a central Institutional Review Board (IRB) at the University of California, San Diego, and, in some cases, by individual site IRBs. Parents or guardians provided written informed consent, and children assented before participation.

## Data availability

Data used in the preparation of this article were obtained from the [Adolescent Brain Cognitive Development™ (ABCD) Study](https://abcdstudy.org/), held in the [NIH Brain Development Cohorts Data Sharing Platform](https://www.nbdc-datahub.org/). This is a multisite, longitudinal study designed to recruit more than 10,000 children aged 9–10 and follow them over 10 years into early adulthood. The ABCD Study® is supported by the **National Institutes of Health** and additional federal partners under award numbers: U01DA041048, U01DA050989, U01DA051016, U01DA041022, U01DA051018, U01DA051037, U01DA050987, U01DA041174, U01DA041106, U01DA041117, U01DA041028, U01DA041134, U01DA050988, U01DA051039, U01DA041156, U01DA041025, U01DA041120, U01DA051038, U01DA041148, U01DA041093, U01DA041089, U24DA041123, U24DA041147. A full list of supporters is available at [Federal Partners – ABCD Study](https://abcdstudy.org/federal-partners.html). ABCD Consortium investigators designed and implemented the study and/or provided data but did not necessarily participate in the analysis or writing of this report. This manuscript reflects the views of the authors and may not reflect the opinions or views of the NIH or ABCD Consortium investigators. The ABCD data repository grows and changes over time. The ABCD data used in this report came from the ABCD 7.0 release.

## Code availability

The code for data preparation, model training and computation of further analyses if available on Gitlab: https://gitlab.com/lwiersch/investigating-the-impact-of-stressful-life-events-on-neuroanatomy-across-adolescence

## Further acknowledgements

During the preparation of this work the authors used LLM to improve readability and language. After using this tool/service, the authors reviewed and edited the content as needed and take full responsibility for the content of the publication.

