## Supplementary_Material for "Investigating the impact of adverse life events on neuroanatomy across adolescence"

Wiersch et al.

**Supplementary Methods**

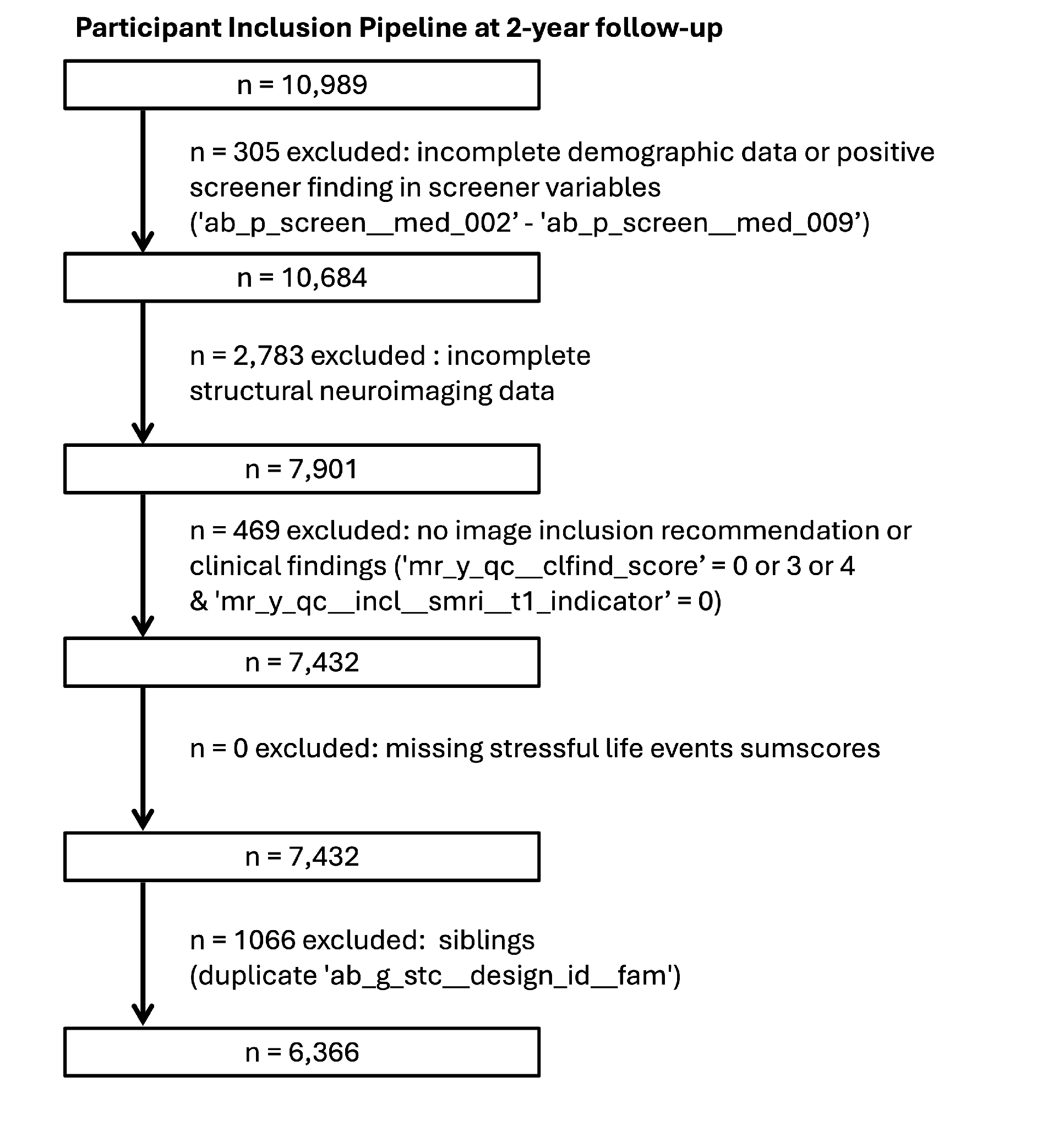

**Supplementary Figure 1A. Participant Inclusion and exclusion workflow at 2-year follow-up.**
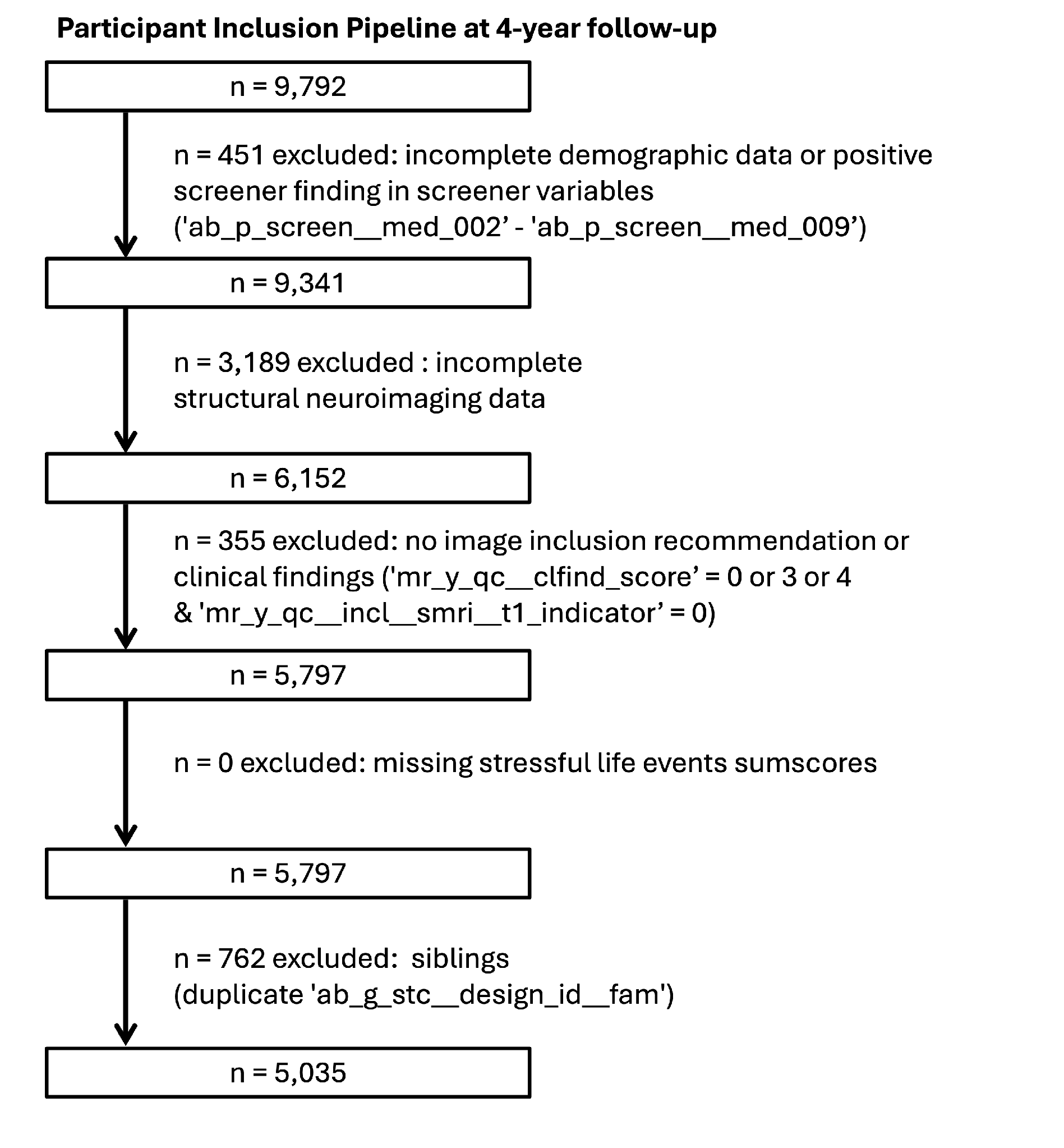

**Supplementary Figure 1B. Participant Inclusion and exclusion workflow at 4-year follow-up.**

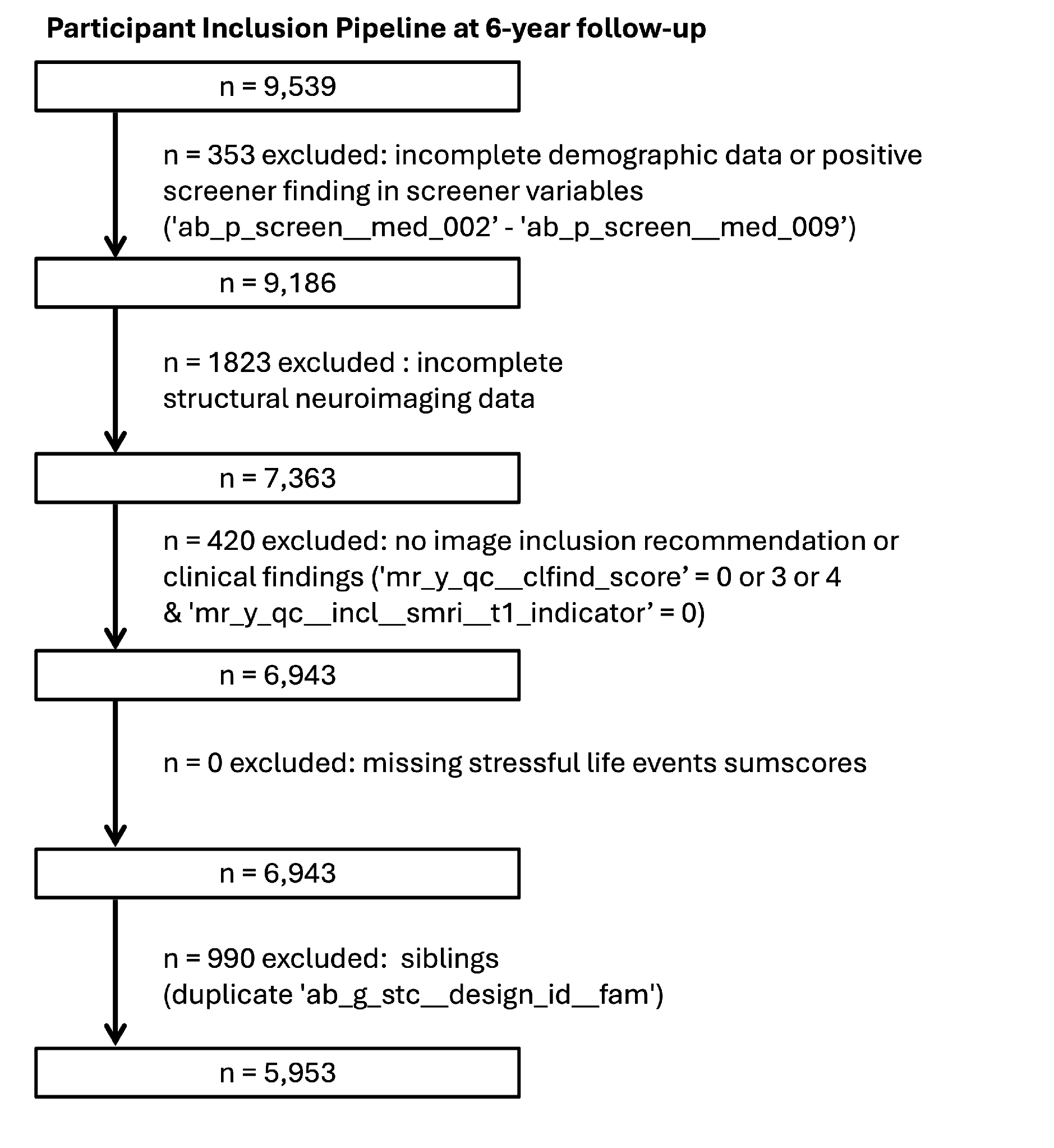

**Supplementary Figure 1C. Participant Inclusion and exclusion workflow at 6-year follow-up.**

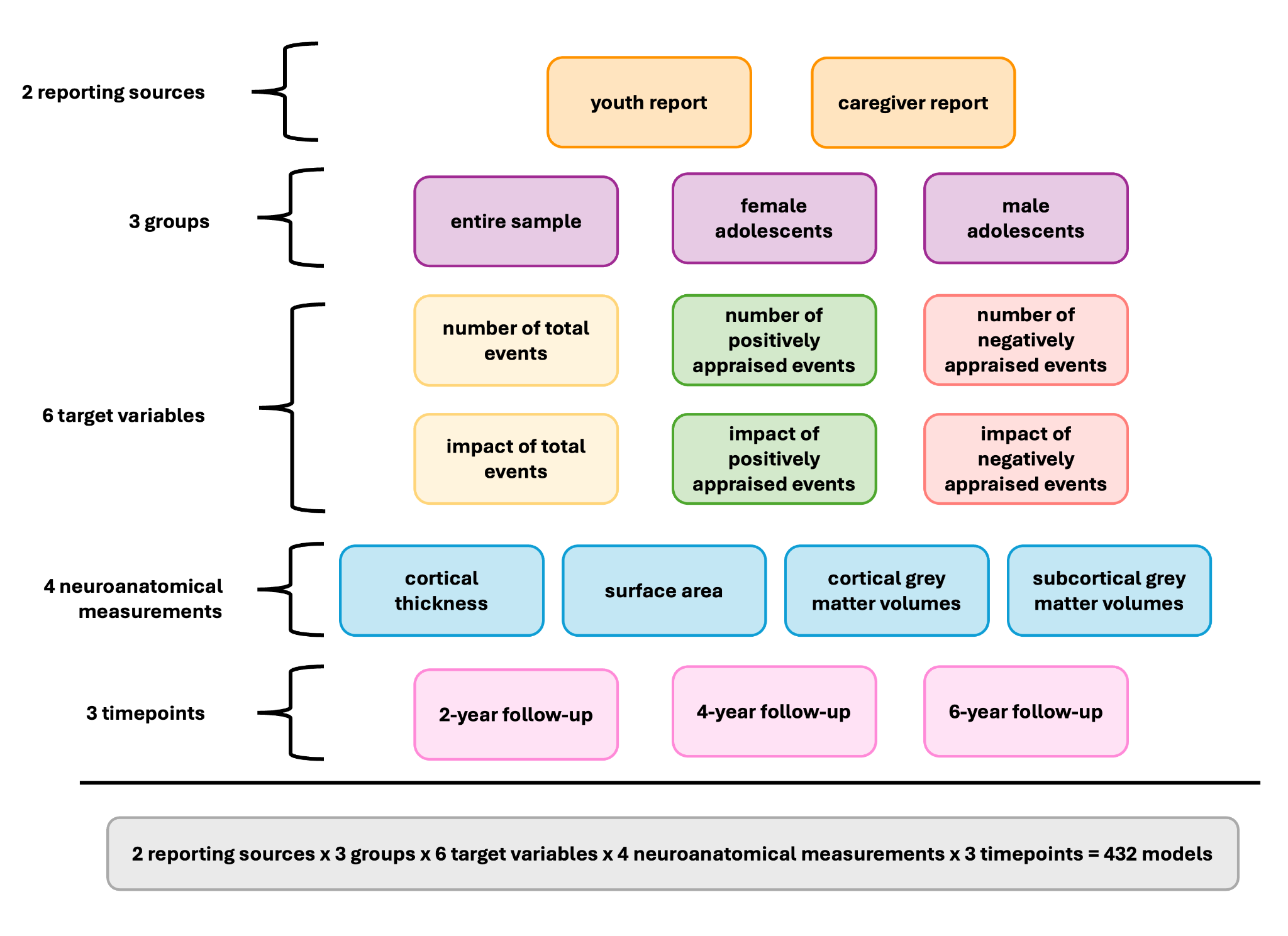

**Supplementary Figure S2. Analysis overview**

To further address a potential zero-inflation of the target variables, we repeated the brain-based predictive modeling analyses after individually filtering zero values from each target variable to maximize sample size retention. Individual sample sizes for the sensitivity analyses are provided in Table S6.

To control for potential confounding effects of socioeconomic status (SES) between stressful life events and neuroanatomical measurements, we repeated the brain-based predictive modeling analyses with SES included as a covariate. SES was approximated as the income-to-needs ratio.

Income was indexed by the combined household income ('ab_p_demo__income__hhold_001__v01'), assessed across income brackets. We excluded persons reporting non-responses (999: Don’t know; 777: Decline to answer) and used the median income value per bracket as approximation for the combined family income.

Poverty thresholds were determined based on the household size (‘ab_p_demo__roster_001’). Participants were excluded if they provided non-informative responses for household size (ab_p_demo__roster_001__dk: 777: Decline to answer; 444: Not applicable) or reported a household size of zero. Based on the 2022 federal poverty guidelines (<https://aspe.hhs.gov/sites/default/files/documents/4b515876c4674466423975826ac57583/Guidelines-2022.pdf>), poverty thresholds were assigned to each participant in accordance with their household size. SES was subsequently operationalized by dividing the household income by the corresponding poverty threshold.

After excluding participants with non-informative responses in the relevant variables, removing siblings, and excluding sites with fewer than 10 participants, final sample sizes were N = 5806 for the 2-year follow-up, N = 4540 for the 4-year follow-up, and N = 5380 for the 6-year follow-up.

The associations between SES and each target variable for each time point were assessed using Spearman rank correlations.

**Supplementary Table 1. Demographic information**
Demographic information (age, ethnicity, and income) for all subjects included for the descriptive and predictive analyses. Demographic information is reported separately for male adolescents and female adolescents for the 2-year (A), 4-year (B), and 6-year follow-up (C). Reported proportions (%) may not sum to exactly 100% due to rounding.

A) **2-year follow-up (N = 6,366)**

|  | male adolescents | female adolescents |
| --- | --- | --- |
| Sample Size (%) | 3442 (54.07%) | 2924 (45.93%) |
| Age (mean, SD) | 12.00 (0.65) | 11.96 (0.65) |
| Ethnicity |  |  |
| Hispanic | 689 (20.02%) | 609 (20.83%) |
| White | 1911 (55.52%) | 1525 (52.15%) |
| Black | 461 (13.39%) | 420 (14.36%) |
| Asian | 62 (1.80%) | 51 (1.74%) |
| Other | 319 (9.27%) | 319 (10.91%) |
| Income Category (count, %) |  |  |
| Less than $5,000 | 96 (2.79%) | 81 (2.77%) |
| $5,000 through $11,999 | 110 (3.20%) | 68 (2.33%) |
| $12,000 through $15,999 | 70 (2.03%) | 57 (1.95%) |
| $16,000 through $24,999 | 113 (3.28%) | 97 (3.32%) |
| $25,000 through $34,999 | 172 (5.00%) | 168 (5.75%) |
| $35,000 through $49,999 | 223 (6.48%) | 222 (7.59%) |
| $50,000 through $74,999 | 423 (12.29%) | 386 (13.20%) |
| $75,000 through $99,999 | 439 (12.75%) | 387 (13.24%) |
| $100,000 through $199,999 | 1091 (31.70%) | 882 (30.16%) |
| $200,000 and greater | 438 (12.73%) | 360 (12.31%) |
| Don't know | 133 (3.86%) | 126 (4.31%) |
| Decline to answer | 134 (3.89%) | 90 (3.08%) |

B) **4-year follow-up (N = 5,035)**

|  | male adolescents | female adolescents |
| --- | --- | --- |
| Sample Size (%) | 2701 (53.64%) | 2334 (46.36%) |
| Age (mean, SD) | 14.15 (0.70) | 14.15 (0.72) |
| Ethnicity |  |  |
| Hispanic | 561 (20.77%) | 487 (20.87%) |
| White | 1454 (53.83%) | 1175 (50.34%) |
| Black | 375 (13.88%) | 369 (15.81%) |
| Asian | 48 (1.78%) | 53 (2.27%) |
| Other | 263 (9.74%) | 250 (10.71%) |
| Income Category (count, %) |  |  |
| Less than $5,000 | 59 (2.18%) | 55 (2.36%) |
| $5,000 through $11,999 | 64 (2.37%) | 50 (2.14%) |
| $12,000 through $15,999 | 45 (1.67%) | 43 (1.84%) |
| $16,000 through $24,999 | 94 (3.48%) | 71 (3.04%) |
| $25,000 through $34,999 | 135 (5.00%) | 116 (4.97%) |
| $35,000 through $49,999 | 163 (6.03%) | 165 (7.07%) |
| $50,000 through $74,999 | 294 (10.88%) | 242 (10.37%) |
| $75,000 through $99,999 | 322 (11.92%) | 279 (11.95%) |
| $100,000 through $199,999 | 843 (31.21%) | 750 (32.13%) |
| $200,000 and greater | 451 (16.70%) | 372 (15.94%) |
| Don't know | 114 (4.22%) | 85 (3.64%) |
| Decline to answer | 117 (4.33%) | 106 (4.54%) |

C) **6-year follow-up (N = 5,953)**

|  | male adolescents | female adolescents |
| --- | --- | --- |
| Sample Size (%) | 3119 (52.39%) | 2834 (47.61%) |
| Age (mean, SD) | 16.08 (0.67) | 16.05 (0.67) |
| Ethnicity |  |  |
| Hispanic | 648 (20.78%) | 574 (20.25%) |
| White | 1726 (55.34%) | 1499 (52.89%) |
| Black | 396 (12.70%) | 402 (14.18%) |
| Asian | 61 (1.96%) | 55 (1.94%) |
| Other | 288 (9.23%) | 304 (10.73%) |
| Income Category (count, %) |  |  |
| Less than $5,000 | 74 (2.37%) | 68 (2.40%) |
| $5,000 through $11,999 | 47 (1.51%) | 55 (1.94%) |
| $12,000 through $15,999 | 45 (1.44%) | 31 (1.09%) |
| $16,000 through $24,999 | 89 (2.85%) | 62 (2.19%) |
| $25,000 through $34,999 | 131 (4.20%) | 110 (3.88%) |
| $35,000 through $49,999 | 186 (5.96%) | 176 (6.21%) |
| $50,000 through $74,999 | 273 (8.75%) | 259 (9.14%) |
| $75,000 through $99,999 | 341 (10.93%) | 307 (10.83%) |
| $100,000 through $199,999 | 1075 (34.47%) | 996 (35.14%) |
| $200,000 and greater | 661 (21.19%) | 602 (21.24%) |
| Don't know | 84 (2.69%) | 66 (2.33%) |
| Decline to answer | 113 (3.62%) | 102 (3.60%) |

**Supplementary Results**

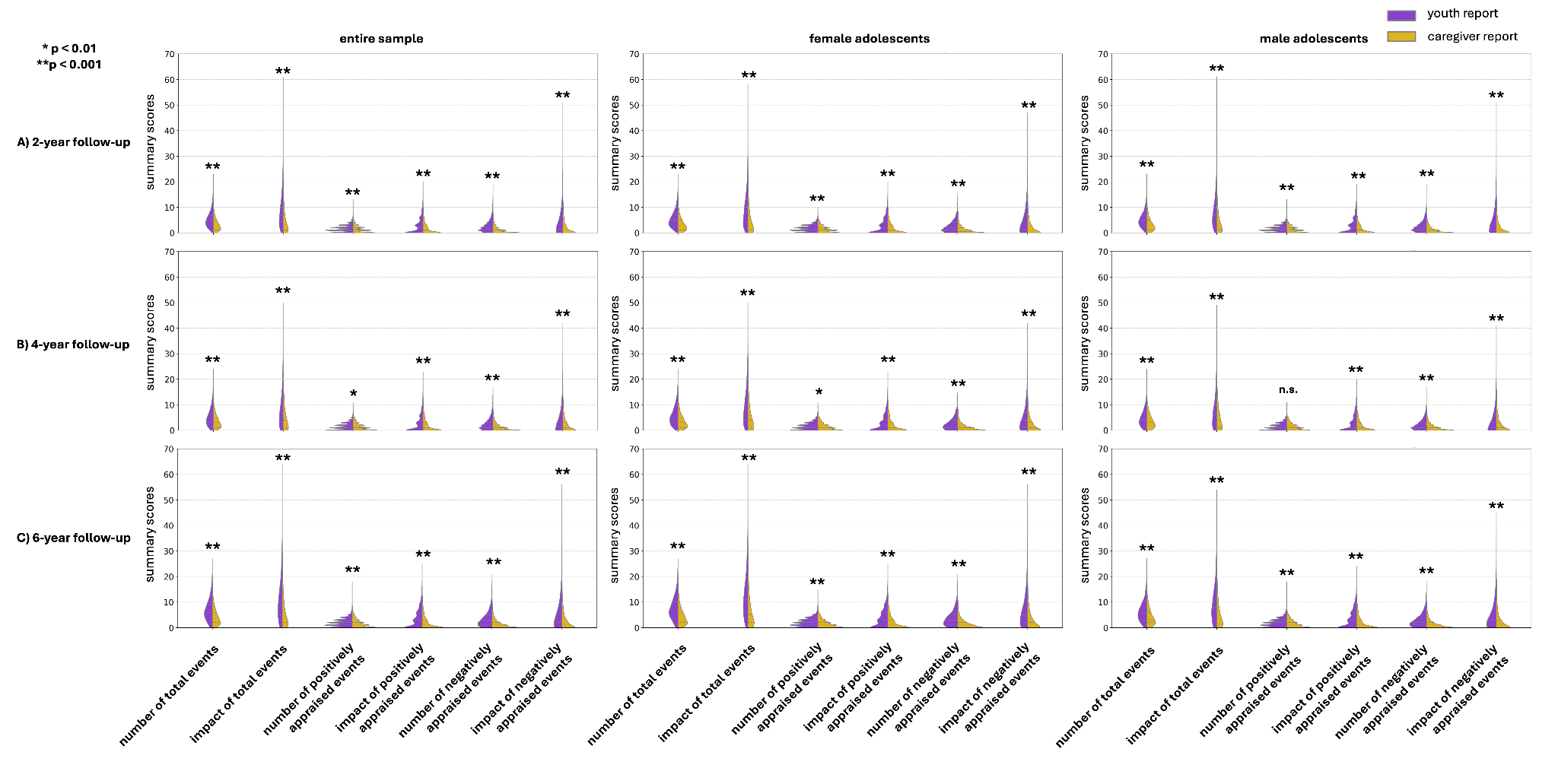

**Supplementary Figure 3. Differences between youth report and caregiver report.** Descriptive statistics and t-tests contrasting the youth and caregiver report at the (A) 2-year follow-up, (B) 4-year follow-up, (B) 6-year follow-up time point. Asterisks indicate a significant difference.

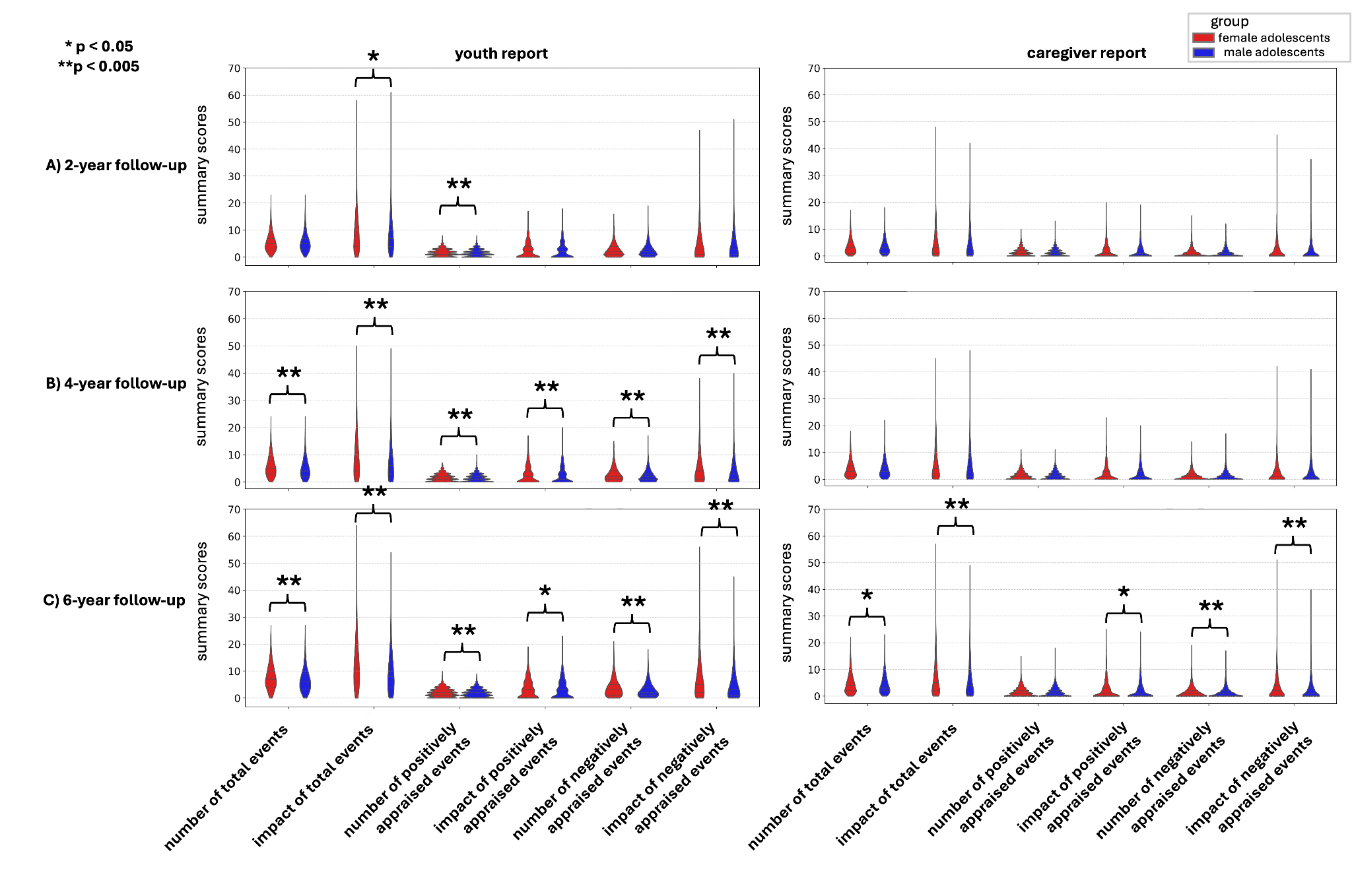

**Supplementary Figure 4. Sex differences in the distribution of summary variables.** Descriptive statistics and t-tests contrasts of sex-specific groups for stressful life events variables at the (A) 2-year follow-up, (B) 4-year follow-up, (B) 6-year follow-up time point for the youth report and caregiver report. Asterisks indicate a significant difference.

**Supplementary Table 2. Descriptive information**

Descriptive statistics of the entire sample, female adolescents and male adolescents for the (A) 2-year follow-up, (B) 4-year follow-up, and (C) 6-year follow-up time point for the youth report and caregiver report.

A) **2-year follow-up**

|  |  |  | **number of total events** | **impact of total events** | **number of positively appraised events** | **impact of positively appraised events** | **number of negatively appraised events** | **impact of negatively appraised events** |
| --- | --- | --- | --- | --- | --- | --- | --- | --- |
| **entire sample** | **youth report** | **mean** | 5.36 | 8.80 | 1.56 | 2.49 | 2.43 | 4.97 |
|  |  | **std** | 3.28 | 6.99 | 1.33 | 2.87 | 2.25 | 5.36 |
|  |  | **median** | 5.00 | 7.00 | 1.00 | 2.00 | 2.00 | 3.00 |
|  |  | **min** | 0.00 | 0.00 | 0.00 | 0.00 | 0.00 | 0.00 |
|  |  | **max** | 23.00 | 61.00 | 8.00 | 18.00 | 19.00 | 51.00 |
|  | **caregiver report** | **mean** | 3.39 | 4.96 | 1.29 | 1.68 | 1.08 | 2.19 |
|  |  | **std** | 2.69 | 5.50 | 1.40 | 2.42 | 1.65 | 3.85 |
|  |  | **median** | 3.00 | 3.00 | 1.00 | 1.00 | 0.00 | 0.00 |
|  |  | **min** | 0.00 | 0.00 | 0.00 | 0.00 | 0.00 | 0.00 |
|  |  | **max** | 18.00 | 48.00 | 13.00 | 20.00 | 15.00 | 45.00 |
| **female adolescents** | **youth report** | **mean** | 5.41 | 9.03 | 1.62 | 2.56 | 2.43 | 5.08 |
|  |  | **std** | 3.35 | 7.12 | 1.34 | 2.91 | 2.26 | 5.43 |
|  |  | **median** | 5.00 | 8.00 | 1.00 | 2.00 | 2.00 | 3.00 |
|  |  | **min** | 0.00 | 0.00 | 0.00 | 0.00 | 0.00 | 0.00 |
|  |  | **max** | 23.00 | 58.00 | 8.00 | 17.00 | 16.00 | 47.00 |
|  | **caregiver report** | **mean** | 3.44 | 5.09 | 1.33 | 1.73 | 1.11 | 2.27 |
|  |  | **std** | 2.72 | 5.61 | 1.42 | 2.44 | 1.69 | 3.98 |
|  |  | **median** | 3.00 | 3.00 | 1.00 | 1.00 | 0.50 | 0.00 |
|  |  | **min** | 0.00 | 0.00 | 0.00 | 0.00 | 0.00 | 0.00 |
|  |  | **max** | 17.00 | 48.00 | 10.00 | 20.00 | 15.00 | 45.00 |
| **male adolescents** | **youth report** | **mean** | 5.32 | 8.61 | 1.52 | 2.44 | 2.44 | 4.87 |
|  |  | **std** | 3.22 | 6.87 | 1.31 | 2.83 | 2.24 | 5.31 |
|  |  | **median** | 5.00 | 7.00 | 1.00 | 2.00 | 2.00 | 3.00 |
|  |  | **min** | 0.00 | 0.00 | 0.00 | 0.00 | 0.00 | 0.00 |
|  |  | **max** | 23.00 | 61.00 | 8.00 | 18.00 | 19.00 | 51.00 |
|  | **caregiver report** | **mean** | 3.36 | 4.86 | 1.26 | 1.63 | 1.05 | 2.13 |
|  |  | **std** | 2.67 | 5.41 | 1.39 | 2.41 | 1.61 | 3.73 |
|  |  | **median** | 3.00 | 3.00 | 1.00 | 1.00 | 0.00 | 0.00 |
|  |  | **min** | 0.00 | 0.00 | 0.00 | 0.00 | 0.00 | 0.00 |
|  |  | **max** | 18.00 | 42.00 | 13.00 | 19.00 | 12.00 | 36.00 |

B) **4-year follow-up**

|  |  |  | **number of total events** | **impact of total events** | **number of positively appraised events** | **impact of positively appraised events** | **number of negatively appraised events** | **impact of negatively appraised events** |
| --- | --- | --- | --- | --- | --- | --- | --- | --- |
| **entire sample** | **youth report** | **mean** | 5.28 | 8.50 | 1.46 | 2.60 | 2.41 | 4.60 |
|  |  | **std** | 3.63 | 7.22 | 1.39 | 3.10 | 2.31 | 5.10 |
|  |  | **median** | 5.00 | 7.00 | 1.00 | 2.00 | 2.00 | 3.00 |
|  |  | **min** | 0.00 | 0.00 | 0.00 | 0.00 | 0.00 | 0.00 |
|  |  | **max** | 24.00 | 50.00 | 10.00 | 20.00 | 17.00 | 40.00 |
|  | **caregiver report** | **mean** | 3.73 | 5.55 | 1.37 | 1.89 | 1.19 | 2.42 |
|  |  | **std** | 2.99 | 5.95 | 1.57 | 2.73 | 1.73 | 3.94 |
|  |  | **median** | 3.00 | 4.00 | 1.00 | 1.00 | 1.00 | 1.00 |
|  |  | **min** | 0.00 | 0.00 | 0.00 | 0.00 | 0.00 | 0.00 |
|  |  | **max** | 22.00 | 48.00 | 11.00 | 23.00 | 17.00 | 42.00 |
| **female adolescents** | **youth report** | **mean** | 5.73 | 9.44 | 1.53 | 2.74 | 2.67 | 5.23 |
|  |  | **std** | 3.80 | 7.65 | 1.42 | 3.18 | 2.44 | 5.44 |
|  |  | **median** | 5.00 | 8.00 | 1.00 | 2.00 | 2.00 | 4.00 |
|  |  | **min** | 0.00 | 0.00 | 0.00 | 0.00 | 0.00 | 0.00 |
|  |  | **max** | 24.00 | 50.00 | 7.00 | 17.00 | 15.00 | 38.00 |
|  | **caregiver report** | **mean** | 3.72 | 5.70 | 1.39 | 1.93 | 1.22 | 2.52 |
|  |  | **std** | 3.02 | 6.07 | 1.60 | 2.79 | 1.75 | 4.08 |
|  |  | **median** | 3.00 | 4.00 | 1.00 | 1.00 | 1.00 | 1.00 |
|  |  | **min** | 0.00 | 0.00 | 0.00 | 0.00 | 0.00 | 0.00 |
|  |  | **max** | 18.00 | 45.00 | 11.00 | 23.00 | 14.00 | 42.00 |
| **male adolescents** | **youth report** | **mean** | 4.89 | 7.68 | 1.40 | 2.49 | 2.20 | 4.06 |
|  |  | **std** | 3.43 | 6.72 | 1.37 | 3.03 | 2.16 | 4.72 |
|  |  | **median** | 4.00 | 6.00 | 1.00 | 1.00 | 2.00 | 3.00 |
|  |  | **min** | 0.00 | 0.00 | 0.00 | 0.00 | 0.00 | 0.00 |
|  |  | **max** | 24.00 | 49.00 | 10.00 | 20.00 | 17.00 | 40.00 |
|  | **caregiver report** | **mean** | 3.73 | 5.41 | 1.36 | 1.85 | 1.16 | 2.33 |
|  |  | **std** | 2.97 | 5.84 | 1.54 | 2.68 | 1.70 | 3.82 |
|  |  | **median** | 3.00 | 4.00 | 1.00 | 1.00 | 1.00 | 1.00 |
|  |  | **min** | 0.00 | 0.00 | 0.00 | 0.00 | 0.00 | 0.00 |
|  |  | **max** | 22.00 | 48.00 | 11.00 | 20.00 | 17.00 | 41.00 |

C) **6-year follow-up**

|  |  |  | **number of total events** | **impact of total events** | **number of positively appraised events** | **impact of positively appraised events** | **number of negatively appraised events** | **impact of negatively appraised events** |
| --- | --- | --- | --- | --- | --- | --- | --- | --- |
| **entire sample** | **youth report** | **mean** | 7.01 | 11.19 | 1.88 | 3.38 | 3.31 | 6.17 |
|  |  | **std** | 4.36 | 8.70 | 1.59 | 3.48 | 2.95 | 6.42 |
|  |  | **median** | 6.00 | 9.00 | 2.00 | 3.00 | 3.00 | 4.00 |
|  |  | **min** | 0.00 | 0.00 | 0.00 | 0.00 | 0.00 | 0.00 |
|  |  | **max** | 27.00 | 64.00 | 10.00 | 23.00 | 21.00 | 56.00 |
|  | **caregiver report** | **mean** | 4.37 | 6.43 | 1.49 | 2.02 | 1.44 | 2.93 |
|  |  | **std** | 3.50 | 6.97 | 1.71 | 2.91 | 2.05 | 4.76 |
|  |  | **median** | 4.00 | 4.00 | 1.00 | 1.00 | 1.00 | 1.00 |
|  |  | **min** | 0.00 | 0.00 | 0.00 | 0.00 | 0.00 | 0.00 |
|  |  | **max** | 23.00 | 57.00 | 18.00 | 25.00 | 19.00 | 51.00 |
| **female adolescents** | **youth report** | **mean** | 7.63 | 12.69 | 1.97 | 3.50 | 3.75 | 7.31 |
|  |  | **std** | 4.52 | 9.35 | 1.60 | 3.47 | 3.17 | 7.09 |
|  |  | **median** | 7.00 | 11.00 | 2.00 | 3.00 | 3.00 | 5.00 |
|  |  | **min** | 0.00 | 0.00 | 0.00 | 0.00 | 0.00 | 0.00 |
|  |  | **max** | 27.00 | 64.00 | 10.00 | 19.00 | 21.00 | 56.00 |
|  | **caregiver report** | **mean** | 4.47 | 6.86 | 1.50 | 2.11 | 1.58 | 3.28 |
|  |  | **std** | 3.55 | 7.35 | 1.71 | 2.98 | 2.16 | 5.14 |
|  |  | **median** | 4.00 | 5.00 | 1.00 | 1.00 | 1.00 | 2.00 |
|  |  | **min** | 0.00 | 0.00 | 0.00 | 0.00 | 0.00 | 0.00 |
|  |  | **max** | 22.00 | 57.00 | 15.00 | 25.00 | 19.00 | 51.00 |
| **male adolescents** | **youth report** | **mean** | 6.45 | 9.83 | 1.79 | 3.27 | 2.91 | 5.13 |
|  |  | **std** | 4.13 | 7.81 | 1.57 | 3.49 | 2.68 | 5.55 |
|  |  | **median** | 6.00 | 8.00 | 2.00 | 3.00 | 2.00 | 3.00 |
|  |  | **min** | 0.00 | 0.00 | 0.00 | 0.00 | 0.00 | 0.00 |
|  |  | **max** | 27.00 | 54.00 | 9.00 | 23.00 | 18.00 | 45.00 |
|  | **caregiver report** | **mean** | 4.29 | 6.04 | 1.47 | 1.93 | 1.31 | 2.62 |
|  |  | **std** | 3.45 | 6.59 | 1.71 | 2.84 | 1.94 | 4.37 |
|  |  | **median** | 3.00 | 4.00 | 1.00 | 1.00 | 1.00 | 1.00 |
|  |  | **min** | 0.00 | 0.00 | 0.00 | 0.00 | 0.00 | 0.00 |
|  |  | **max** | 23.00 | 49.00 | 18.00 | 24.00 | 17.00 | 40.00 |

**Supplementary Table 3. Comparing the number and impact of positively versus negatively appraised stressful life events**

|  |  |  | **number of negatively vs. positively perceived events** | **impact of negatively vs. positively perceived events** |
| --- | --- | --- | --- | --- |
| **A) 2-year follow-up** | **youth report** | **entire sample** | t = 26.62, p < 0.001 | t = 32.48, p < 0.001 |
|  |  | **female adolescents** | t = 16.81, p < 0.001 | t = 22.16, p < 0.001 |
|  |  | **male adolescents** | t = 20.72, p < 0.001 | t = 23.75, p < 0.001 |
|  | **caregiver report** | **entire sample** | t = -7.90, p < 0.001 | t = 9.11, p < 0.001 |
|  |  | **female adolescents** | t = -5.37, p < 0.001 | t = 6.23, p < 0.001 |
|  |  | **male adolescents** | t = -5.81, p < 0.001 | t = 6.65, p < 0.001 |
| **B) 4-year follow-up** | **youth report** | **entire sample** | t = 25.05, p < 0.001 | t = 23.75, p < 0.001 |
|  |  | **female adolescents** | t = 19.37, p < 0.001 | t = 19.09, p < 0.001 |
|  |  | **male adolescents** | t = 16.17, p < 0.001 | t = 14.56, p < 0.001 |
|  | **caregiver report** | **entire sample** | t = -5.65, p < 0.001 | t = 7.80, p < 0.001 |
|  |  | **female adolescents** | t = -3.53, 0 = 0.0004 | t = 5.75, p < 0.001 |
|  |  | **male adolescents** | t = -4.44, p < 0.001 | t = 5.29, p < 0.001 |
| **C) 6-year follow-up** | **youth report** | **entire sample** | t = 33.02, p < 0.001 | t = 29.45, p < 0.001 |
|  |  | **female adolescents** | t = 26.78, p < 0.001 | t = 25.65, p < 0.001 |
|  |  | **male adolescents** | t = 20.07, p < 0.001 | t = 15.90, p < 0.001 |
|  | **caregiver report** | **entire sample** | t = -1.34, p = 0.1792 | t = 12.70, p < 0.001 |
|  |  | **female adolescents** | t = 1.46, p = 0.1434 | t = 10.41, p < 0.001 |
|  |  | **male adolescents** | t = -3.41, p = 0.0006 | t = 7.47, p < 0.001 |

T-tests contrasting positively versus negatively appraised number and impact of stressful life events for the entire sample, female adolescents and male adolescents for the (A) 2-year follow-up, (B) 4-year follow-up, and (C) 6-year follow-up time point for the youth report and caregiver report.

**Supplementary Table 4. Individual items adverse life events questionnaire**

**Table S4.1. Youth report – 2 year follow-up**

| **Item** | **Code** | **2 year follow-up** | | | | | | | | | | | |
| --- | --- | --- | --- | --- | --- | --- | --- | --- | --- | --- | --- | --- | --- |
|  |  | **Count (#, %)** | | | | **Experience (#, % of people who have count==1)** | | | | **Severity (#, % of people who have count==1)** | | | |
|  |  | **1** | **0** | **999** | **777** | **1** | **2** | **999** | **444** | **0** | **1** | **2** | **3** |
|  |  | **Yes** | **No** | **Dont know** | **Decline to answer** | **Mostly good** | **Mostly bad** | **Dont know** | **Not applicable** | **Not at all** | **A little** | **Some** | **A lot** |
| **SLE questions** | | | | | | | | | | | | | |
| In your life time - Someone in family died? | 001 | 4943 (77.65%) | 1418 (22.27%) | 0 (0.00%) | 0 (0.00%) | 82 (1.66%) | 3526 (71.33%) | 960 (19.42%) | 375 (7.59%) | 541 (10.94%) | 1564 (31.64%) | 1323 (26.77%) | 1515 (30.65%) |
| In your life time - Family member was seriously injured? [Youth] | 002 | 2418 (37.98%) | 3943 (61.94%) | 0 (0.00%) | 0 (0.00%) | 59 (2.44%) | 1801 (74.48%) | 376 (15.55%) | 183 (7.57%) | 332 (13.73%) | 786 (32.51%) | 633 (26.18%) | 668 (27.63%) |
| In your life time - Saw crime or accident? [Youth] | 003 | 1821 (28.61%) | 4540 (71.32%) | 0 (0.00%) | 0 (0.00%) | 57 (3.13%) | 978 (53.71%) | 487 (26.74%) | 299 (16.42%) | 944 (51.84%) | 517 (28.39%) | 206 (11.31%) | 154 (8.46%) |
| In your life time - Close friend died? [Youth] | 004 | 240 (3.77%) | 6121 (96.15%) | 0 (0.00%) | 0 (0.00%) | 2 (0.83%) | 224 (93.33%) | 7 (2.92%) | 8 (3.33%) | 4 (1.67%) | 27 (11.25%) | 67 (27.92%) | 143 (59.58%) |
| In your life time - Lost a close friend? [Youth] | 005 | 1331 (20.91%) | 5030 (79.01%) | 0 (0.00%) | 0 (0.00%) | 91 (6.84%) | 1019 (76.56%) | 147 (11.04%) | 74 (5.56%) | 89 (6.69%) | 322 (24.19%) | 354 (26.60%) | 566 (42.52%) |
| In your life time - Close friend was seriously sick/injured? [Youth] | 006 | 878 (13.79%) | 5483 (86.13%) | 0 (0.00%) | 0 (0.00%) | 24 (2.73%) | 680 (77.45%) | 105 (11.96%) | 69 (7.86%) | 109 (12.41%) | 273 (31.09%) | 258 (29.38%) | 238 (27.11%) |
| In your life time - Negative change in parent's financial situation? [Youth] | 007 | 547 (8.59%) | 5814 (91.33%) | 0 (0.00%) | 0 (0.00%) | 11 (2.01%) | 414 (75.69%) | 86 (15.72%) | 36 (6.58%) | 46 (8.41%) | 162 (29.62%) | 174 (31.81%) | 165 (30.16%) |
| In your life time - Family member had drug and/or alcohol problem? [Youth] | 008 | 822 (12.91%) | 5538 (86.99%) | 0 (0.00%) | 0 (0.00%) | 17 (2.07%) | 551 (67.03%) | 148 (18.00%) | 106 (12.90%) | 195 (23.72%) | 215 (26.16%) | 163 (19.83%) | 249 (30.29%) |
| In your life time - You got seriously sick? [Youth] | 009 | 887 (13.93%) | 5474 (85.99%) | 0 (0.00%) | 0 (0.00%) | 21 (2.37%) | 763 (86.02%) | 74 (8.34%) | 29 (3.27%) | 66 (7.44%) | 176 (19.84%) | 194 (21.87%) | 451 (50.85%) |
| In your life time - You got seriously injured? [Youth] | 010 | 1015 (15.94%) | 5346 (83.98%) | 0 (0.00%) | 0 (0.00%) | 39 (3.84%) | 853 (84.04%) | 79 (7.78%) | 44 (4.33%) | 81 (7.98%) | 207 (20.39%) | 205 (20.20%) | 522 (51.43%) |
| In your life time - Parents argued more than previously? [Youth] | 011 | 523 (8.22%) | 5838 (91.71%) | 0 (0.00%) | 0 (0.00%) | 12 (2.29%) | 405 (77.44%) | 66 (12.62%) | 40 (7.65%) | 60 (11.47%) | 153 (29.25%) | 125 (23.90%) | 185 (35.37%) |
| In your life time - Parental figure lost job? [Youth] | 012 | 642 (10.08%) | 5719 (89.84%) | 0 (0.00%) | 0 (0.00%) | 56 (8.72%) | 331 (51.56%) | 158 (24.61%) | 97 (15.11%) | 146 (22.74%) | 242 (37.69%) | 148 (23.05%) | 106 (16.51%) |
| In your life time - One parent was away from home more often? [Youth] | 013 | 1192 (18.72%) | 5169 (81.20%) | 0 (0.00%) | 0 (0.00%) | 173 (14.51%) | 494 (41.44%) | 345 (28.94%) | 180 (15.10%) | 312 (26.17%) | 390 (32.72%) | 245 (20.55%) | 245 (20.55%) |
| In your life time - Someone in the family was arrested? [Youth] | 014 | 787 (12.36%) | 5574 (87.56%) | 0 (0.00%) | 0 (0.00%) | 31 (3.94%) | 458 (58.20%) | 186 (23.63%) | 112 (14.23%) | 227 (28.84%) | 190 (24.14%) | 137 (17.41%) | 233 (29.61%) |
| In your life time - Family member had mental/emotional problem? [Youth] | 015 | 694 (10.90%) | 5667 (89.02%) | 0 (0.00%) | 0 (0.00%) | 19 (2.74%) | 457 (65.85%) | 129 (18.59%) | 89 (12.82%) | 112 (16.14%) | 207 (29.83%) | 172 (24.78%) | 203 (29.25%) |
| In your life time - Sibling left home? [Youth] | 016 | 720 (11.31%) | 5641 (88.61%) | 0 (0.00%) | 0 (0.00%) | 232 (32.22%) | 244 (33.89%) | 158 (21.94%) | 86 (11.94%) | 149 (20.69%) | 226 (31.39%) | 157 (21.81%) | 188 (26.11%) |
| In your life time - Was a victim of crime/violence/assault? [Youth] | 017 | 90 (1.41%) | 6271 (98.51%) | 0 (0.00%) | 0 (0.00%) | 3 (3.33%) | 80 (88.89%) | 7 (7.78%) | 0 (0.00%) | 9 (10.00%) | 11 (12.22%) | 17 (18.89%) | 53 (58.89%) |
| In your life time - Parents separated or divorced? [Youth] | 018 | 1479 (23.23%) | 4882 (76.69%) | 0 (0.00%) | 0 (0.00%) | 124 (8.38%) | 804 (54.36%) | 409 (27.65%) | 142 (9.60%) | 264 (17.85%) | 295 (19.95%) | 257 (17.38%) | 663 (44.83%) |
| In your life time - Parents/caregiver got into trouble with the law? [Youth] | 019 | 327 (5.14%) | 6034 (94.78%) | 0 (0.00%) | 0 (0.00%) | 18 (5.50%) | 190 (58.10%) | 77 (23.55%) | 42 (12.84%) | 101 (30.89%) | 75 (22.94%) | 57 (17.43%) | 94 (28.75%) |
| In your life time - Attended a new school? [Youth] | 020 | 3554 (55.83%) | 2807 (44.09%) | 0 (0.00%) | 0 (0.00%) | 2539 (71.44%) | 381 (10.72%) | 444 (12.49%) | 190 (5.35%) | 842 (23.69%) | 809 (22.76%) | 657 (18.49%) | 1246 (35.06%) |
| In your life time - Family moved? [Youth] | 021 | 3033 (47.64%) | 3328 (52.28%) | 0 (0.00%) | 0 (0.00%) | 1938 (63.90%) | 309 (10.19%) | 580 (19.12%) | 206 (6.79%) | 890 (29.34%) | 620 (20.44%) | 555 (18.30%) | 968 (31.92%) |
| In your life time - One of the parents/caregivers went to jail? [Youth] | 022 | 416 (6.53%) | 5945 (93.39%) | 0 (0.00%) | 0 (0.00%) | 16 (3.85%) | 261 (62.74%) | 106 (25.48%) | 33 (7.93%) | 100 (24.04%) | 83 (19.95%) | 83 (19.95%) | 150 (36.06%) |
| In your life time - Got new step-parent? [Youth] | 023 | 832 (13.07%) | 5529 (86.85%) | 0 (0.00%) | 0 (0.00%) | 530 (63.70%) | 95 (11.42%) | 140 (16.83%) | 67 (8.05%) | 218 (26.20%) | 173 (20.79%) | 158 (18.99%) | 283 (34.01%) |
| In your life time - Parent/caregiver got a new job? [Youth] | 024 | 2433 (38.22%) | 3928 (61.70%) | 0 (0.00%) | 0 (0.00%) | 1880 (77.27%) | 49 (2.01%) | 319 (13.11%) | 185 (7.60%) | 1071 (44.02%) | 635 (26.10%) | 397 (16.32%) | 330 (13.56%) |
| In your life time - Got new sibling? [Youth] | 025 | 2482 (38.99%) | 3879 (60.93%) | 0 (0.00%) | 0 (0.00%) | 1982 (79.85%) | 136 (5.48%) | 276 (11.12%) | 88 (3.55%) | 404 (16.28%) | 310 (12.49%) | 280 (11.28%) | 1488 (59.95%) |

**Table S4.2. Youth report – 4 year follow-up**

| **Item** | **Code** | **4 year follow-up** | | | | | | | | | | | |
| --- | --- | --- | --- | --- | --- | --- | --- | --- | --- | --- | --- | --- | --- |
|  |  | **Count (#, %)** | | | | **Experience (#, % of people who have count==1)** | | | | **Severity (#, % of people who have count==1)** | | | |
|  |  | **1** | **0** | **999** | **777** | **1** | **2** | **999** | **444** | **0** | **1** | **2** | **3** |
|  |  | **Yes** | **No** | **Dont know** | **Decline to answer** | **Mostly good** | **Mostly bad** | **Dont know** | **Not applicable** | **Not at all** | **A little** | **Some** | **A lot** |
| **SLE questions** | | | | | | | | | | | | | |
| In your life time - Someone in family died? | 001 | 3718 (73.84%) | 1303 (25.88%) | 0 (0.00%) | 0 (0.00%) | 77 (2.07%) | 2650 (71.27%) | 680 (18.29%) | 311 (8.36%) | 466 (12.53%) | 1307 (35.15%) | 1028 (27.65%) | 917 (24.66%) |
| In your life time - Family member was seriously injured? [Youth] | 002 | 1572 (31.22%) | 3449 (68.50%) | 0 (0.00%) | 0 (0.00%) | 34 (2.16%) | 1196 (76.08%) | 204 (12.98%) | 138 (8.78%) | 217 (13.80%) | 594 (37.79%) | 471 (29.96%) | 290 (18.45%) |
| In your life time - Saw crime or accident? [Youth] | 003 | 985 (19.56%) | 4036 (80.16%) | 0 (0.00%) | 0 (0.00%) | 32 (3.25%) | 517 (52.49%) | 256 (25.99%) | 180 (18.27%) | 501 (50.86%) | 310 (31.47%) | 115 (11.68%) | 59 (5.99%) |
| In your life time - Close friend died? [Youth] | 004 | 217 (4.31%) | 4804 (95.41%) | 0 (0.00%) | 0 (0.00%) | 2 (0.92%) | 203 (93.55%) | 5 (2.30%) | 7 (3.23%) | 3 (1.38%) | 19 (8.76%) | 53 (24.42%) | 142 (65.44%) |
| In your life time - Lost a close friend? [Youth] | 005 | 899 (17.86%) | 4122 (81.87%) | 0 (0.00%) | 0 (0.00%) | 88 (9.79%) | 660 (73.41%) | 105 (11.68%) | 46 (5.12%) | 41 (4.56%) | 201 (22.36%) | 269 (29.92%) | 388 (43.16%) |
| In your life time - Close friend was seriously sick/injured? [Youth] | 006 | 427 (8.48%) | 4594 (91.24%) | 0 (0.00%) | 0 (0.00%) | 8 (1.87%) | 340 (79.63%) | 46 (10.77%) | 33 (7.73%) | 48 (11.24%) | 141 (33.02%) | 145 (33.96%) | 93 (21.78%) |
| In your life time - Negative change in parent's financial situation? [Youth] | 007 | 475 (9.43%) | 4546 (90.29%) | 0 (0.00%) | 0 (0.00%) | 12 (2.53%) | 361 (76.00%) | 64 (13.47%) | 38 (8.00%) | 38 (8.00%) | 148 (31.16%) | 171 (36.00%) | 118 (24.84%) |
| In your life time - Family member had drug and/or alcohol problem? [Youth] | 008 | 622 (12.35%) | 4399 (87.37%) | 0 (0.00%) | 0 (0.00%) | 10 (1.61%) | 461 (74.12%) | 89 (14.31%) | 62 (9.97%) | 113 (18.17%) | 194 (31.19%) | 129 (20.74%) | 186 (29.90%) |
| In your life time - You got seriously sick? [Youth] | 009 | 449 (8.92%) | 4572 (90.80%) | 0 (0.00%) | 0 (0.00%) | 15 (3.34%) | 382 (85.08%) | 33 (7.35%) | 19 (4.23%) | 45 (10.02%) | 75 (16.70%) | 137 (30.51%) | 192 (42.76%) |
| In your life time - You got seriously injured? [Youth] | 010 | 533 (10.59%) | 4488 (89.14%) | 0 (0.00%) | 0 (0.00%) | 31 (5.82%) | 448 (84.05%) | 36 (6.75%) | 17 (3.19%) | 32 (6.00%) | 116 (21.76%) | 139 (26.08%) | 245 (45.97%) |
| In your life time - Parents argued more than previously? [Youth] | 011 | 519 (10.31%) | 4502 (89.41%) | 0 (0.00%) | 0 (0.00%) | 6 (1.16%) | 427 (82.27%) | 56 (10.79%) | 30 (5.78%) | 34 (6.55%) | 134 (25.82%) | 166 (31.98%) | 185 (35.65%) |
| In your life time - Parental figure lost job? [Youth] | 012 | 552 (10.96%) | 4469 (88.76%) | 0 (0.00%) | 0 (0.00%) | 56 (10.14%) | 293 (53.08%) | 127 (23.01%) | 76 (13.77%) | 128 (23.19%) | 194 (35.14%) | 164 (29.71%) | 66 (11.96%) |
| In your life time - One parent was away from home more often? [Youth] | 013 | 693 (13.76%) | 4328 (85.96%) | 0 (0.00%) | 0 (0.00%) | 119 (17.17%) | 270 (38.96%) | 192 (27.71%) | 112 (16.16%) | 118 (17.03%) | 231 (33.33%) | 184 (26.55%) | 160 (23.09%) |
| In your life time - Someone in the family was arrested? [Youth] | 014 | 460 (9.14%) | 4561 (90.59%) | 0 (0.00%) | 0 (0.00%) | 20 (4.35%) | 255 (55.43%) | 113 (24.57%) | 72 (15.65%) | 148 (32.17%) | 148 (32.17%) | 62 (13.48%) | 102 (22.17%) |
| In your life time - Family member had mental/emotional problem? [Youth] | 015 | 670 (13.31%) | 4351 (86.42%) | 0 (0.00%) | 0 (0.00%) | 8 (1.19%) | 510 (76.12%) | 105 (15.67%) | 47 (7.01%) | 83 (12.39%) | 220 (32.84%) | 195 (29.10%) | 172 (25.67%) |
| In your life time - Sibling left home? [Youth] | 016 | 703 (13.96%) | 4318 (85.76%) | 0 (0.00%) | 0 (0.00%) | 282 (40.11%) | 159 (22.62%) | 151 (21.48%) | 111 (15.79%) | 125 (17.78%) | 237 (33.71%) | 207 (29.45%) | 134 (19.06%) |
| In your life time - Was a victim of crime/violence/assault? [Youth] | 017 | 141 (2.80%) | 4880 (96.92%) | 0 (0.00%) | 0 (0.00%) | 3 (2.13%) | 125 (88.65%) | 7 (4.96%) | 6 (4.26%) | 9 (6.38%) | 25 (17.73%) | 32 (22.70%) | 75 (53.19%) |
| In your life time - Parents separated or divorced? [Youth] | 018 | 1138 (22.60%) | 3883 (77.12%) | 0 (0.00%) | 0 (0.00%) | 169 (14.85%) | 529 (46.49%) | 312 (27.42%) | 128 (11.25%) | 206 (18.10%) | 203 (17.84%) | 219 (19.24%) | 510 (44.82%) |
| In your life time - Parents/caregiver got into trouble with the law? [Youth] | 019 | 199 (3.95%) | 4822 (95.77%) | 0 (0.00%) | 0 (0.00%) | 6 (3.02%) | 110 (55.28%) | 55 (27.64%) | 28 (14.07%) | 64 (32.16%) | 47 (23.62%) | 32 (16.08%) | 56 (28.14%) |
| In your life time - Attended a new school? [Youth] | 020 | 2703 (53.68%) | 2318 (46.04%) | 0 (0.00%) | 0 (0.00%) | 2032 (75.18%) | 202 (7.47%) | 307 (11.36%) | 162 (5.99%) | 396 (14.65%) | 543 (20.09%) | 672 (24.86%) | 1092 (40.40%) |
| In your life time - Family moved? [Youth] | 021 | 1918 (38.09%) | 3103 (61.63%) | 0 (0.00%) | 0 (0.00%) | 1369 (71.38%) | 144 (7.51%) | 265 (13.82%) | 140 (7.30%) | 345 (17.99%) | 364 (18.98%) | 408 (21.27%) | 801 (41.76%) |
| In your life time - One of the parents/caregivers went to jail? [Youth] | 022 | 212 (4.21%) | 4809 (95.51%) | 0 (0.00%) | 0 (0.00%) | 11 (5.19%) | 121 (57.08%) | 52 (24.53%) | 28 (13.21%) | 53 (25.00%) | 56 (26.42%) | 41 (19.34%) | 62 (29.25%) |
| In your life time - Got new step-parent? [Youth] | 023 | 636 (12.63%) | 4385 (87.09%) | 0 (0.00%) | 0 (0.00%) | 425 (66.82%) | 71 (11.16%) | 108 (16.98%) | 32 (5.03%) | 116 (18.24%) | 120 (18.87%) | 143 (22.48%) | 257 (40.41%) |
| In your life time - Parent/caregiver got a new job? [Youth] | 024 | 1566 (31.10%) | 3455 (68.62%) | 0 (0.00%) | 0 (0.00%) | 1251 (79.89%) | 31 (1.98%) | 164 (10.47%) | 120 (7.66%) | 571 (36.46%) | 524 (33.46%) | 311 (19.86%) | 160 (10.22%) |
| In your life time - Got new sibling? [Youth] | 025 | 1390 (27.61%) | 3631 (72.12%) | 0 (0.00%) | 0 (0.00%) | 1137 (81.80%) | 48 (3.45%) | 139 (10.00%) | 66 (4.75%) | 130 (9.35%) | 138 (9.93%) | 202 (14.53%) | 920 (66.19%) |
| In your life time - Parent or caregiver deported? [Youth] | 026 | 20 (0.40%) | 4915 (97.62%) | 0 (0.00%) | 0 (0.00%) | 3 (15.00%) | 13 (65.00%) | 2 (10.00%) | 2 (10.00%) | 5 (25.00%) | 3 (15.00%) | 4 (20.00%) | 8 (40.00%) |
| In your life time - Parent or caregiver hospitalized? [Youth] | 027 | 692 (13.74%) | 4243 (84.27%) | 0 (0.00%) | 0 (0.00%) | 46 (6.65%) | 495 (71.53%) | 90 (13.01%) | 61 (8.82%) | 78 (11.27%) | 214 (30.92%) | 211 (30.49%) | 189 (27.31%) |
| In your life time - You were placed in foster care? [Youth] | 028 | 43 (0.85%) | 4892 (97.16%) | 0 (0.00%) | 0 (0.00%) | 6 (13.95%) | 19 (44.19%) | 13 (30.23%) | 5 (11.63%) | 3 (6.98%) | 11 (25.58%) | 5 (11.63%) | 24 (55.81%) |
| In your life time - Saw or heard someone getting hit, slapped, punched, or beaten up in your school or neighborhood? [Youth] | 029 | 1001 (19.88%) | 3934 (78.13%) | 0 (0.00%) | 0 (0.00%) | 46 (4.60%) | 353 (35.26%) | 335 (33.47%) | 267 (26.67%) | 694 (69.33%) | 223 (22.28%) | 56 (5.59%) | 28 (2.80%) |
| In your life time - Saw or heard someone being shot at (but not actually wounded) in your school or neighborhood? [Youth] | 030 | 231 (4.59%) | 4704 (93.43%) | 0 (0.00%) | 0 (0.00%) | 1 (0.43%) | 108 (46.75%) | 72 (31.17%) | 50 (21.65%) | 125 (54.11%) | 70 (30.30%) | 17 (7.36%) | 19 (8.23%) |
| In your life time - Had a lockdown at your school due to concerns about a school shooting or violence? [Youth] | 031 | 1126 (22.36%) | 3809 (75.65%) | 0 (0.00%) | 0 (0.00%) | 60 (5.33%) | 648 (57.55%) | 234 (20.78%) | 184 (16.34%) | 452 (40.14%) | 414 (36.77%) | 182 (16.16%) | 78 (6.93%) |
| In your life time - Your family was without a home or place to stay/live? [Youth] | 032 | 76 (1.51%) | 4945 (98.21%) | 0 (0.00%) | 0 (0.00%) | 0 (0.00%) | 10 (13.16%) | 3 (3.95%) | 3 (3.95%) | 5 (6.58%) | 1 (1.32%) | 1 (1.32%) | 9 (11.84%) |
| In your life time - Do you know anyone who has attempted suicide? [Youth] | 034 | 1110 (22.05%) | 3911 (77.68%) | 0 (0.00%) | 0 (0.00%) |  |  |  |  | 58 (5.23%) | 152 (13.69%) | 154 (13.87%) | 204 (18.38%) |

**Table S4.3. Youth report – 6 year follow-up**

| **Item** | **Code** | **6 year follow-up** | | | | | | | | | | | |
| --- | --- | --- | --- | --- | --- | --- | --- | --- | --- | --- | --- | --- | --- |
|  |  | **Count (#, %)** | | | | **Experience (#, % of people who have count==1)** | | | | **Severity (#, % of people who have count==1)** | | | |
|  |  | **1** | **0** | **999** | **777** | **1** | **2** | **999** | **444** | **0** | **1** | **2** | **3** |
|  |  | **Yes** | **No** | **Dont know** | **Decline to answer** | **Mostly good** | **Mostly bad** | **Dont know** | **Not applicable** | **Not at all** | **A little** | **Some** | **A lot** |
| **SLE questions** | | | | | | | | | | | | | |
| In your life time - Someone in family died? | 001 | 4675 (78.53%) | 1259 (21.15%) | 0 (0.00%) | 1 (0.02%) | 157 (3.36%) | 3372 (72.13%) | 720 (15.40%) | 426 (9.11%) | 613 (13.11%) | 1807 (38.65%) | 1284 (27.47%) | 971 (20.77%) |
| In your life time - Family member was seriously injured? [Youth] | 002 | 2089 (35.09%) | 3842 (64.54%) | 4 (0.07%) | 0 (0.00%) | 64 (3.06%) | 1605 (76.83%) | 252 (12.06%) | 168 (8.04%) | 321 (15.37%) | 802 (38.39%) | 605 (28.96%) | 361 (17.28%) |
| In your life time - Saw crime or accident? [Youth] | 003 | 1662 (27.92%) | 4271 (71.75%) | 2 (0.03%) | 0 (0.00%) | 62 (3.73%) | 843 (50.72%) | 379 (22.80%) | 378 (22.74%) | 851 (51.20%) | 491 (29.54%) | 206 (12.39%) | 114 (6.86%) |
| In your life time - Close friend died? [Youth] | 004 | 470 (7.90%) | 5465 (91.80%) | 0 (0.00%) | 0 (0.00%) | 10 (2.13%) | 429 (91.28%) | 22 (4.68%) | 9 (1.91%) | 6 (1.28%) | 73 (15.53%) | 141 (30.00%) | 250 (53.19%) |
| In your life time - Lost a close friend? [Youth] | 005 | 1570 (26.37%) | 4364 (73.31%) | 0 (0.00%) | 1 (0.02%) | 171 (10.89%) | 1156 (73.63%) | 155 (9.87%) | 88 (5.61%) | 66 (4.20%) | 373 (23.76%) | 491 (31.27%) | 640 (40.76%) |
| In your life time - Close friend was seriously sick/injured? [Youth] | 006 | 493 (8.28%) | 5441 (91.40%) | 0 (0.00%) | 1 (0.02%) | 16 (3.25%) | 403 (81.74%) | 39 (7.91%) | 35 (7.10%) | 54 (10.95%) | 172 (34.89%) | 176 (35.70%) | 91 (18.46%) |
| In your life time - Negative change in parent's financial situation? [Youth] | 007 | 1143 (19.20%) | 4789 (80.45%) | 3 (0.05%) | 0 (0.00%) | 41 (3.59%) | 878 (76.82%) | 142 (12.42%) | 82 (7.17%) | 53 (4.64%) | 349 (30.53%) | 440 (38.50%) | 301 (26.33%) |
| In your life time - Family member had drug and/or alcohol problem? [Youth] | 008 | 1178 (19.79%) | 4753 (79.84%) | 3 (0.05%) | 1 (0.02%) | 17 (1.44%) | 870 (73.85%) | 153 (12.99%) | 138 (11.71%) | 232 (19.69%) | 350 (29.71%) | 249 (21.14%) | 347 (29.46%) |
| In your life time - You got seriously sick? [Youth] | 009 | 693 (11.64%) | 5240 (88.02%) | 2 (0.03%) | 0 (0.00%) | 21 (3.03%) | 585 (84.42%) | 57 (8.23%) | 30 (4.33%) | 67 (9.67%) | 156 (22.51%) | 190 (27.42%) | 280 (40.40%) |
| In your life time - You got seriously injured? [Youth] | 010 | 767 (12.88%) | 5167 (86.80%) | 0 (0.00%) | 0 (0.00%) | 45 (5.87%) | 648 (84.49%) | 48 (6.26%) | 26 (3.39%) | 58 (7.56%) | 160 (20.86%) | 223 (29.07%) | 326 (42.50%) |
| In your life time - Parents argued more than previously? [Youth] | 011 | 1032 (17.34%) | 4902 (82.35%) | 1 (0.02%) | 0 (0.00%) | 13 (1.26%) | 896 (86.82%) | 86 (8.33%) | 37 (3.59%) | 42 (4.07%) | 208 (20.16%) | 384 (37.21%) | 398 (38.57%) |
| In your life time - Parental figure lost job? [Youth] | 012 | 1006 (16.90%) | 4928 (82.78%) | 0 (0.00%) | 1 (0.02%) | 101 (10.04%) | 548 (54.47%) | 229 (22.76%) | 128 (12.72%) | 219 (21.77%) | 381 (37.87%) | 257 (25.55%) | 149 (14.81%) |
| In your life time - One parent was away from home more often? [Youth] | 013 | 1192 (20.02%) | 4741 (79.64%) | 0 (0.00%) | 2 (0.03%) | 157 (13.17%) | 519 (43.54%) | 281 (23.57%) | 235 (19.71%) | 197 (16.53%) | 411 (34.48%) | 323 (27.10%) | 261 (21.90%) |
| In your life time - Someone in the family was arrested? [Youth] | 014 | 775 (13.02%) | 5158 (86.65%) | 1 (0.02%) | 1 (0.02%) | 38 (4.90%) | 439 (56.65%) | 161 (20.77%) | 137 (17.68%) | 251 (32.39%) | 229 (29.55%) | 136 (17.55%) | 159 (20.52%) |
| In your life time - Family member had mental/emotional problem? [Youth] | 015 | 1283 (21.55%) | 4649 (78.10%) | 1 (0.02%) | 2 (0.03%) | 20 (1.56%) | 966 (75.29%) | 184 (14.34%) | 113 (8.81%) | 177 (13.80%) | 402 (31.33%) | 369 (28.76%) | 335 (26.11%) |
| In your life time - Sibling left home? [Youth] | 016 | 1454 (24.42%) | 4480 (75.26%) | 0 (0.00%) | 1 (0.02%) | 616 (42.37%) | 362 (24.90%) | 243 (16.71%) | 233 (16.02%) | 225 (15.47%) | 486 (33.43%) | 449 (30.88%) | 294 (20.22%) |
| In your life time - Was a victim of crime/violence/assault? [Youth] | 017 | 294 (4.94%) | 5640 (94.74%) | 0 (0.00%) | 1 (0.02%) | 4 (1.36%) | 262 (89.12%) | 13 (4.42%) | 15 (5.10%) | 20 (6.80%) | 45 (15.31%) | 82 (27.89%) | 147 (50.00%) |
| In your life time - Parents separated or divorced? [Youth] | 018 | 1515 (25.45%) | 4418 (74.21%) | 0 (0.00%) | 2 (0.03%) | 290 (19.14%) | 690 (45.54%) | 328 (21.65%) | 207 (13.66%) | 227 (14.98%) | 252 (16.63%) | 309 (20.40%) | 727 (47.99%) |
| In your life time - Parents/caregiver got into trouble with the law? [Youth] | 019 | 337 (5.66%) | 5596 (94.00%) | 2 (0.03%) | 0 (0.00%) | 14 (4.15%) | 208 (61.72%) | 71 (21.07%) | 44 (13.06%) | 94 (27.89%) | 84 (24.93%) | 76 (22.55%) | 83 (24.63%) |
| In your life time - Attended a new school? [Youth] | 020 | 3612 (60.68%) | 2322 (39.01%) | 0 (0.00%) | 1 (0.02%) | 2732 (75.64%) | 294 (8.14%) | 394 (10.91%) | 192 (5.32%) | 428 (11.85%) | 655 (18.13%) | 868 (24.03%) | 1661 (45.99%) |
| In your life time - Family moved? [Youth] | 021 | 2853 (47.93%) | 3081 (51.76%) | 0 (0.00%) | 1 (0.02%) | 2003 (70.21%) | 235 (8.24%) | 397 (13.92%) | 218 (7.64%) | 490 (17.17%) | 532 (18.65%) | 663 (23.24%) | 1168 (40.94%) |
| In your life time - One of the parents/caregivers went to jail? [Youth] | 022 | 335 (5.63%) | 5600 (94.07%) | 0 (0.00%) | 0 (0.00%) | 12 (3.58%) | 186 (55.52%) | 89 (26.57%) | 48 (14.33%) | 91 (27.16%) | 80 (23.88%) | 58 (17.31%) | 106 (31.64%) |
| In your life time - Got new step-parent? [Youth] | 023 | 929 (15.61%) | 5006 (84.09%) | 0 (0.00%) | 0 (0.00%) | 590 (63.51%) | 137 (14.75%) | 133 (14.32%) | 69 (7.43%) | 121 (13.02%) | 167 (17.98%) | 269 (28.96%) | 372 (40.04%) |
| In your life time - Parent/caregiver got a new job? [Youth] | 024 | 2382 (40.01%) | 3552 (59.67%) | 0 (0.00%) | 1 (0.02%) | 1921 (80.65%) | 44 (1.85%) | 235 (9.87%) | 182 (7.64%) | 847 (35.56%) | 851 (35.73%) | 484 (20.32%) | 200 (8.40%) |
| In your life time - Got new sibling? [Youth] | 025 | 2120 (35.61%) | 3815 (64.09%) | 0 (0.00%) | 0 (0.00%) | 1767 (83.35%) | 64 (3.02%) | 199 (9.39%) | 90 (4.25%) | 151 (7.12%) | 211 (9.95%) | 361 (17.03%) | 1397 (65.90%) |
| In your life time - Parent or caregiver deported? [Youth] | 026 | 33 (0.55%) | 5894 (99.01%) | 0 (0.00%) | 0 (0.00%) | 3 (9.09%) | 18 (54.55%) | 8 (24.24%) | 4 (12.12%) | 5 (15.15%) | 8 (24.24%) | 7 (21.21%) | 13 (39.39%) |
| In your life time - Parent or caregiver hospitalized? [Youth] | 027 | 1078 (18.11%) | 4847 (81.42%) | 2 (0.03%) | 0 (0.00%) | 60 (5.57%) | 800 (74.21%) | 131 (12.15%) | 87 (8.07%) | 122 (11.32%) | 352 (32.65%) | 334 (30.98%) | 270 (25.05%) |
| In your life time - You were placed in foster care? [Youth] | 028 | 63 (1.06%) | 5864 (98.50%) | 0 (0.00%) | 0 (0.00%) | 12 (19.05%) | 21 (33.33%) | 19 (30.16%) | 11 (17.46%) | 9 (14.29%) | 6 (9.52%) | 6 (9.52%) | 42 (66.67%) |
| In your life time - Saw or heard someone getting hit, slapped, punched, or beaten up in your school or neighborhood? [Youth] | 029 | 1669 (28.04%) | 4257 (71.51%) | 0 (0.00%) | 1 (0.02%) | 88 (5.27%) | 505 (30.26%) | 516 (30.92%) | 560 (33.55%) | 1261 (75.55%) | 301 (18.03%) | 69 (4.13%) | 38 (2.28%) |
| In your life time - Saw or heard someone being shot at (but not actually wounded) in your school or neighborhood? [Youth] | 030 | 452 (7.59%) | 5474 (91.95%) | 0 (0.00%) | 1 (0.02%) | 6 (1.33%) | 190 (42.04%) | 141 (31.19%) | 115 (25.44%) | 249 (55.09%) | 127 (28.10%) | 53 (11.73%) | 23 (5.09%) |
| In your life time - Had a lockdown at your school due to concerns about a school shooting or violence? [Youth] | 031 | 2349 (39.46%) | 3577 (60.09%) | 0 (0.00%) | 1 (0.02%) | 105 (4.47%) | 1360 (57.90%) | 459 (19.54%) | 425 (18.09%) | 945 (40.23%) | 968 (41.21%) | 330 (14.05%) | 106 (4.51%) |
| In your life time - Your family was without a home or place to stay/live? [Youth] | 032 | 156 (2.62%) | 5779 (97.08%) | 0 (0.00%) | 0 (0.00%) | 8 (5.13%) | 113 (72.44%) | 21 (13.46%) | 14 (8.97%) | 19 (12.18%) | 40 (25.64%) | 30 (19.23%) | 67 (42.95%) |
| In your life time - You were thrown out of your house? [Youth] | 033 | 90 (1.51%) | 5836 (98.03%) | 0 (0.00%) | 1 (0.02%) | 8 (8.89%) | 68 (75.56%) | 7 (7.78%) | 7 (7.78%) | 12 (13.33%) | 18 (20.00%) | 23 (25.56%) | 37 (41.11%) |
| In your life time - Do you know anyone who has attempted suicide? [Youth] | 034 | 1965 (33.01%) | 3957 (66.47%) | 4 (0.07%) | 1 (0.02%) |  |  |  |  | 50 (2.54%) | 161 (8.19%) | 205 (10.43%) | 228 (11.60%) |

**Table S4.4. Caregiver report – 2 year follow-up**

| **Item** | **Code** | **2 year follow-up** | | | | | | | | | | | |
| --- | --- | --- | --- | --- | --- | --- | --- | --- | --- | --- | --- | --- | --- |
|  |  | **Count (#, %)** | | | | **Experience (#, % of people who have count==1)** | | | | **Severity (#, % of people who have count==1)** | | | |
|  |  | **1** | **0** | **999** | **777** | **1** | **2** | **999** | **444** | **0** | **1** | **2** | **3** |
|  |  | **Yes** | **No** | **Dont know** | **Decline to answer** | **Mostly good** | **Mostly bad** | **Dont know** | **Not applicable** | **Not at all** | **A little** | **Some** | **A lot** |
| **SLE questions** | | | | | | | | | | | | | |
| Someone in family died? [Parent] | 001 | 4019 (63.13%) | 2347 (36.87%) | 0 (0.00%) | 0 (0.00%) | 876 (21.80%) | 1567 (38.99%) | 788 (19.61%) | 788 (19.61%) | 800 (19.91%) | 1746 (43.44%) | 931 (23.16%) | 542 (13.49%) |
| Family member was seriously injured? [Parent] | 002 | 630 (9.90%) | 5736 (90.10%) | 0 (0.00%) | 0 (0.00%) | 78 (12.38%) | 322 (51.11%) | 119 (18.89%) | 111 (17.62%) | 86 (13.65%) | 237 (37.62%) | 180 (28.57%) | 127 (20.16%) |
| Saw crime or accident? [Parent] | 003 | 367 (5.77%) | 5999 (94.23%) | 0 (0.00%) | 0 (0.00%) | 28 (7.63%) | 226 (61.58%) | 74 (20.16%) | 39 (10.63%) | 73 (19.89%) | 167 (45.50%) | 70 (19.07%) | 57 (15.53%) |
| Close friend died? [Parent] | 004 | 138 (2.17%) | 6228 (97.83%) | 0 (0.00%) | 0 (0.00%) | 15 (10.87%) | 82 (59.42%) | 25 (18.12%) | 16 (11.59%) | 10 (7.25%) | 48 (34.78%) | 44 (31.88%) | 36 (26.09%) |
| Lost a close friend? [Parent] | 005 | 327 (5.14%) | 6039 (94.86%) | 0 (0.00%) | 0 (0.00%) | 18 (5.50%) | 234 (71.56%) | 46 (14.07%) | 29 (8.87%) | 7 (2.14%) | 94 (28.75%) | 120 (36.70%) | 106 (32.42%) |
| Close friend was seriously sick/injured? [Parent] | 006 | 188 (2.95%) | 6178 (97.05%) | 0 (0.00%) | 0 (0.00%) | 48 (25.53%) | 95 (50.53%) | 29 (15.43%) | 16 (8.51%) | 15 (7.98%) | 80 (42.55%) | 56 (29.79%) | 37 (19.68%) |
| Negative change in parent's financial situation? [Parent] | 007 | 1025 (16.10%) | 5341 (83.90%) | 0 (0.00%) | 0 (0.00%) | 105 (10.24%) | 406 (39.61%) | 298 (29.07%) | 216 (21.07%) | 153 (14.93%) | 463 (45.17%) | 295 (28.78%) | 114 (11.12%) |
| Family member had drug and/or alcohol problem? [Parent] | 008 | 763 (11.99%) | 5603 (88.01%) | 0 (0.00%) | 0 (0.00%) | 50 (6.55%) | 391 (51.25%) | 177 (23.20%) | 145 (19.00%) | 154 (20.18%) | 281 (36.83%) | 188 (24.64%) | 140 (18.35%) |
| Child was seriously sick? [Parent] | 009 | 151 (2.37%) | 6215 (97.63%) | 0 (0.00%) | 0 (0.00%) | 19 (12.58%) | 100 (66.23%) | 17 (11.26%) | 15 (9.93%) | 16 (10.60%) | 38 (25.17%) | 58 (38.41%) | 39 (25.83%) |
| Child was seriously injured? [Parent] | 010 | 118 (1.85%) | 6240 (98.02%) | 0 (0.00%) | 0 (0.00%) | 21 (17.80%) | 75 (63.56%) | 14 (11.86%) | 8 (6.78%) | 5 (4.24%) | 51 (43.22%) | 34 (28.81%) | 28 (23.73%) |
| Parents argued more than previously? [Parent] | 011 | 747 (11.73%) | 5619 (88.27%) | 0 (0.00%) | 0 (0.00%) | 22 (2.95%) | 543 (72.69%) | 152 (20.35%) | 30 (4.02%) | 33 (4.42%) | 281 (37.62%) | 267 (35.74%) | 166 (22.22%) |
| Parent figure lost job? [Parent] | 012 | 942 (14.80%) | 5424 (85.20%) | 0 (0.00%) | 0 (0.00%) | 181 (19.21%) | 252 (26.75%) | 267 (28.34%) | 242 (25.69%) | 280 (29.72%) | 395 (41.93%) | 194 (20.59%) | 73 (7.75%) |
| One parent was away from home more often? [Parent] | 013 | 1182 (18.57%) | 5184 (81.43%) | 0 (0.00%) | 0 (0.00%) | 245 (20.73%) | 472 (39.93%) | 340 (28.76%) | 125 (10.58%) | 109 (9.22%) | 493 (41.71%) | 355 (30.03%) | 225 (19.04%) |
| Someone in the family was arrested? [Parent] | 014 | 389 (6.11%) | 5977 (93.89%) | 0 (0.00%) | 0 (0.00%) | 17 (4.37%) | 221 (56.81%) | 88 (22.62%) | 63 (16.20%) | 70 (17.99%) | 131 (33.68%) | 91 (23.39%) | 97 (24.94%) |
| Family member had mental/emotional problem? [Parent] | 015 | 988 (15.52%) | 5378 (84.48%) | 0 (0.00%) | 0 (0.00%) | 70 (7.09%) | 471 (47.67%) | 320 (32.39%) | 127 (12.85%) | 112 (11.34%) | 408 (41.30%) | 311 (31.48%) | 157 (15.89%) |
| Sibling left home? [Parent] | 016 | 604 (9.49%) | 5762 (90.51%) | 0 (0.00%) | 0 (0.00%) | 237 (39.24%) | 161 (26.66%) | 127 (21.03%) | 79 (13.08%) | 76 (12.58%) | 273 (45.20%) | 175 (28.97%) | 80 (13.25%) |
| Was a victim of crime/violence/assault? [Parent] | 017 | 79 (1.24%) | 6287 (98.76%) | 0 (0.00%) | 0 (0.00%) | 3 (3.80%) | 70 (88.61%) | 3 (3.80%) | 3 (3.80%) | 5 (6.33%) | 17 (21.52%) | 19 (24.05%) | 38 (48.10%) |
| Parents separated or divorced? [Parent] | 018 | 949 (14.91%) | 5417 (85.09%) | 0 (0.00%) | 0 (0.00%) | 186 (19.60%) | 435 (45.84%) | 220 (23.18%) | 108 (11.38%) | 67 (7.06%) | 224 (23.60%) | 310 (32.67%) | 348 (36.67%) |
| Parents/caregiver got into trouble with the law? [Parent] | 019 | 205 (3.22%) | 6161 (96.78%) | 0 (0.00%) | 0 (0.00%) | 10 (4.88%) | 125 (60.98%) | 40 (19.51%) | 30 (14.63%) | 24 (11.71%) | 67 (32.68%) | 57 (27.80%) | 57 (27.80%) |
| Attended a new school? [Parent] | 020 | 2623 (41.20%) | 3743 (58.80%) | 0 (0.00%) | 0 (0.00%) | 2155 (82.16%) | 189 (7.21%) | 213 (8.12%) | 66 (2.52%) | 536 (20.43%) | 849 (32.37%) | 680 (25.92%) | 558 (21.27%) |
| Family moved? [Parent] | 021 | 1685 (26.47%) | 4681 (73.53%) | 0 (0.00%) | 0 (0.00%) | 1315 (78.04%) | 144 (8.55%) | 182 (10.80%) | 44 (2.61%) | 375 (22.26%) | 543 (32.23%) | 398 (23.62%) | 369 (21.90%) |
| One of the parents/caregivers went to jail? [Parent] | 022 | 236 (3.71%) | 6130 (96.29%) | 0 (0.00%) | 0 (0.00%) | 12 (5.08%) | 143 (60.59%) | 54 (22.88%) | 27 (11.44%) | 30 (12.71%) | 70 (29.66%) | 70 (29.66%) | 66 (27.97%) |
| Got new step-parent? [Parent] | 023 | 457 (7.18%) | 5909 (92.82%) | 0 (0.00%) | 0 (0.00%) | 289 (63.24%) | 49 (10.72%) | 88 (19.26%) | 31 (6.78%) | 68 (14.88%) | 122 (26.70%) | 108 (23.63%) | 159 (34.79%) |
| Parent/caregiver got a new job? [Parent] | 024 | 1719 (27.00%) | 4647 (73.00%) | 0 (0.00%) | 0 (0.00%) | 1286 (74.81%) | 45 (2.62%) | 239 (13.90%) | 149 (8.67%) | 656 (38.16%) | 615 (35.78%) | 313 (18.21%) | 135 (7.85%) |
| Got new sibling? [Parent] | 025 | 1074 (16.87%) | 5292 (83.13%) | 0 (0.00%) | 0 (0.00%) | 936 (87.15%) | 38 (3.54%) | 81 (7.54%) | 19 (1.77%) | 163 (15.18%) | 232 (21.60%) | 265 (24.67%) | 414 (38.55%) |

**Table S4.5. Caregiver report – 4 year follow-up**

| **Item** | **Code** | **4 year follow-up** | | | | | | | | | | | |
| --- | --- | --- | --- | --- | --- | --- | --- | --- | --- | --- | --- | --- | --- |
|  |  | **Count (#, %)** | | | | **Experience (#, % of people who have count==1)** | | | | **Severity (#, % of people who have count==1)** | | | |
|  |  | **1** | **0** | **999** | **777** | **1** | **2** | **999** | **444** | **0** | **1** | **2** | **3** |
|  |  | **Yes** | **No** | **Dont know** | **Decline to answer** | **Mostly good** | **Mostly bad** | **Dont know** | **Not applicable** | **Not at all** | **A little** | **Some** | **A lot** |
| **SLE questions** | | | | | | | | | | | | | |
| Someone in family died? [Parent] | 001 | 3353 (66.59%) | 1675 (33.27%) | 0 (0.00%) | 0 (0.00%) | 681 (20.31%) | 1373 (40.95%) | 683 (20.37%) | 616 (18.37%) | 620 (18.49%) | 1408 (41.99%) | 863 (25.74%) | 462 (13.78%) |
| Family member was seriously injured? [Parent] | 002 | 486 (9.65%) | 4542 (90.21%) | 0 (0.00%) | 0 (0.00%) | 61 (12.55%) | 273 (56.17%) | 82 (16.87%) | 70 (14.40%) | 59 (12.14%) | 184 (37.86%) | 140 (28.81%) | 103 (21.19%) |
| Saw crime or accident? [Parent] | 003 | 304 (6.04%) | 4724 (93.82%) | 0 (0.00%) | 0 (0.00%) | 25 (8.22%) | 181 (59.54%) | 61 (20.07%) | 37 (12.17%) | 59 (19.41%) | 131 (43.09%) | 60 (19.74%) | 54 (17.76%) |
| Close friend died? [Parent] | 004 | 146 (2.90%) | 4882 (96.96%) | 0 (0.00%) | 0 (0.00%) | 10 (6.85%) | 101 (69.18%) | 22 (15.07%) | 13 (8.90%) | 11 (7.53%) | 37 (25.34%) | 52 (35.62%) | 46 (31.51%) |
| Lost a close friend? [Parent] | 005 | 337 (6.69%) | 4691 (93.17%) | 0 (0.00%) | 0 (0.00%) | 30 (8.90%) | 254 (75.37%) | 41 (12.17%) | 12 (3.56%) | 8 (2.37%) | 81 (24.04%) | 134 (39.76%) | 114 (33.83%) |
| Close friend was seriously sick/injured? [Parent] | 006 | 118 (2.34%) | 4910 (97.52%) | 0 (0.00%) | 0 (0.00%) | 20 (16.95%) | 62 (52.54%) | 22 (18.64%) | 14 (11.86%) | 5 (4.24%) | 56 (47.46%) | 34 (28.81%) | 23 (19.49%) |
| Negative change in parent's financial situation? [Parent] | 007 | 798 (15.85%) | 4230 (84.01%) | 0 (0.00%) | 0 (0.00%) | 99 (12.41%) | 320 (40.10%) | 213 (26.69%) | 166 (20.80%) | 112 (14.04%) | 384 (48.12%) | 212 (26.57%) | 90 (11.28%) |
| Family member had drug and/or alcohol problem? [Parent] | 008 | 600 (11.92%) | 4428 (87.94%) | 0 (0.00%) | 0 (0.00%) | 42 (7.00%) | 330 (55.00%) | 137 (22.83%) | 91 (15.17%) | 91 (15.17%) | 201 (33.50%) | 174 (29.00%) | 134 (22.33%) |
| Child was seriously sick? [Parent] | 009 | 157 (3.12%) | 4871 (96.74%) | 0 (0.00%) | 0 (0.00%) | 26 (16.56%) | 92 (58.60%) | 23 (14.65%) | 16 (10.19%) | 13 (8.28%) | 45 (28.66%) | 51 (32.48%) | 48 (30.57%) |
| Child was seriously injured? [Parent] | 010 | 121 (2.40%) | 4902 (97.36%) | 0 (0.00%) | 0 (0.00%) | 18 (14.88%) | 85 (70.25%) | 11 (9.09%) | 7 (5.79%) | 5 (4.13%) | 34 (28.10%) | 45 (37.19%) | 37 (30.58%) |
| Parents argued more than previously? [Parent] | 011 | 598 (11.88%) | 4430 (87.98%) | 0 (0.00%) | 0 (0.00%) | 20 (3.34%) | 437 (73.08%) | 117 (19.57%) | 24 (4.01%) | 16 (2.68%) | 198 (33.11%) | 252 (42.14%) | 132 (22.07%) |
| Parent figure lost job? [Parent] | 012 | 789 (15.67%) | 4239 (84.19%) | 0 (0.00%) | 0 (0.00%) | 163 (20.66%) | 199 (25.22%) | 243 (30.80%) | 184 (23.32%) | 199 (25.22%) | 365 (46.26%) | 161 (20.41%) | 64 (8.11%) |
| One parent was away from home more often? [Parent] | 013 | 829 (16.46%) | 4199 (83.40%) | 0 (0.00%) | 0 (0.00%) | 177 (21.35%) | 292 (35.22%) | 256 (30.88%) | 104 (12.55%) | 85 (10.25%) | 318 (38.36%) | 283 (34.14%) | 143 (17.25%) |
| Someone in the family was arrested? [Parent] | 014 | 299 (5.94%) | 4729 (93.92%) | 0 (0.00%) | 0 (0.00%) | 18 (6.02%) | 156 (52.17%) | 60 (20.07%) | 65 (21.74%) | 60 (20.07%) | 101 (33.78%) | 66 (22.07%) | 72 (24.08%) |
| Family member had mental/emotional problem? [Parent] | 015 | 852 (16.92%) | 4176 (82.94%) | 0 (0.00%) | 0 (0.00%) | 63 (7.39%) | 428 (50.23%) | 254 (29.81%) | 107 (12.56%) | 77 (9.04%) | 327 (38.38%) | 333 (39.08%) | 115 (13.50%) |
| Sibling left home? [Parent] | 016 | 685 (13.60%) | 4343 (86.26%) | 0 (0.00%) | 0 (0.00%) | 339 (49.49%) | 104 (15.18%) | 155 (22.63%) | 87 (12.70%) | 98 (14.31%) | 302 (44.09%) | 217 (31.68%) | 68 (9.93%) |
| Was a victim of crime/violence/assault? [Parent] | 017 | 103 (2.05%) | 4925 (97.82%) | 0 (0.00%) | 0 (0.00%) | 1 (0.97%) | 92 (89.32%) | 5 (4.85%) | 5 (4.85%) | 3 (2.91%) | 19 (18.45%) | 41 (39.81%) | 40 (38.83%) |
| Parents separated or divorced? [Parent] | 018 | 855 (16.98%) | 4173 (82.88%) | 0 (0.00%) | 0 (0.00%) | 233 (27.25%) | 346 (40.47%) | 190 (22.22%) | 86 (10.06%) | 66 (7.72%) | 207 (24.21%) | 298 (34.85%) | 284 (33.22%) |
| Parents/caregiver got into trouble with the law? [Parent] | 019 | 157 (3.12%) | 4871 (96.74%) | 0 (0.00%) | 0 (0.00%) | 10 (6.37%) | 93 (59.24%) | 31 (19.75%) | 23 (14.65%) | 25 (15.92%) | 43 (27.39%) | 46 (29.30%) | 43 (27.39%) |
| Attended a new school? [Parent] | 020 | 2095 (41.61%) | 2933 (58.25%) | 0 (0.00%) | 0 (0.00%) | 1670 (79.71%) | 153 (7.30%) | 208 (9.93%) | 64 (3.05%) | 354 (16.90%) | 593 (28.31%) | 598 (28.54%) | 550 (26.25%) |
| Family moved? [Parent] | 021 | 1353 (26.87%) | 3675 (72.99%) | 0 (0.00%) | 0 (0.00%) | 1068 (78.94%) | 111 (8.20%) | 123 (9.09%) | 51 (3.77%) | 249 (18.40%) | 421 (31.12%) | 356 (26.31%) | 327 (24.17%) |
| One of the parents/caregivers went to jail? [Parent] | 022 | 163 (3.24%) | 4865 (96.62%) | 0 (0.00%) | 0 (0.00%) | 12 (7.36%) | 88 (53.99%) | 37 (22.70%) | 26 (15.95%) | 25 (15.34%) | 49 (30.06%) | 48 (29.45%) | 41 (25.15%) |
| Got new step-parent? [Parent] | 023 | 390 (7.75%) | 4638 (92.12%) | 0 (0.00%) | 0 (0.00%) | 276 (70.77%) | 43 (11.03%) | 59 (15.13%) | 12 (3.08%) | 53 (13.59%) | 91 (23.33%) | 113 (28.97%) | 133 (34.10%) |
| Parent/caregiver got a new job? [Parent] | 024 | 1411 (28.02%) | 3617 (71.84%) | 0 (0.00%) | 0 (0.00%) | 1052 (74.56%) | 31 (2.20%) | 174 (12.33%) | 154 (10.91%) | 548 (38.84%) | 535 (37.92%) | 247 (17.51%) | 81 (5.74%) |
| Got new sibling? [Parent] | 025 | 823 (16.35%) | 4205 (83.52%) | 0 (0.00%) | 0 (0.00%) | 708 (86.03%) | 34 (4.13%) | 67 (8.14%) | 14 (1.70%) | 104 (12.64%) | 155 (18.83%) | 194 (23.57%) | 370 (44.96%) |
| One of the parents/caregivers was deported? [Parent] | 026 | 3 (0.06%) | 1877 (37.28%) | 0 (0.00%) | 0 (0.00%) | 1 (33.33%) | 0 (0.00%) | 2 (66.67%) | 0 (0.00%) | 1 (33.33%) | 0 (0.00%) | 2 (66.67%) | 0 (0.00%) |
| One of the parents/caregivers was hospitalized? [Parent] | 027 | 177 (3.52%) | 1703 (33.82%) | 0 (0.00%) | 0 (0.00%) | 29 (16.38%) | 75 (42.37%) | 41 (23.16%) | 32 (18.08%) | 27 (15.25%) | 79 (44.63%) | 44 (24.86%) | 27 (15.25%) |
| Child placed in foster care? [Parent] | 028 | 16 (0.32%) | 1864 (37.02%) | 0 (0.00%) | 0 (0.00%) | 5 (31.25%) | 9 (56.25%) | 2 (12.50%) | 0 (0.00%) | 0 (0.00%) | 0 (0.00%) | 2 (12.50%) | 14 (87.50%) |
| Saw or heard someone getting hit, slapped, punched, or beaten up in their school or neighborhood? [Parent] | 029 | 216 (4.29%) | 1664 (33.05%) | 0 (0.00%) | 0 (0.00%) | 9 (4.17%) | 69 (31.94%) | 88 (40.74%) | 49 (22.69%) | 89 (41.20%) | 84 (38.89%) | 29 (13.43%) | 13 (6.02%) |
| Saw or heard someone being shot at (but not actually wounded) in their school or in their neighborhood? [Parent] | 030 | 44 (0.87%) | 1836 (36.46%) | 0 (0.00%) | 0 (0.00%) | 0 (0.00%) | 24 (54.55%) | 10 (22.73%) | 10 (22.73%) | 7 (15.91%) | 19 (43.18%) | 10 (22.73%) | 8 (18.18%) |
| Had a lockdown at their school due to concerns about a school shooting or violence? [Parent] | 031 | 413 (8.20%) | 1467 (29.14%) | 0 (0.00%) | 0 (0.00%) | 52 (12.59%) | 128 (30.99%) | 166 (40.19%) | 67 (16.22%) | 151 (36.56%) | 192 (46.49%) | 54 (13.08%) | 16 (3.87%) |
| Your family was without a home or place to stay/live? [Parent] | 032 | 68 (1.35%) | 4960 (98.51%) | 0 (0.00%) | 0 (0.00%) | 0 (0.00%) | 0 (0.00%) | 0 (0.00%) | 0 (0.00%) | 0 (0.00%) | 0 (0.00%) | 0 (0.00%) | 0 (0.00%) |

**Table S4.6. Caregiver report – 6 year follow-up**

| **Item** | **Code** | **6 year follow-up** | | | | | | | | | | | |
| --- | --- | --- | --- | --- | --- | --- | --- | --- | --- | --- | --- | --- | --- |
|  |  | **Count (#, %)** | | | | **Experience (#, % of people who have count==1)** | | | | **Severity (#, % of people who have count==1)** | | | |
|  |  | **1** | **0** | **999** | **777** | **1** | **2** | **999** | **444** | **0** | **1** | **2** | **3** |
|  |  | **Yes** | **No** | **Dont know** | **Decline to answer** | **Mostly good** | **Mostly bad** | **Dont know** | **Not applicable** | **Not at all** | **A little** | **Some** | **A lot** |
| **SLE questions** | | | | | | | | | | | | | |
| Someone in family died? [Parent] | 001 | 4060 (68.20%) | 1884 (31.65%) | 0 (0.00%) | 6 (0.10%) | 853 (21.01%) | 1636 (40.30%) | 825 (20.32%) | 744 (18.33%) | 699 (17.22%) | 1763 (43.42%) | 1021 (25.15%) | 558 (13.74%) |
| Family member was seriously injured? [Parent] | 002 | 582 (9.78%) | 5363 (90.09%) | 2 (0.03%) | 3 (0.05%) | 90 (15.46%) | 301 (51.72%) | 102 (17.53%) | 89 (15.29%) | 64 (11.00%) | 236 (40.55%) | 173 (29.73%) | 106 (18.21%) |
| Saw crime or accident? [Parent] | 003 | 389 (6.53%) | 5546 (93.16%) | 13 (0.22%) | 2 (0.03%) | 30 (7.71%) | 238 (61.18%) | 66 (16.97%) | 55 (14.14%) | 66 (16.97%) | 160 (41.13%) | 93 (23.91%) | 67 (17.22%) |
| Close friend died? [Parent] | 004 | 269 (4.52%) | 5677 (95.36%) | 2 (0.03%) | 2 (0.03%) | 23 (8.55%) | 198 (73.61%) | 29 (10.78%) | 19 (7.06%) | 6 (2.23%) | 68 (25.28%) | 96 (35.69%) | 99 (36.80%) |
| Lost a close friend? [Parent] | 005 | 487 (8.18%) | 5452 (91.58%) | 9 (0.15%) | 2 (0.03%) | 42 (8.62%) | 358 (73.51%) | 68 (13.96%) | 19 (3.90%) | 10 (2.05%) | 120 (24.64%) | 192 (39.43%) | 164 (33.68%) |
| Close friend was seriously sick/injured? [Parent] | 006 | 159 (2.67%) | 5784 (97.16%) | 6 (0.10%) | 1 (0.02%) | 27 (16.98%) | 80 (50.31%) | 36 (22.64%) | 16 (10.06%) | 14 (8.81%) | 60 (37.74%) | 49 (30.82%) | 34 (21.38%) |
| Negative change in parent's financial situation? [Parent] | 007 | 985 (16.55%) | 4956 (83.25%) | 7 (0.12%) | 2 (0.03%) | 91 (9.24%) | 447 (45.38%) | 256 (25.99%) | 189 (19.19%) | 123 (12.49%) | 406 (41.22%) | 304 (30.86%) | 140 (14.21%) |
| Family member had drug and/or alcohol problem? [Parent] | 008 | 854 (14.35%) | 5088 (85.47%) | 6 (0.10%) | 2 (0.03%) | 41 (4.80%) | 511 (59.84%) | 192 (22.48%) | 110 (12.88%) | 88 (10.30%) | 292 (34.19%) | 271 (31.73%) | 203 (23.77%) |
| Child was seriously sick? [Parent] | 009 | 183 (3.07%) | 5765 (96.84%) | 1 (0.02%) | 1 (0.02%) | 21 (11.48%) | 128 (69.95%) | 24 (13.11%) | 10 (5.46%) | 12 (6.56%) | 42 (22.95%) | 47 (25.68%) | 82 (44.81%) |
| Child was seriously injured? [Parent] | 010 | 144 (2.42%) | 5798 (97.40%) | 1 (0.02%) | 2 (0.03%) | 16 (11.11%) | 99 (68.75%) | 18 (12.50%) | 11 (7.64%) | 2 (1.39%) | 34 (23.61%) | 51 (35.42%) | 57 (39.58%) |
| Parents argued more than previously? [Parent] | 011 | 713 (11.98%) | 5226 (87.79%) | 7 (0.12%) | 4 (0.07%) | 18 (2.52%) | 531 (74.47%) | 129 (18.09%) | 35 (4.91%) | 18 (2.52%) | 222 (31.14%) | 278 (38.99%) | 193 (27.07%) |
| Parent figure lost job? [Parent] | 012 | 974 (16.36%) | 4972 (83.52%) | 3 (0.05%) | 1 (0.02%) | 176 (18.07%) | 288 (29.57%) | 280 (28.75%) | 229 (23.51%) | 279 (28.64%) | 378 (38.81%) | 231 (23.72%) | 77 (7.91%) |
| One parent was away from home more often? [Parent] | 013 | 940 (15.79%) | 5000 (83.99%) | 6 (0.10%) | 4 (0.07%) | 221 (23.51%) | 301 (32.02%) | 304 (32.34%) | 113 (12.02%) | 105 (11.17%) | 355 (37.77%) | 298 (31.70%) | 171 (18.19%) |
| Someone in the family was arrested? [Parent] | 014 | 428 (7.19%) | 5518 (92.69%) | 2 (0.03%) | 2 (0.03%) | 25 (5.84%) | 225 (52.57%) | 97 (22.66%) | 80 (18.69%) | 91 (21.26%) | 123 (28.74%) | 98 (22.90%) | 108 (25.23%) |
| Family member had mental/emotional problem? [Parent] | 015 | 1266 (21.27%) | 4674 (78.52%) | 8 (0.13%) | 2 (0.03%) | 86 (6.79%) | 608 (48.03%) | 403 (31.83%) | 168 (13.27%) | 148 (11.69%) | 468 (36.97%) | 418 (33.02%) | 220 (17.38%) |
| Sibling left home? [Parent] | 016 | 1226 (20.59%) | 4722 (79.32%) | 1 (0.02%) | 1 (0.02%) | 650 (53.02%) | 161 (13.13%) | 283 (23.08%) | 131 (10.69%) | 196 (15.99%) | 563 (45.92%) | 328 (26.75%) | 130 (10.60%) |
| Was a victim of crime/violence/assault? [Parent] | 017 | 167 (2.81%) | 5778 (97.06%) | 3 (0.05%) | 2 (0.03%) | 3 (1.80%) | 144 (86.23%) | 12 (7.19%) | 8 (4.79%) | 7 (4.19%) | 30 (17.96%) | 51 (30.54%) | 76 (45.51%) |
| Parents separated or divorced? [Parent] | 018 | 1154 (19.39%) | 4790 (80.46%) | 0 (0.00%) | 6 (0.10%) | 317 (27.47%) | 430 (37.26%) | 278 (24.09%) | 127 (11.01%) | 81 (7.02%) | 265 (22.96%) | 374 (32.41%) | 424 (36.74%) |
| Parents/caregiver got into trouble with the law? [Parent] | 019 | 238 (4.00%) | 5708 (95.88%) | 1 (0.02%) | 3 (0.05%) | 16 (6.72%) | 117 (49.16%) | 49 (20.59%) | 55 (23.11%) | 50 (21.01%) | 65 (27.31%) | 60 (25.21%) | 60 (25.21%) |
| Attended a new school? [Parent] | 020 | 2359 (39.63%) | 3589 (60.29%) | 0 (0.00%) | 2 (0.03%) | 1904 (80.71%) | 175 (7.42%) | 206 (8.73%) | 73 (3.09%) | 403 (17.08%) | 666 (28.23%) | 651 (27.60%) | 627 (26.58%) |
| Family moved? [Parent] | 021 | 1642 (27.58%) | 4307 (72.35%) | 0 (0.00%) | 1 (0.02%) | 1257 (76.55%) | 132 (8.04%) | 189 (11.51%) | 64 (3.90%) | 329 (20.04%) | 464 (28.26%) | 426 (25.94%) | 415 (25.27%) |
| One of the parents/caregivers went to jail? [Parent] | 022 | 235 (3.95%) | 5713 (95.97%) | 0 (0.00%) | 2 (0.03%) | 14 (5.96%) | 128 (54.47%) | 55 (23.40%) | 38 (16.17%) | 38 (16.17%) | 72 (30.64%) | 49 (20.85%) | 70 (29.79%) |
| Got new step-parent? [Parent] | 023 | 542 (9.10%) | 5404 (90.78%) | 1 (0.02%) | 3 (0.05%) | 369 (68.08%) | 49 (9.04%) | 100 (18.45%) | 24 (4.43%) | 59 (10.89%) | 137 (25.28%) | 151 (27.86%) | 192 (35.42%) |
| Parent/caregiver got a new job? [Parent] | 024 | 1727 (29.01%) | 4221 (70.91%) | 1 (0.02%) | 1 (0.02%) | 1283 (74.29%) | 36 (2.08%) | 243 (14.07%) | 165 (9.55%) | 682 (39.49%) | 608 (35.21%) | 301 (17.43%) | 126 (7.30%) |
| Got new sibling? [Parent] | 025 | 1049 (17.62%) | 4900 (82.31%) | 0 (0.00%) | 1 (0.02%) | 880 (83.89%) | 36 (3.43%) | 114 (10.87%) | 18 (1.72%) | 121 (11.53%) | 228 (21.73%) | 252 (24.02%) | 440 (41.94%) |
| One of the parents/caregivers was deported? [Parent] | 026 | 17 (0.29%) | 5930 (99.61%) | 0 (0.00%) | 2 (0.03%) | 1 (5.88%) | 9 (52.94%) | 6 (35.29%) | 1 (5.88%) | 4 (23.53%) | 1 (5.88%) | 5 (29.41%) | 7 (41.18%) |
| One of the parents/caregivers was hospitalized? [Parent] | 027 | 708 (11.89%) | 5240 (88.02%) | 0 (0.00%) | 1 (0.02%) | 138 (19.49%) | 273 (38.56%) | 207 (29.24%) | 90 (12.71%) | 132 (18.64%) | 270 (38.14%) | 196 (27.68%) | 107 (15.11%) |
| Child placed in foster care? [Parent] | 028 | 41 (0.69%) | 5907 (99.23%) | 0 (0.00%) | 1 (0.02%) | 16 (39.02%) | 16 (39.02%) | 8 (19.51%) | 1 (2.44%) | 2 (4.88%) | 3 (7.32%) | 5 (12.20%) | 30 (73.17%) |
| Saw or heard someone getting hit, slapped, punched, or beaten up in their school or neighborhood? [Parent] | 029 | 598 (10.05%) | 5318 (89.33%) | 31 (0.52%) | 2 (0.03%) | 17 (2.84%) | 190 (31.77%) | 266 (44.48%) | 125 (20.90%) | 242 (40.47%) | 230 (38.46%) | 78 (13.04%) | 34 (5.69%) |
| Saw or heard someone being shot at (but not actually wounded) in their school or in their neighborhood? [Parent] | 030 | 107 (1.80%) | 5830 (97.93%) | 11 (0.18%) | 1 (0.02%) | 0 (0.00%) | 68 (63.55%) | 30 (28.04%) | 9 (8.41%) | 15 (14.02%) | 43 (40.19%) | 27 (25.23%) | 18 (16.82%) |
| Had a lockdown at their school due to concerns about a school shooting or violence? [Parent] | 031 | 1650 (27.72%) | 4277 (71.85%) | 20 (0.34%) | 2 (0.03%) | 212 (12.85%) | 565 (34.24%) | 602 (36.48%) | 271 (16.42%) | 606 (36.73%) | 739 (44.79%) | 235 (14.24%) | 48 (2.91%) |
| Your family was without a home or place to stay/live? [Parent] | 032 | 111 (1.86%) | 5837 (98.05%) | 0 (0.00%) | 1 (0.02%) | 11 (9.91%) | 73 (65.77%) | 19 (17.12%) | 7 (6.31%) | 3 (2.70%) | 25 (22.52%) | 30 (27.03%) | 51 (45.95%) |
| Your child was thrown out of their house? [Parent] | 033 | 28 (0.47%) | 5920 (99.45%) | 0 (0.00%) | 1 (0.02%) | 2 (7.14%) | 22 (78.57%) | 2 (7.14%) | 2 (7.14%) | 1 (3.57%) | 3 (10.71%) | 7 (25.00%) | 16 (57.14%) |

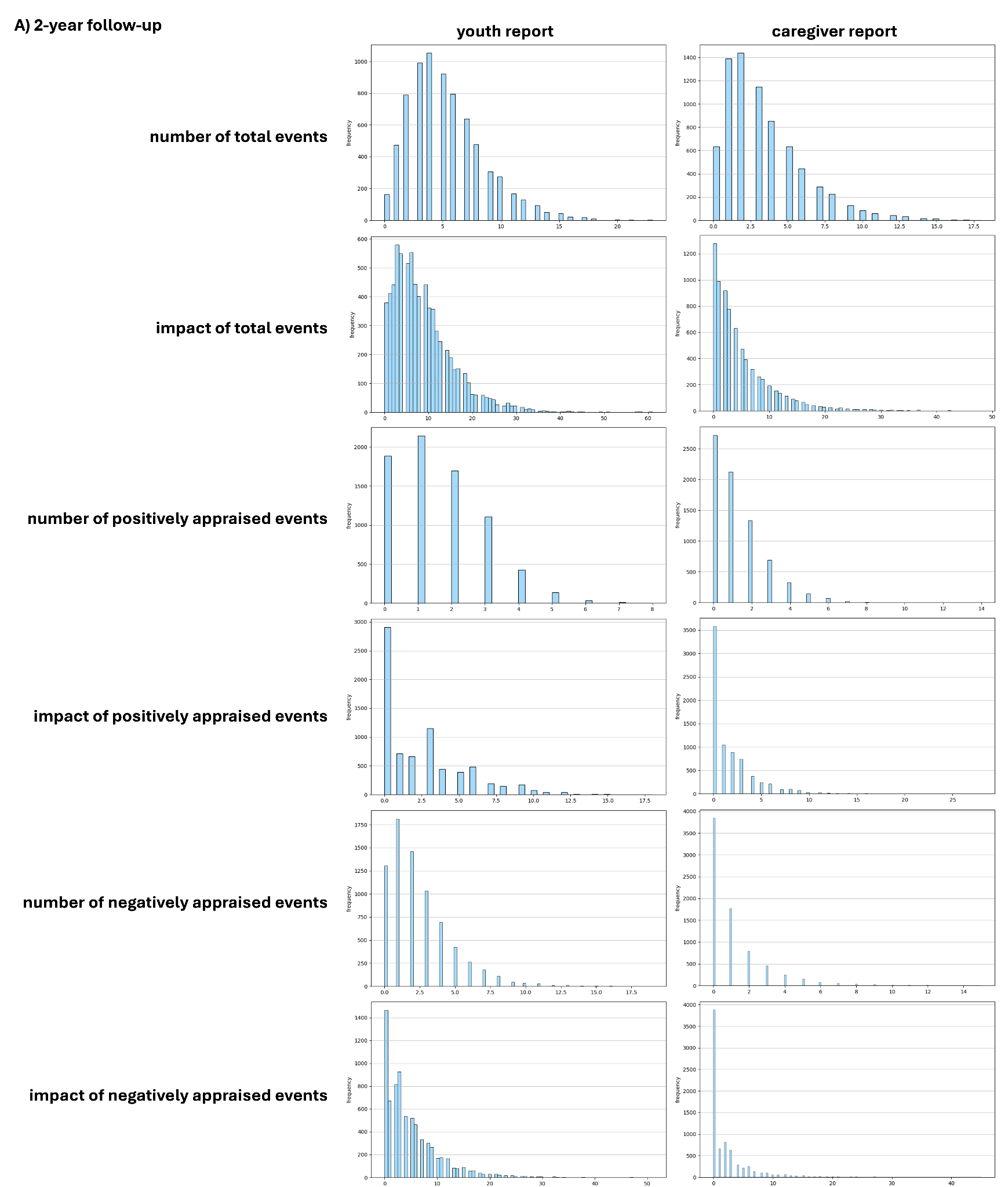

**Supplementary Figure 5A. Histograms depicting the distribution of values in the target variables at the 2-year follow-up.**

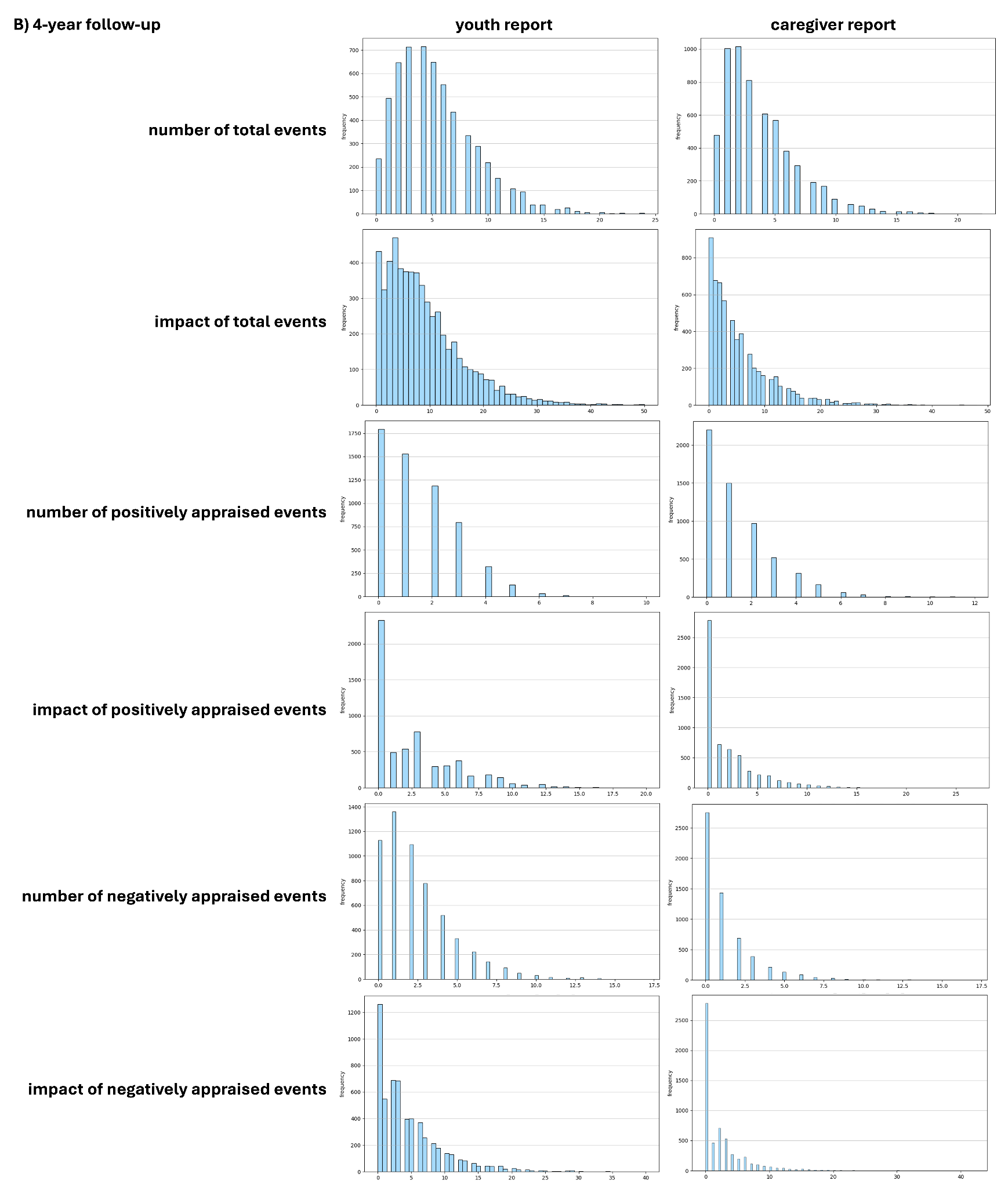

**Supplementary Figure 5B. Histograms depicting the distribution of values in the target variables at the 4-year follow-up.**

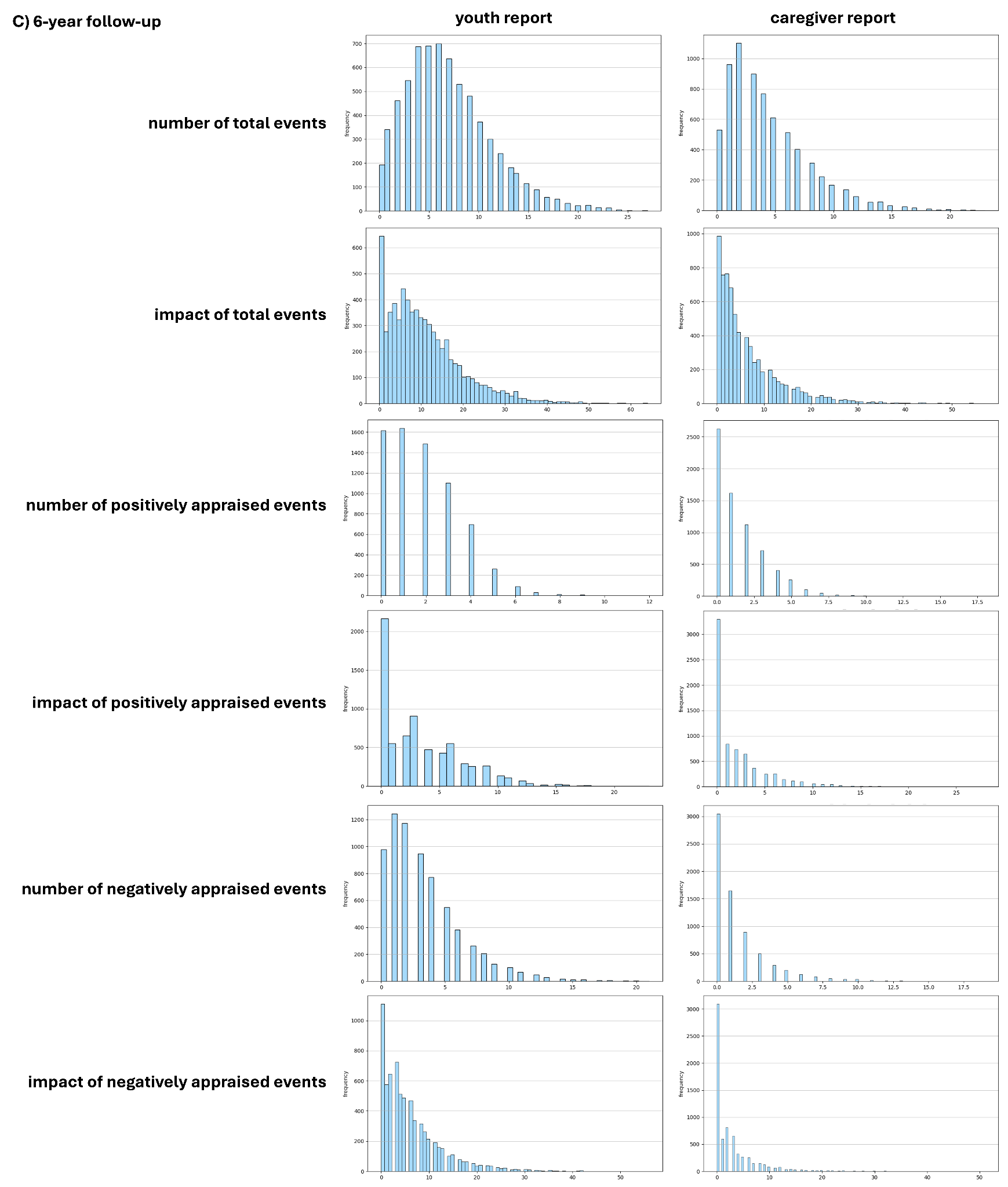

**Supplementary Figure 5C. Histograms depicting the distribution of values in the target variables at the 6-year follow-up.**

**Table S5. Counts of zeros per target variable**

Count of zeros for each individual target variable at the (A) 2-year follow-up, (B) 4-year follow-up, and (C) 6-year follow-up time point for the youth report and caregiver report.

|  |  | **youth report** | **caregiver report** |
| --- | --- | --- | --- |
| **A) 2-year follow-up** | **number of total events** | 162 | 634 |
|  | **impact of total events** | 379 | 1278 |
|  | **number of positively appraised events** | 1884 | 2717 |
|  | **impact of positively appraised events** | 2907 | 3581 |
|  | **number of negatively appraised events** | 1306 | 3841 |
|  | **impact of negatively appraised events** | 1465 | 3880 |
| **B) 4-year follow-up** | **number of total events** | 236 | 478 |
|  | **impact of total events** | 432 | 909 |
|  | **number of positively appraised events** | 1793 | 2201 |
|  | **impact of positively appraised events** | 2326 | 2787 |
|  | **number of negatively appraised events** | 1128 | 2752 |
|  | **impact of negatively appraised events** | 1261 | 2782 |
| **C) 6-year follow-up** | **number of total events** | 193 | 531 |
|  | **impact of total events** | 366 | 986 |
|  | **number of positively appraised events** | 1615 | 2628 |
|  | **impact of positively appraised events** | 2165 | 3296 |
|  | **number of negatively appraised events** | 977 | 3045 |
|  | **impact of negatively appraised events** | 1110 | 3093 |

**Table S6. Individual sample sizes per target variable**

Sample sizes after excluding zeros for each individual target variable, filtering for siblings and small sites for the (A) 2-year follow-up, (B) 4-year follow-up, and (C) 6-year follow-up time point for the youth report and caregiver report.

|  |  | **youth report** | **caregiver report** |
| --- | --- | --- | --- |
| **A) 2-year follow-up** | **number of total events** | 6250 | 5888 |
|  | **impact of total events** | 6092 | 5371 |
|  | **number of positively appraised events** | 4922 | 4179 |
|  | **impact of positively appraised events** | 4118 | 3441 |
|  | **number of negatively appraised events** | 5408 | 3199 |
|  | **impact of negatively appraised events** | 5284 | 3161 |
| **B) 4-year follow-up** | **number of total events** | 4851 | 4659 |
|  | **impact of total events** | 4694 | 4292 |
|  | **number of positively appraised events** | 3605 | 3190 |
|  | **impact of positively appraised events** | 3156 | 2677 |
|  | **number of negatively appraised events** | 4145 | 2715 |
|  | **impact of negatively appraised events** | 4035 | 2691 |
| **C) 6-year follow-up** | **number of total events** | 5810 | 5550 |
|  | **impact of total events** | 5690 | 5190 |
|  | **number of positively appraised events** | 4715 | 3790 |
|  | **impact of positively appraised events** | 4272 | 3234 |
|  | **number of negatively appraised events** | 5234 | 3457 |
|  | **impact of negatively appraised events** | 5134 | 3419 |

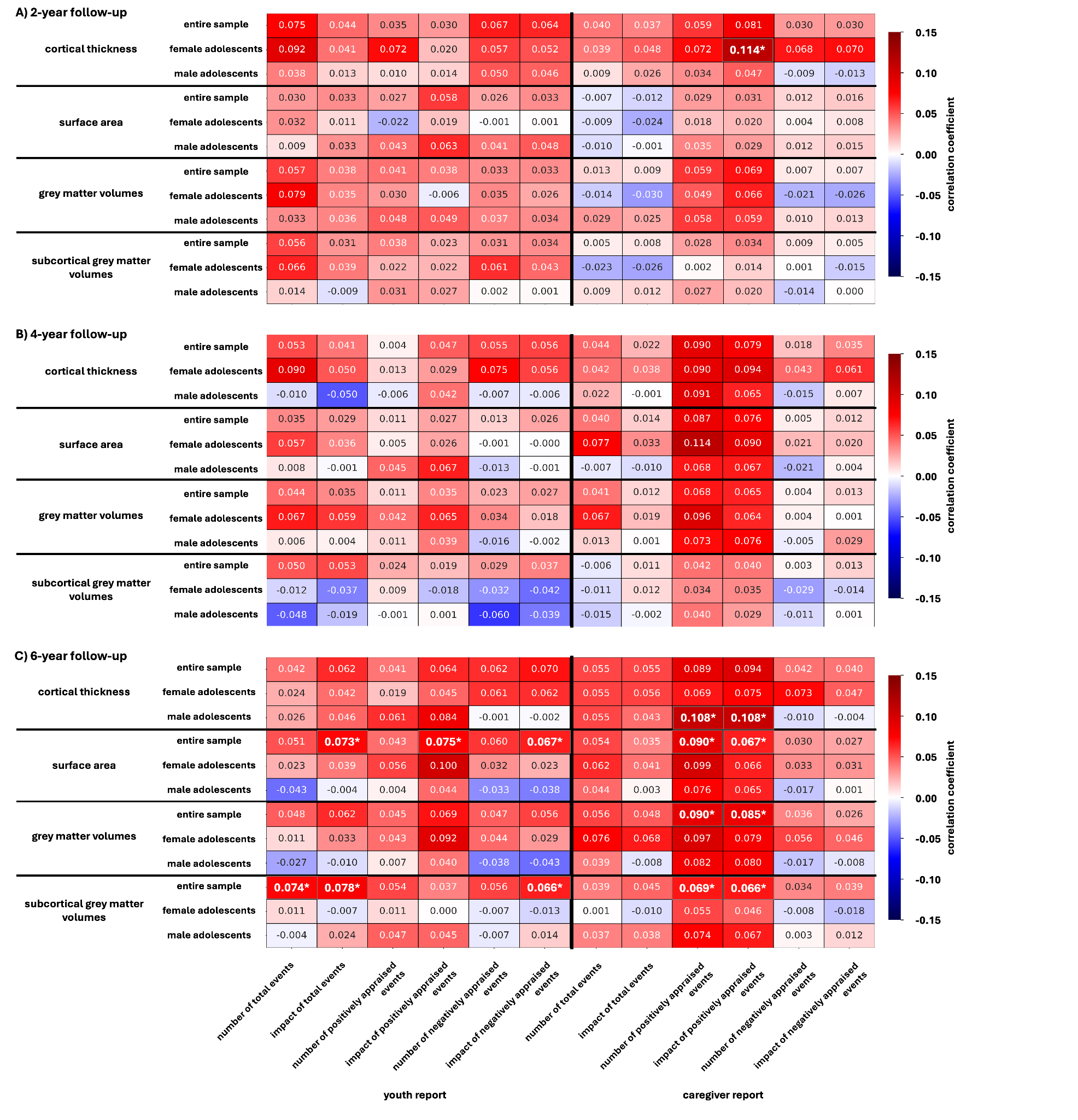
**Supplementary Figure 6. Predictive performance of neuroanatomical measures.** This heatmap illustrates the mean Pearson correlation coefficients capturing the association between predicted and actual data for models trained to predict the number, appraisal, or impact of stressful life events based on different neuroanatomical measurements (Cortical thickness, surface area, cortical and subcortical grey matter volumes) at the (A) 2-year follow-up, (B) 4-year follow-up, and (C) 6-year follow-up. The colors of the heatmap indicate the strength of the association depicted by the mean correlation coefficient. Negative associations are depicted in blue, positive associations are depicted in red. Measurements of cortical thickness, surface area and grey matter volumes are based on the Desikan-Killiany parcellation.

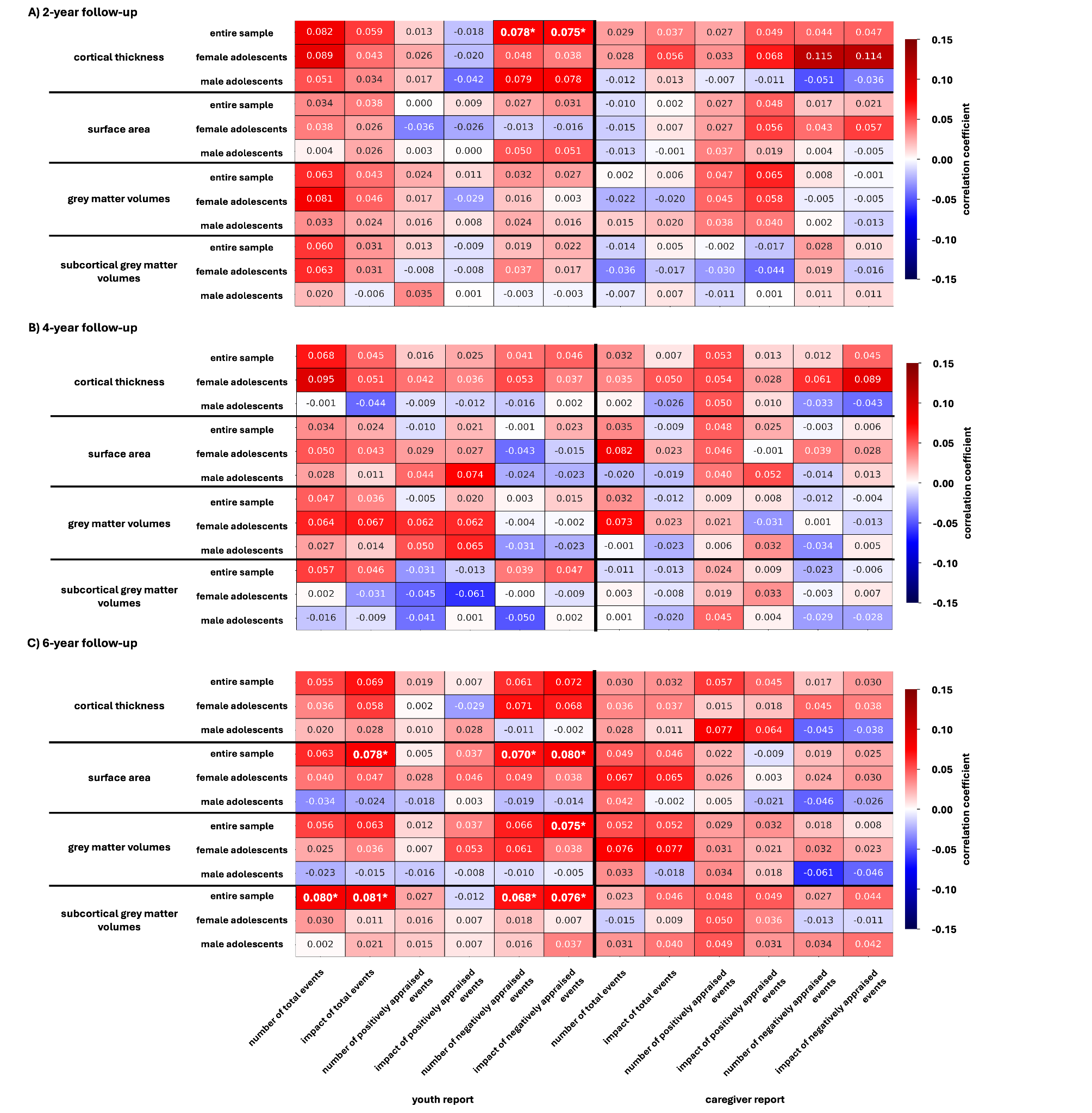

**Supplementary Figure 7. Predictive performance of neuroanatomical measures filtered for zero-values.** This heatmap illustrates the mean Pearson correlation coefficients capturing the association between predicted and actual data for models trained to predict the number, appraisal, or impact of stressful life events based on different neuroanatomical measurements (Cortical thickness, surface area, cortical and subcortical grey matter volumes) at the (A) 2-year follow-up, (B) 4-year follow-up, and (C) 6-year follow-up after filtering each individual target variable for zeros. The colors of the heatmap indicate the strength of the association depicted by the mean correlation coefficient. Negative associations are depicted in blue, positive associations are depicted in red. Measurements of cortical thickness, surface area and grey matter volumes are based on the Desikan-Killiany parcellation.
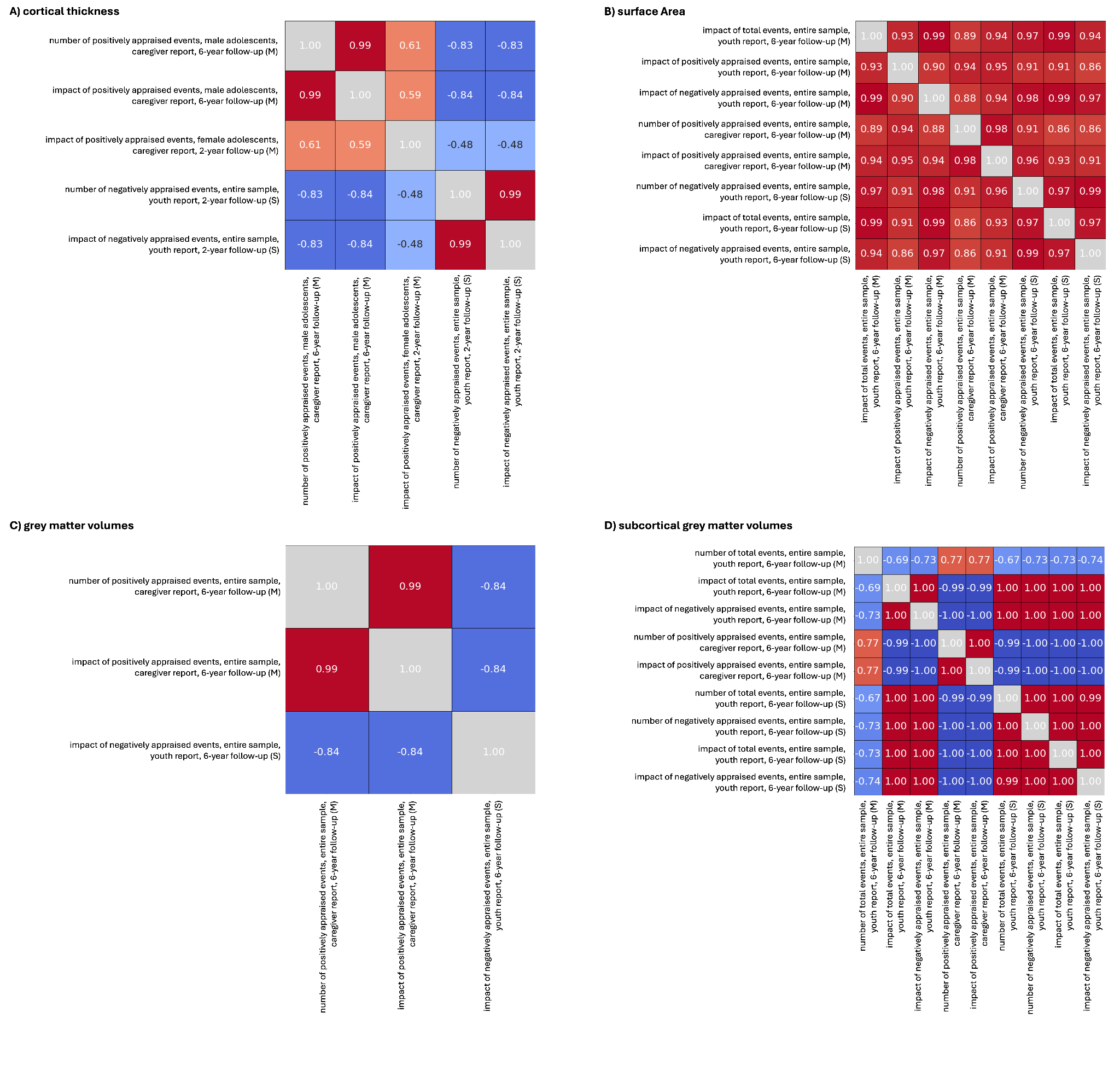

**Supplementary Figure 8. Correlations of feature weights.**

Correlations of feature weights of significant predictions for (A) Cortical thickness, (B) Surface Area, (C) Grey matter volumes, and (D) Subcortical grey matter volumes. All correlations are significant on a level of p < 0.01.

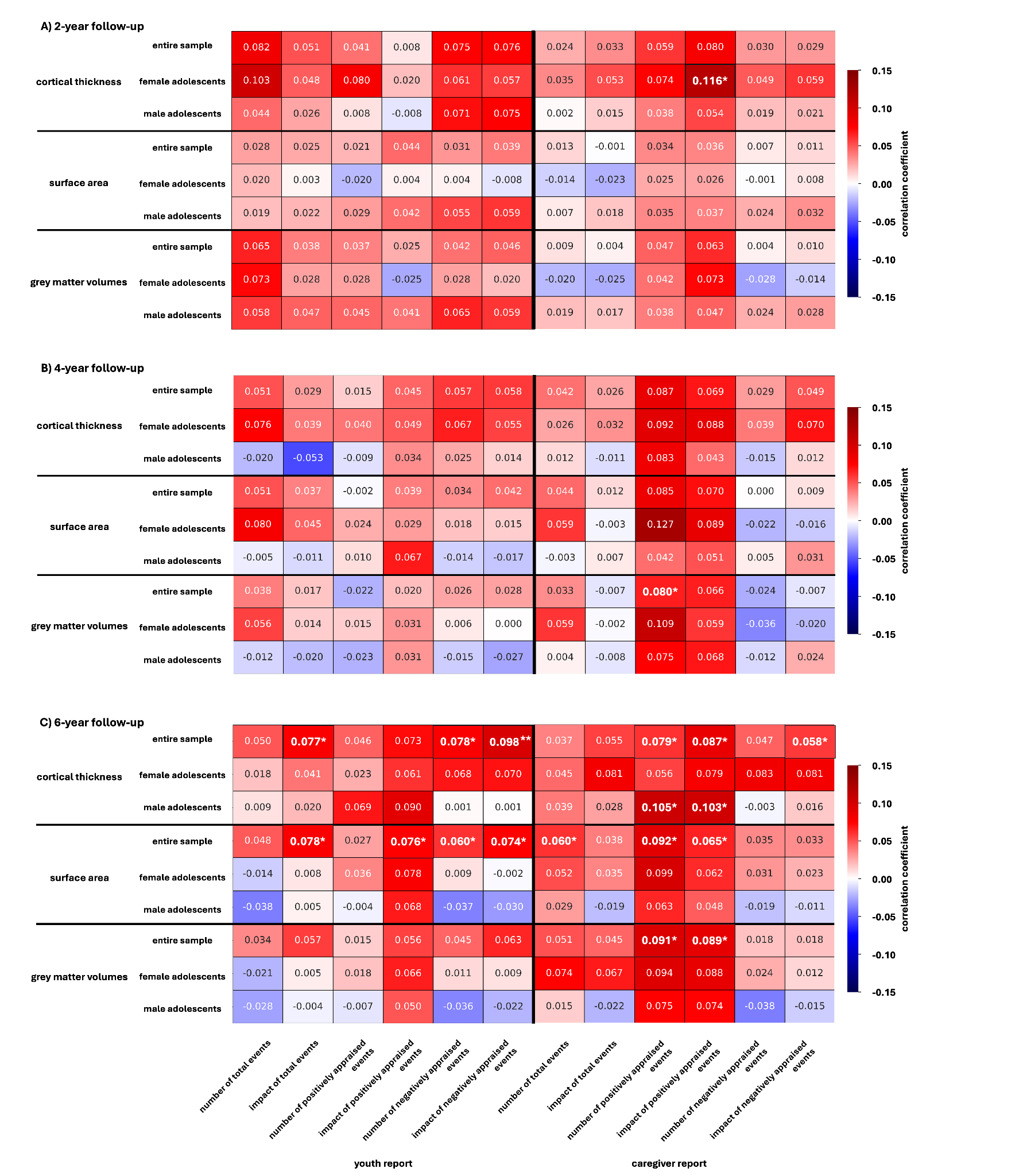
**Supplementary Figure 9. Predictive performance of neuroanatomical measures based on the Destrieux parcellation.** This heatmap illustrates the mean Pearson correlation coefficients capturing the association between predicted and actual data for models trained to predict the number, appraisal, or impact of stressful life events based on different neuroanatomical measurements (Cortical thickness, surface area, and cortical grey matter volumes) at the (A) 2-year follow-up, (B) 4-year follow-up, and (C) 6-year follow-up. The colors of the heatmap indicate the strength of the association depicted by the mean correlation coefficient. Negative associations are depicted in blue; positive associations are depicted in red. Significant predictions are depicted in bold.

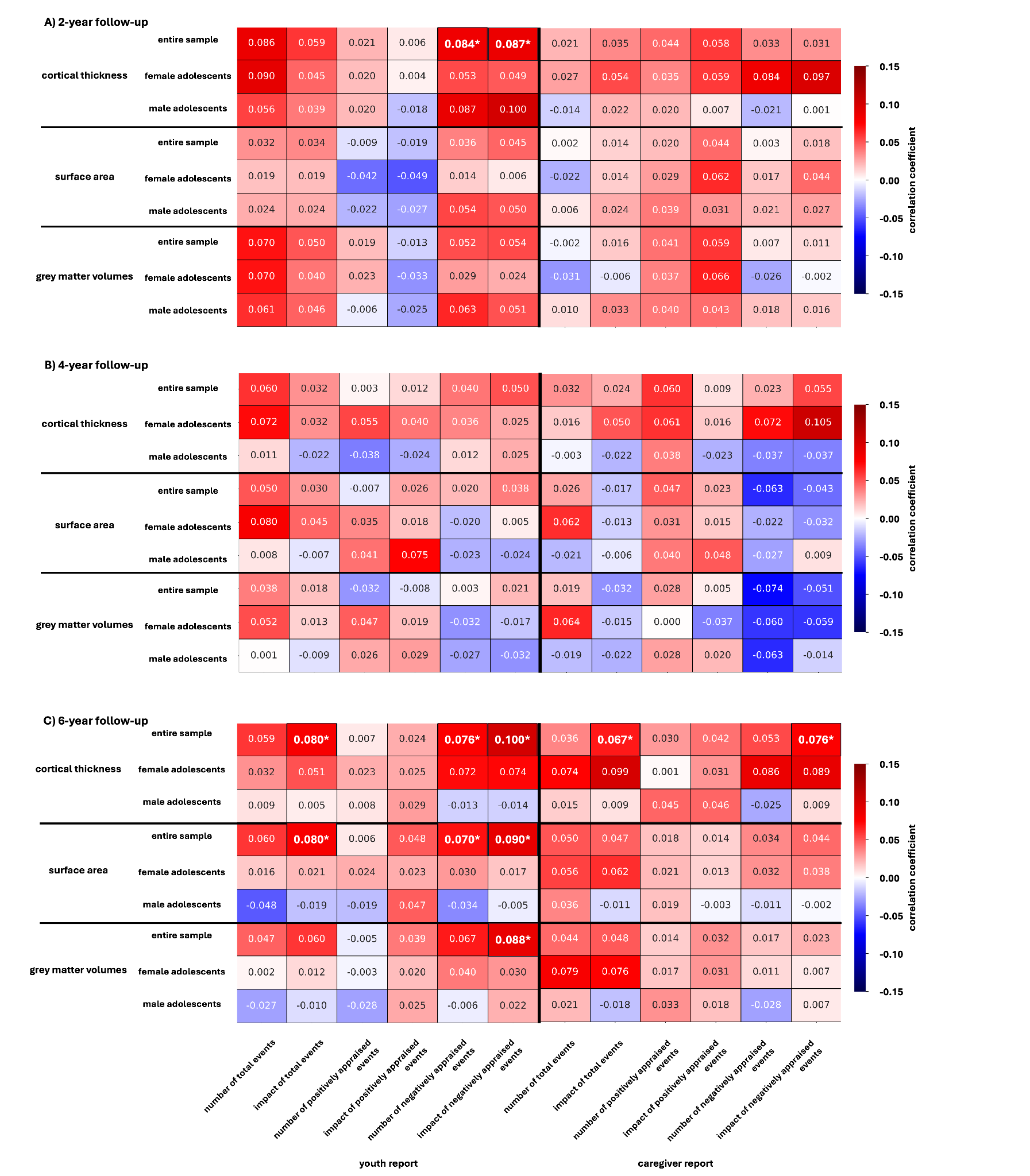
**Supplementary Figure 10. Predictive performance of neuroanatomical measures filtered for zero-values based on the Destrieux parcellation.** This heatmap illustrates the mean Pearson correlation coefficients capturing the association between predicted and actual data for models trained to predict the number, appraisal, or impact of stressful life events based on different neuroanatomical measurements (Cortical thickness, surface area, and cortical grey matter volumes) at the (A) 2-year follow-up, (B) 4-year follow-up, and (C) 6-year follow-up after filtering each individual target variable for zeros. The colors of the heatmap indicate the strength of the association depicted by the mean correlation coefficient. Negative associations are depicted in blue; positive associations are depicted in red. Significant predictions are depicted in bold.

**Supplementary Table 7. Correlations between SES and target variables**

Spearman rank correlations and p-values depicting the relationship between SES and number, perception, and impact of stressful life events for the (A) 2-year follow-up, (B) 4-year follow-up, and (C) 6-year follow-up time point for the youth report and caregiver report.

| **2-year follow-up** |  | **target variable** | **spearman correlation (ρ)** | **p-value** |
| --- | --- | --- | --- | --- |
|  | **youth report** | **number of total events** | -0.1982893 | p < 0.001* |
|  |  | **impact of total events** | -0.1444131 | p < 0.001* |
|  |  | **number of positively appraised events** | -0.0793764 | p < 0.001* |
|  |  | **impact of positively appraised events** | 0.03740037 | p = 0.00437 |
|  |  | **number of negatively appraised events** | -0.1777297 | p < 0.001* |
|  |  | **impact of negatively appraised events** | -0.1839963 | p < 0.001* |
|  | **caregiver report** | **number of total events** | -0.0602842 | p < 0.001* |
|  |  | **impact of total events** | -0.0642413 | p < 0.001* |
|  |  | **number of positively appraised events** | 0.09983607 | p < 0.001* |
|  |  | **impact of positively appraised events** | 0.10100879 | p < 0.001* |
|  |  | **number of negatively appraised events** | -0.1331706 | p < 0.001* |
|  |  | **impact of negatively appraised events** | -0.1385192 | p < 0.001* |
| **4-year follow-up** | **youth report** | **number of total events** | -0.0698132 | p < 0.001* |
|  |  | **impact of total events** | -0.0437235 | p = 0.00321 |
|  |  | **number of positively appraised events** | 0.01660357 | p = 0.26335 |
|  |  | **impact of positively appraised events** | 0.07631181 | p < 0.001* |
|  |  | **number of negatively appraised events** | -0.0588177 | p < 0.001* |
|  |  | **impact of negatively appraised events** | -0.0846036 | p < 0.001* |
|  | **caregiver report** | **number of total events** | -0.0161062 | p = 0.27792 |
|  |  | **impact of total events** | -0.0401103 | p = 0.00687 |
|  |  | **number of positively appraised events** | 0.14775611 | p < 0.001* |
|  |  | **impact of positively appraised events** | 0.14102281 | p < 0.001* |
|  |  | **number of negatively appraised events** | -0.1167136 | p < 0.001* |
|  |  | **impact of negatively appraised events** | -0.1328952 | p < 0.001* |
| **6-year follow-up** | **youth report** | **number of total events** | -0.037193 | p = 0.00636 |
|  |  | **impact of total events** | -0.0230563 | p = 0.09084 |
|  |  | **number of positively appraised events** | 0.05473539 | p < 0.001* |
|  |  | **impact of positively appraised events** | 0.09028058 | p < 0.001* |
|  |  | **number of negatively appraised events** | -0.0333513 | p = 0.01443 |
|  |  | **impact of negatively appraised events** | -0.061763 | p < 0.001* |
|  | **caregiver report** | **number of total events** | -0.0194611 | p = 0.15351 |
|  |  | **impact of total events** | -0.0517809 | p = 0.00014 |
|  |  | **number of positively appraised events** | 0.14055041 | p < 0.001* |
|  |  | **impact of positively appraised events** | 0.12348803 | p < 0.001* |
|  |  | **number of negatively appraised events** | -0.1061415 | p < 0.001* |
|  |  | **impact of negatively appraised events** | -0.1153217 | p < 0.001* |

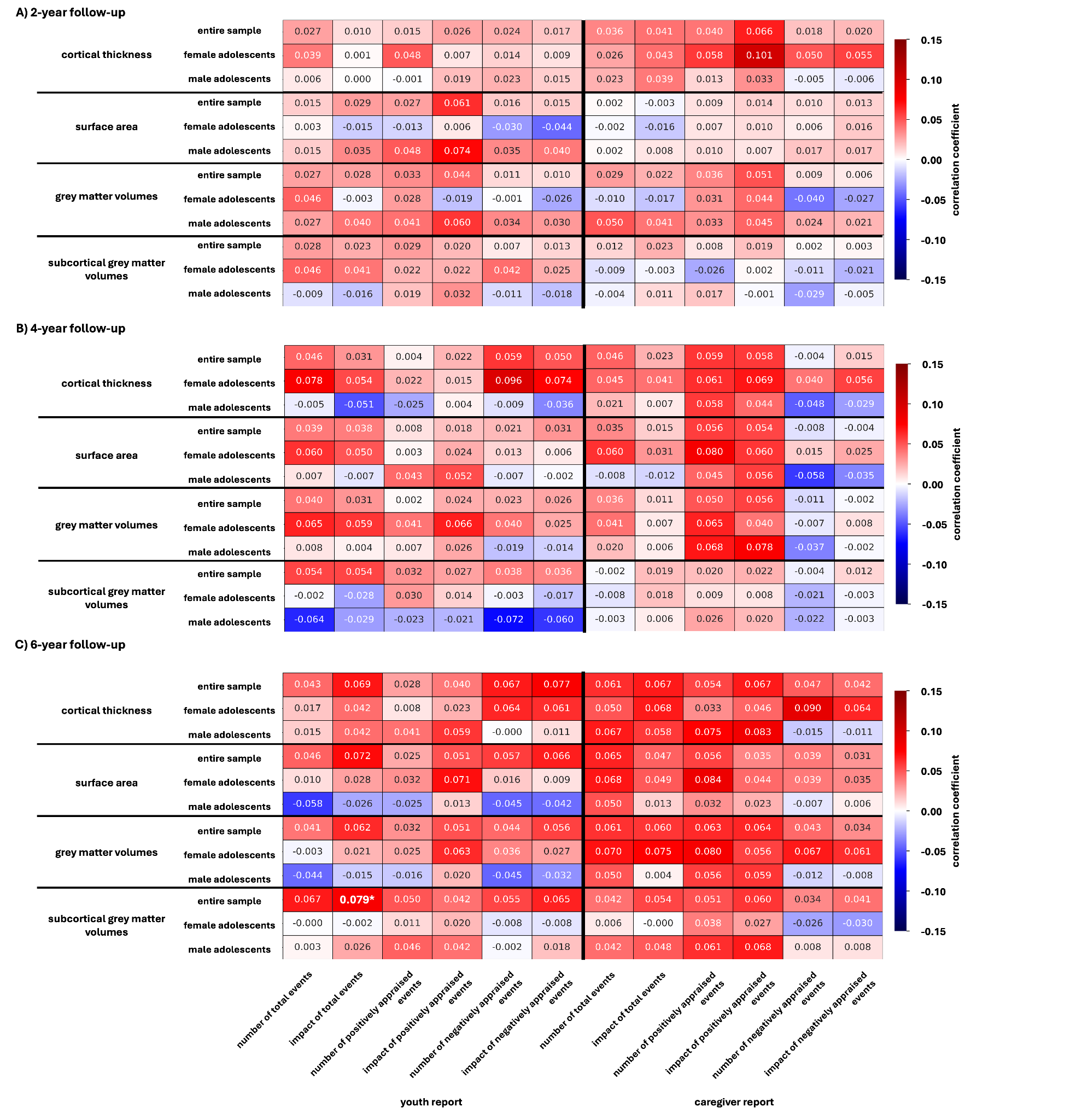
 **Supplementary Figure 11. Predictive performance of neuroanatomical measures controlled for SES.** This heatmap illustrates the mean Pearson correlation coefficients capturing the association between predicted and actual data for models trained to predict the number, appraisal, or impact of stressful life events based on different neuroanatomical measurements (Cortical thickness, surface area, cortical and subcortical grey matter volumes) at the (A) 2-year follow-up, (B) 4-year follow-up, and (C) 6-year follow-up while controlling for SES as a covariate. The colors of the heatmap indicate the strength of the association depicted by the mean correlation coefficient. Negative associations are depicted in blue, positive associations are depicted in red. Cortical measurements of cortical thickness, surface area and grey matter volumes are based on the Desikan-Killiany parcellation.

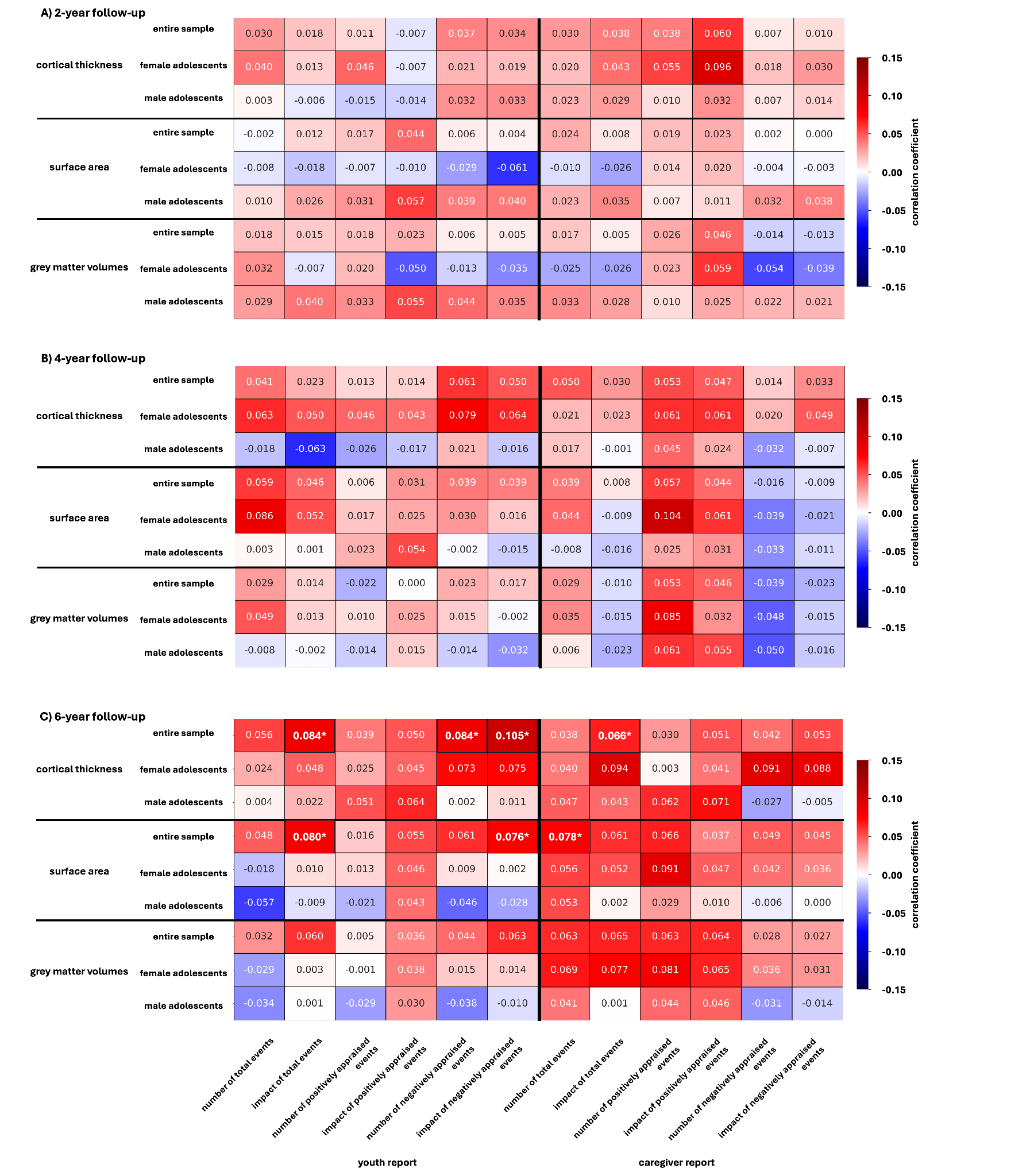
 **Supplementary Figure 12. Predictive performance of neuroanatomical measures controlled for SES.** This heatmap illustrates the mean Pearson correlation coefficients capturing the association between predicted and actual data for models trained to predict the number, appraisal, or impact of stressful life events based on different neuroanatomical measurements (Cortical thickness, surface area, and cortical grey matter volumes) at the (A) 2-year follow-up, (B) 4-year follow-up, and (C) 6-year follow-up while controlling for SES as a covariate. The colors of the heatmap indicate the strength of the association depicted by the mean correlation coefficient. Negative associations are depicted in blue, positive associations are depicted in red. Cortical measurements of cortical thickness, surface area and grey matter volumes are based on the Destrieux parcellation.
